# Computer Simulation and Mathematical Modeling of the Interactions Between Ecological and Sexual Selection to Reveal the Mechanism of Sympatric Speciation

**DOI:** 10.1101/2023.12.12.562603

**Authors:** John Lin, James Lin, Natalie Wassall

## Abstract

In modern evolutionary theory, the mechanism of sympatric speciation is still an unsolved mystery. Ecological niche specialization and the subsequent premating reproductive isolation (RI) are the putative initial steps of sympatric speciation, even though the exact mechanism of how the former leads to the latter remains unclear. This study aims to develop a simple, intuitive, and realistic model ecosystem that uses the fewest variables possible but is still comprehensive enough to uncover the fundamental mechanisms of sympatric speciation. Applying the concept of an adaptive fitness landscape, the study used an individual-based computer simulation to investigate how ecological selection and sexual selection may interact to produce sympatric premating RI. Mathematical models were derived to describe the nonlinear adaptive dynamics of the system. A user-friendly computer application was able to solve, numerically, all the trajectories, fixed points, and bifurcation points of the models and display the solutions as phase portraits. The findings reveal that under favorable conditions, fixed points may emerge in the system’s phase portraits to create stable mating-bias-allele polymorphisms and varying degrees of premating RI between niche ecotypes. Solving the nonlinear equations of the mathematical models establishes the precise parametric values required to produce such fixed points. It also specifies the conditions necessary for high-mating-bias mutant alleles to invade and produce, or enhance, premating RI. Restricting migration between niches impedes mutant invasion in parapatric and allopatric populations. Multi-locus mating traits may produce hybrids that prevent speciation. Simulation results of a gonochoric, multi-niche system demonstrate that linkage disequilibrium can emerge spontaneously among ecological, male-trait, and female-preference genotypes to produce premating RI. From these results, a five-stage mechanism of sympatric speciation is formulated: Initially, disruptive ecological selection creates selection pressure for high-mating-bias alleles to invade and rapidly establish premating RI. This then paves the way for the recruitment of late-stage mechanisms, such as adaptive coupling and post-zygotic isolation, to complete the speciation process. Applying nonlinear dynamics theories to model and analyze sympatric speciation has yielded novel findings that support and extend the results of prior studies. These new discoveries may help to overcome the theoretical objections to sympatric speciation and establish it as a prevalent mode of speciation in nature.

## I. Introduction

**M**ORE than 160 years after Darwin published *The Origin of Species*, the strife at the house of speciation is far from being settled. Allopatric speciation has emerged as the widely accepted ruler of the speciation process, with its close cousin, parapatric speciation, by its side, eager and ready to invoke the possibility of secondary contact to address any difficulties that allopatric speciation may encounter. Meanwhile, sympatric speciation remains the outlier whose legitimacy and validity are constantly being questioned. An annoyed Ernst Walter Mayr once called it “the Lernaean hydra which grew two new heads whenever one of its old heads was cut off” [1]. This is despite mounting theoretical and empirical evidence that sympatric speciation happened and was likely more prevalent than evolutionists believed [2-4]. The most compelling example is probably the rapid evolution and explosive speciation of sympatric cichlid fishes in African rift lakes and Central American crater lakes, which have baffled evolutionary biologists to this day [5-11].

According to the Bateson–Dobzhansky–Muller (BDM) model, allopatric speciation develops reproductive isolation (RI) as an incidental byproduct of genetic drift or genetic divergence after geographically isolated populations have adapted to their different environments [12, 13]. This is a process that usually takes a long time [14, 15]. In contrast, sympatric speciation dictates that speciation must occur in a common arena without geographical barriers. As a consequence, the resulting gene flows among sympatric populations are liable to break up any favorable, locally adaptive gene combinations through recombination and homogenize the genotypes of the populations [16-18]. This can make speciation challenging, if not impossible. For this reason, sympatric speciation, when it occurs, tends to be faster and develop stronger mating biases than allopatric speciation [19-21], presumably because the incipient species need to actively counter the homogenizing effect of gene flow from their cohabitating sister species. Consensus among theoreticians to date suggests that the most likely mechanism of sympatric speciation involves the following steps: (1) Disruptive ecological selection causes the emergence of separate, distinct sympatric populations of ecotypes that are most adaptive to their different niches. (2) Assortative mating and reproductive isolation develop between locally adapted populations of ecotypes, reduce gene flow between them, and help to complete the speciation process [3]. Currently, the most controversial debates center around Step 2, that is, how reproductive isolation develops among locally adapted niche populations in sympatry [2, 22]. Many empirical studies have suggested that premating reproductive isolation plays a prominent role in incipient sympatric speciation [15, 23-29]. On the theoretical front, computer modeling and simulation have shown that under certain favorable, albeit stringent, conditions [30], frequency-dependent disruptive selection—caused by unfit hybrids or because of competition—can cause marker alleles that control mating biases to assort spontaneously among ecotypes to achieve reproductive isolation [31-33].

In the fossil record, speciation events tended to happen in short bursts followed by long periods of stasis as described by the phenomenon of punctuated equilibrium [34, 35]. This makes it difficult to go back in time, either through paleontological records or genetic phylogeny studies, to uncover the exact mechanisms or favorable conditions that could have caused the speciation events. Still, even though speciation time may be short in evolutionary time, it remains too long for evolutionary biologists to observe and study in human time. For these reasons, computer modeling and simulations have become an attractive alternative to decipher the mechanisms and conditions necessary for sympatric speciation to occur.

Over the past two decades, the growing accessibility and processing power of modern computers have spurred the development and testing of many theoretical models exploring the evolution of premating RI in sympatric speciation [36]. The most promising approach appears to be those based on adaptive population dynamics [31, 32, 37-40], which, in computer simulations, have been shown to produce spontaneous mating-bias-allele assortment between different sympatric niche-ecotypes under disruptive selection. However, many evolutionary biologists have criticized such models as inadequately simplistic, focusing on particular aspects of the problem while neglecting others, and making unrealistic assumptions that do not represent real-world systems [3, 39, 41-45]. Moreover, speciation driven by selection is inherently complex. Existing theoretical models tend to utilize complicated mathematical formulations involving differential equations with many arbitrary variables and initial conditions, which lack analytical solutions [40]. This makes them difficult to decipher without a high level of mathematical competency [30]. The large number of variables and the high dimensionality of the solution space also hinder intuitive understanding and extraction of general principles that help to discern the forest for the trees [46]. The particular nature of the models makes it difficult to compare and generalize results across different studies to gain robust insights [30]. Nonetheless, one result that has emerged reliably from all the research studies is that the development of premating RI in sympatric speciation involves the intricate interplay between ecological selection and sexual selection [22, 30, 47], despite our lack of a clear understanding of how these two selection forces interact and the precise parameters required to induce mating-bias-allele assortment and RI [43, 48, 49].

To address these shortcomings, our study aims to develop a simple, intuitive, and realistic model ecosystem that uses the minimum number of variables possible yet is still comprehensive enough to allow us to uncover the fundamental mechanisms of sympatric speciation. Our study follows the “two-allele model” of sympatric speciation [3, 16], wherein ecological alleles and mating-bias alleles are located on separate, unlinked genetic loci. By applying the theories of adaptive population dynamics, we investigated how the interplay between ecological selection and sexual selection can lead to premating RI in sympatric speciation. Our goal is to solve the “hard process” of sympatric speciation [2] by assuming the absence of geographical barriers that can impede mating encounters or gene flow. The expectation is that once we can explain hard sympatric speciation, other “easy processes” of sympatric speciation, such as those due to magic trait [12, 50, 51], secondary contact [2, 52], or limited gene flow [53-56], become readily solvable.

To accomplish our goal, we first used an individual-based computer simulation to investigate how ecological selection and sexual selection may interact to produce premating RI in a simplified 2-niche sympatric ecosystem. Individuals in the simulation have genotypes that consist of three loci for ecological alleles and a locus for two mating-bias alleles. These individuals were allowed to encounter one another by random chance and reproduce offspring based on the compatibility of their mating-bias alleles (or mating based on matching rules [51]). The evolution of the population’s genotype composition was visualized on a three-dimensional genotype/phenotype fitness landscape, by showing individual genotypes as dots in the 3-D plot. In such a fitness landscape, an individual’s ecological phenotype is determined by epistatic and pleiotropic interactions among the individual’s ecological alleles.

The use of the adaptive fitness-landscape metaphor in evolution research has a long and established history [12, 57-61]. Our computer simulation showed that when individuals of our panmictic population in such a fitness landscape were allowed to reproduce offspring genotypes by random assortment of their parental alleles, the entire population’s genotypes (visualized as clouds of dots in the 3-D plot) exhibited swarming behavior [62, 63]. This behavior was driven by ecological selection and resulted in the aggregation of genotypes into distinct ecotype groups at different niche locations. However, the emergence of such distinct adaptive ecotype groups in the fitness landscape was not always guaranteed. We identified a phenomenon that we termed “incumbent selection,” whereby a larger group of individuals in one niche could use their numerical advantage to eliminate a smaller group in another niche. We also demonstrated that by manipulating parametric variables within our 2-niche model system—such as hybrid viability, premating trait bias, the cost of assortative mating, offspring fecundity, and the maximum niche carrying capacity—we could observe spontaneous assortment of mating-bias alleles in the two groups of niche ecotypes. This assortment continued until a stable polymorphic equilibrium was attained, leading to varying degrees of premating RI between the two niche groups. These findings align with the results of earlier researchers [30].

To determine the precise parametric values that can produce premating RI, we derived a simple yet generalizable mathematical model to describe the population dynamics in our 2-niche ecosystem. The mathematical model consists of sets of difference equations. A user-friendly computer application was developed to solve the difference equations by numerical methods so that their phase-portrait solutions could be plotted and visualized swiftly across all possible ranges of parametric values. This gives us a bird’s-eye view of the entire solution landscape and enables us to examine all the system’s vector-field trajectories, fixed points, and bifurcation points to perform a comprehensive analysis of its dynamic behavior. The model uses normalized parametric variables which makes our computer-aided analysis generalizable and applicable to a wide range of different scenarios. The model’s accuracy and validity were confirmed by the close match between its computed numerical solutions and the results from the individual-based computer simulation.

Mathematical modeling of our idealized 2-niche sympatric ecosystem proved that stable premating RI between different niche ecotypes cannot be achieved unless the carrying capacities of the niches are limited. Moreover, one of the following two conditions must be met: (1) the adaptive ecotypes reproduce sufficient offspring to completely fill the carrying capacities of their respective niches, or (2) ecotypes residing in a smaller, unsaturated niche can proportionately increase their offspring numbers based on their increased fitness to escape incumbent selection from ecotypes in larger niches. Consistent with the findings in existing literature, analysis of our model showed that when the assortative mating cost is low, the effects of ecological selection and sexual selection are complementary in producing premating RI [64]. The emergence of premating RI is most likely when there is strong ecological selection against hybrids, a high mating bias, a low cost of assortative mating, and approximately equal population sizes in the two niches [30, 36]. In such favorable circumstances, we could observe the appearance of fixed points in the phase-portrait solutions of the system. These fixed points represent stable polymorphisms of mating-bias alleles that create various degrees of premating RI between the ecotypes [65].

The evolution of a sympatric species into separate sister species by ecological specialization likely requires the invasion or co-optation of high-mating-bias alleles into the initially panmictic population to establish strong premating RI. Analysis of our model’s dynamic behaviors has revealed that when the origin of its phase portrait falls within the basin of attraction of a fixed point, the system can create positive selection pressure for the associated high-mating-bias mutant allele to invade a population that has only a single type of mating-bias allele. Subsequently, the invading mutant can achieve fixed-point mating-bias-allele polymorphism with the preexisting mating-bias allele and produce premating RI.

High-mating-bias mutant alleles can also invade a system with low-mating-bias alleles to incrementally increase its premating RI. An individual-based computer simulation was used to demonstrate how a mutant meta-allele producing a high mating bias can invade a population of low-mating-bias meta-alleles to increase the premating RI between the niches. Similarly, analyses of a 2-niche, 3-mating-bias-allele model showed that, under favorable conditions, a third high-mating-bias mutant allele can invade a 2-mating-bias-allele system and increase its premating RI. In general, it is easier for a high-mating-bias meta-allele or a third high-mating-bias allele to invade a system that already has a low degree of premating RI than a system without any fixed-point premating RI. The existence of such “invasion thresholds” might be nature’s way to prevent excessive speciation [45].

We created contact barriers between the niches in our models to investigate how the presence of geographical barriers in parapatry and allopatry could affect the development of premating RI. The findings revealed that restricting migration between niches impedes invasion by high-mating-bias alleles.

Computer analysis of a model with multi-locus mating-bias genotypes revealed that, just as the presence of viable ecological hybrids weakens ecological selection, the presence of mating-bias hybrids weakens sexual selection and hinders the emergence of premating RI and speciation. However, this effect could be mitigated by disruptive sexual selections that decrease the fitness of mating-bias hybrids.

An individual-based computer simulation was developed to explore a gonochoric “two-allele model” of sympatric speciation [3, 16] in a multi-niche ecoscape. The results demonstrated that under favorable parametric conditions, initially unlinked, randomly distributed male-trait and female-preference genotypes would cluster in different groups of niche ecotypes to establish premating RI between the different niche groups. This spontaneous occurrence of linkage disequilibrium among male-trait alleles, female-preference alleles, and ecological alleles to produce premating RI and reduce hybrid loss among niche ecotypes appears to be an emergence property of sympatric systems undergoing disruptive selection. Despite its increased complexity, the system exhibited analogous population and invasion dynamics as those in its simpler, unisex, 2-mating-bias-allele, 2-niche counterparts, suggesting the existence of fundamental, discoverable general operating principles underlying all forms of sympatric speciation.

Lastly, we used the results of the computer simulation and mathematical modeling to synthesize a five-stage process of sympatric speciation driven by selection. In the initial stages, a sympatric species occupying a single niche is driven by intraspecies sexual selection to eliminate the less common, high-mating-bias alleles in the population. The situation changes when an environmental upheaval creates disruptive ecological selection that favors distinct adaptive niche ecotypes and selects against their less-fit hybrid offspring. High-mating-bias alleles, previously selected against, are now positively selected to invade and establish premating RI between the different niche ecotypes. The driver of this positive selection pressure comes from newly created niche resources that are the result of unfit hybrid loss or underutilized ecological niches. Even though this initial premating RI, mediated by reproductive character displacement and the system’s population dynamics, can evolve rapidly, it tends to be weak and reversible. Nevertheless, it facilitates the evolution of other slower, but more permanent, late-stage mechanisms—such as intrinsic post-zygotic barriers through adaptive coupling [66, 67] or the BDM model of incompatibility—to strengthen and consolidate the RI between niche populations. After speciation is completed, intraspecies sexual selection again dominates to eliminate incompatible high-mating-bias alleles in the now reproductively isolated sympatric species.

The theory of evolution forms the bedrock of modern biology, and understanding the process of speciation lies at the heart of evolutionary theory. Allopatric speciation facilitated by geographical barriers, originally proposed by Darwin, makes intuitive sense and has received wide empirical support. Meanwhile, our understanding of sympatric speciation is still lacking. The major obstacle to the wide acceptance of sympatric speciation appears to be theoretical [16, 31, 68], in that there is still not a satisfactory and realistic mechanism to adequately explain how speciation could have happened in the presence of the homogenizing effect of gene flow.

Our study has proposed a simple, yet intuitive and generalizable model of sympatric ecosystems that can capture the underlying principles of sympatric speciation. Employing the concept of an adaptive fitness landscape, our individual-based computer simulation elucidates the adaptive dynamics of the model systems and confirms the accuracy of their mathematical models. These mathematical models can serve as a foundational framework for developing additional analytical theorems and solutions. It turns out that sympatric speciation models are best analyzed as nonlinear dynamic systems. Moreover, we can completely solve the behaviors of such systems using the theories of nonlinear dynamics, aided by the power of modern computers and numerical analysis techniques. Displaying the computed results graphically as phase portraits helps us gain a comprehensive view of the entire solution landscape and better understand the systems’ dynamic behaviors. This methodology allows us to precisely determine the normalized parametric values that are required to produce reproductive character displacement and premating RI, as well as elucidate the fundamental principles governing the interactions between ecological selection and sexual selection in sympatric speciation. The ability to visualize our model’s dynamic behavior offers an intuitive understanding of how disruptive ecological selection can generate new niche resources and create positive selection pressures for suitable mutant mating-bias alleles to invade and create, or enhance, premating RI. Our findings are consistent with previous studies but provide a more generalizable and in-depth understanding of the conditions under which sympatric speciation can occur. Lastly, based on the results of our study, we have proposed a mechanism of sympatric speciation that, we hope, is intuitive and realistic enough to address many of the theoretical objections to sympatric speciation and help to unravel its mystery.

The novel approaches of our study have revealed new findings and a mechanism of sympatric speciation that is actively driven by selection and can occur rapidly. This contrasts with allopatric speciation which tends to occur slowly and arise as a chance byproduct of genetic drift and adaptations to different environments. Considering the more abundant opportunities for sympatric speciation to occur, our study results permit us to offer arguments suggesting that sympatric speciation might be more prevalent than allopatric speciation as the primary mode of speciation in nature [69].

## II. Methodology

A MATLAB (version R2021a) software program was used to develop an individual-based computer simulation. Utilizing the App Designer tool in MATLAB, a user-friendly graphical user interface (GUI) application was designed to solve, using numerical methods, the sets of nonlinear difference equations that describe the mathematical models derived in the study and display the results visually.

## III. Individual-based Computer Simulation

Fig 1 shows the flowchart of a computer program that was developed to simulate the life cycle of a sympatric population. Each individual in the population has a genotype that consists of three loci for ecological alleles (*e*_1_, *e*_2_, *e*_3_) and a locus for mating-bias alleles (*s*), as shown in Fig 2. To facilitate conceptual visualization, we specified that there are 60 alleles in each ecological locus, numbered from 1 to 60. At the mating-bias allele locus, there are two alleles, *X* and *Y*, which interact according to the matching compatibility table shown in Fig 3.

**Fig 1.**
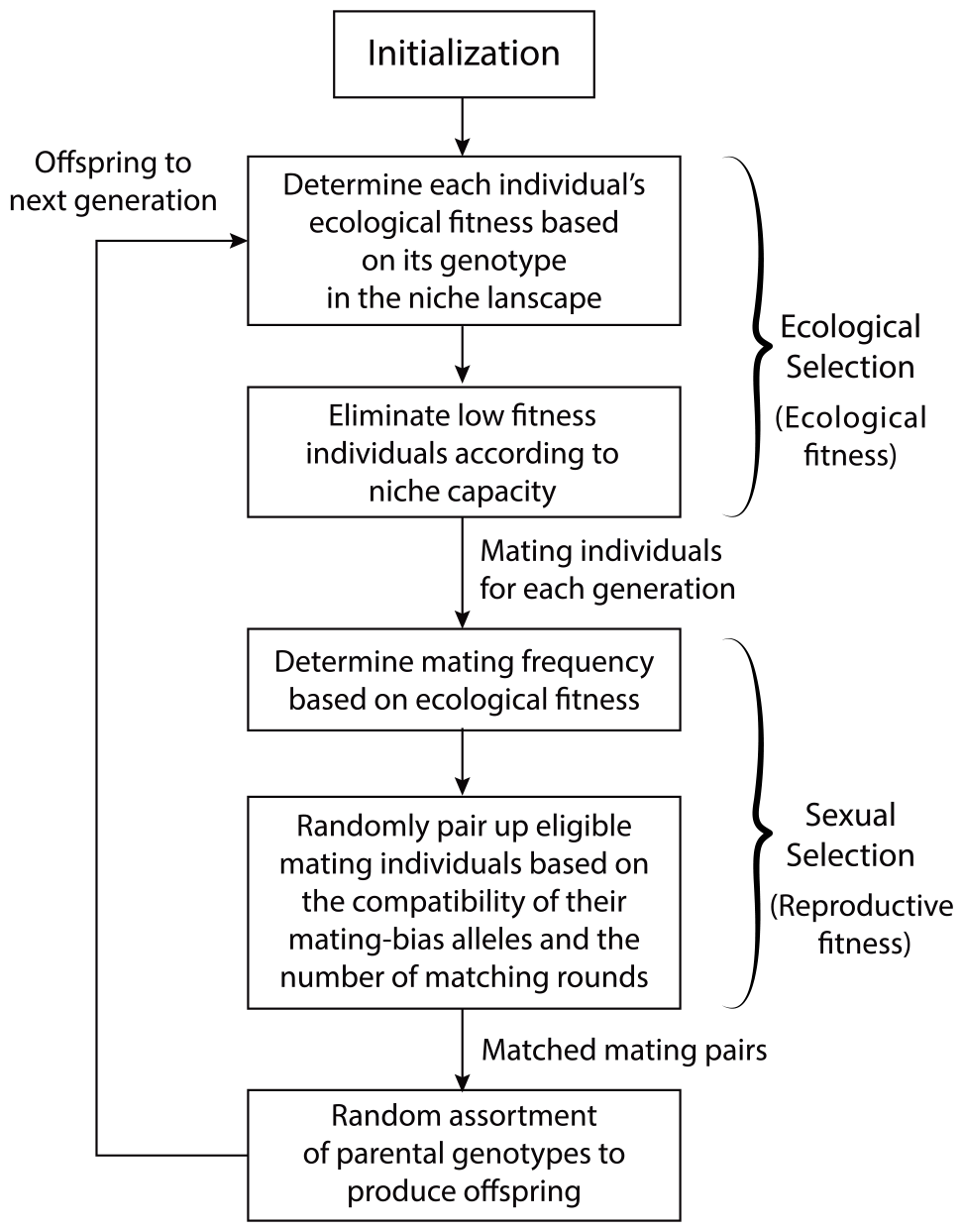
A flowchart showing the life cycle of a sympatric population in an individual-based computer simulation.

**Fig 2.**
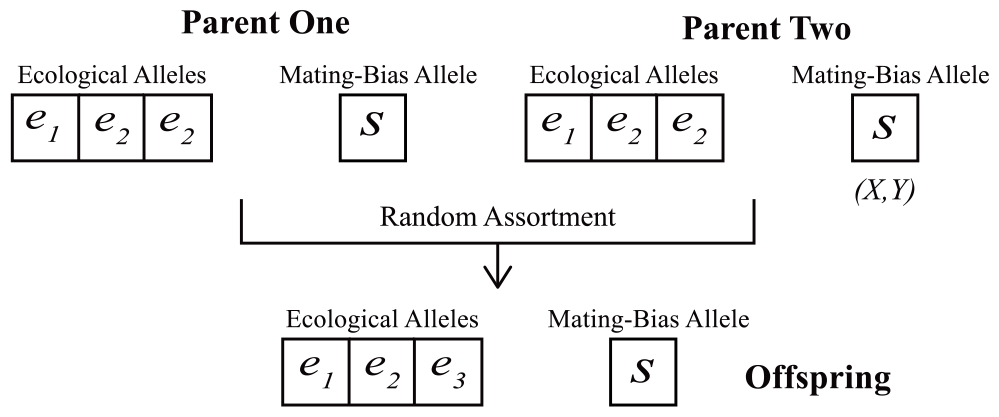
Genotype of an individual used in an individual-based computer simulation. Alleles of parental genotypes combined by random assortment to produce offspring genotypes.

**Fig 3.**
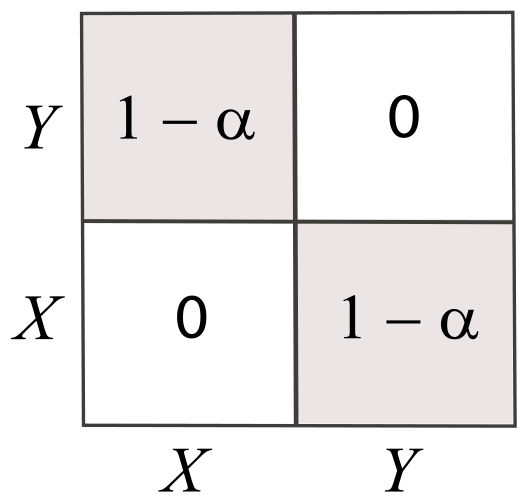
A matching compatibility table for mating-bias alleles. When two individuals possess the same mating-bias alleles, the mating barrier between them is zero, and they are perfectly compatible. When they possess different mating-bias alleles, their compatibility is determined by variable *α*.

When two individuals with the same mating-bias allele (*X* or *Y*) meet, the mating barrier is zero, and they are perfectly compatible. They then become a matched mating pair and are taken out of the eligible mating pool. When the two individuals possess different mating-bias alleles, however, the probability of them becoming a matched mating pair is determined by a probability ratio *α*, the value of which ranges from 0 to 1. The higher the value of *α*, the higher the chance of successful mating. All individual ecological genotypes can be plotted as points on a three-dimensional ecoscape in Fig 4a, where the *x, y*, and *z* axes represent the allele types at each ecological locus.

**Fig 4a.**
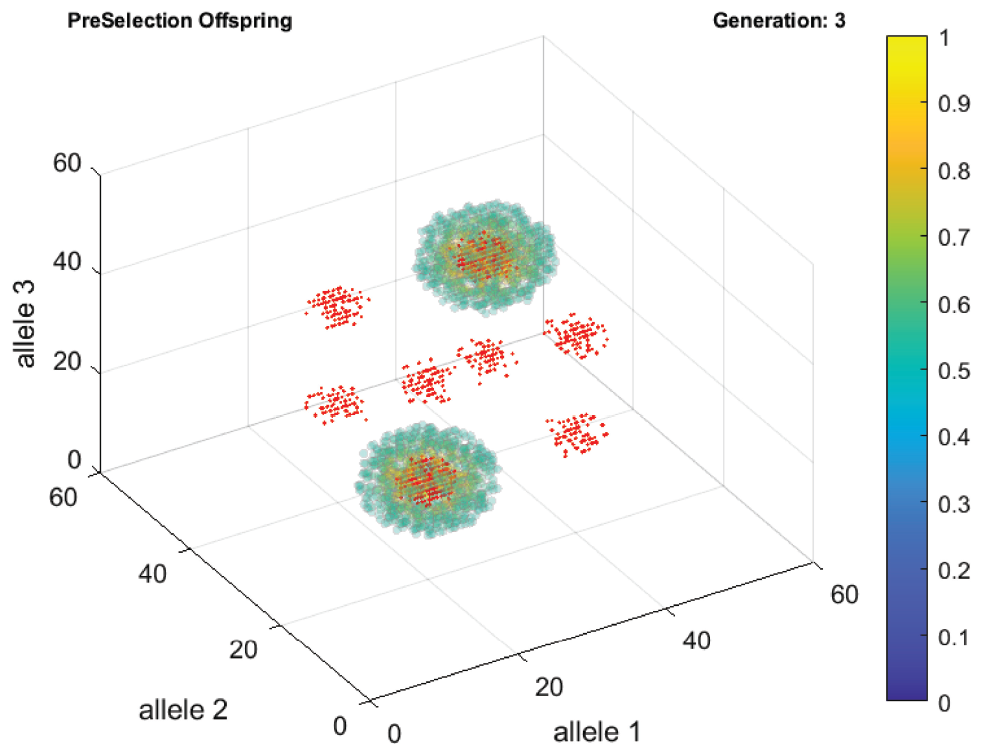
A computer screenshot showing a three-dimensional genotype-to-phenotype map. The *x, y*, and *z* axes represent the three ecological allele types, *e*_1_, *e*_2_, and *e*_3_, numbered from 1 to 60. Individual ecological genotypes are shown as dots. Ecological niches are shown as shaded ellipsoid volumes. The side colorbar shows the probability density in the ellipsoid volumes.

Fig 4b shows a central histogram that is a two-dimensional compression of the three-dimensional plot in Fig 4a, and the side histograms display the frequency distribution of the alleles on each axis. The locations of the niches for a 2-niche ecosystem are shown as shaded ellipsoids in Fig 4a. Each ellipsoid represents a 3-D volume of normal probability density distribution, defined by the means and variances of three separate normal probability density curves along the *x, y*, and *z* axes. Where an individual genotype lands in an ellipsoid volume determines how effectively the individual can extract food resources from the niche, which is a measure of its fitness in the niche. Genotypes that are near the center of an ellipsoid volume possess a greater ability to extract resources from the niche than genotypes at the periphery. The carrying capacity of each niche limits the maximum number of individuals it can support based on its finite food resources.

**Fig 4b.**
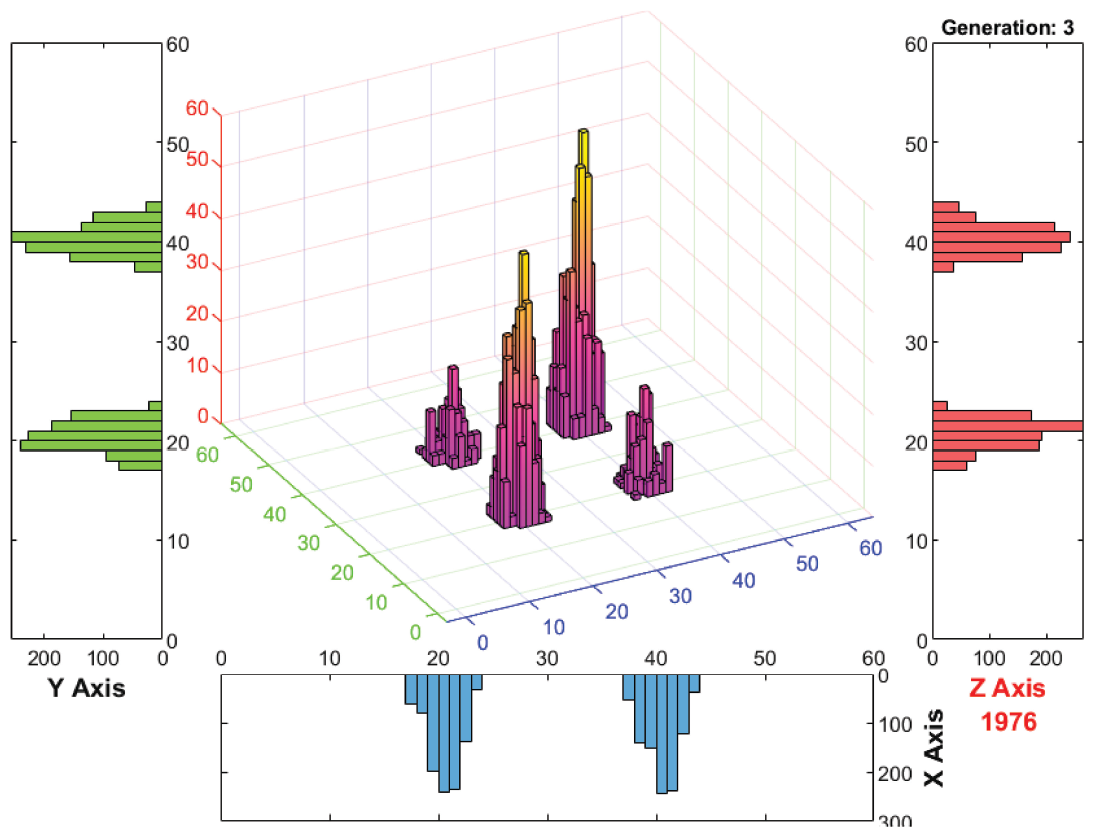
A computer screenshot of histograms that show the genotype and allele frequencies in Fig 4a. The center histogram is a projection of the three-dimensional distribution of individual genotypes in Fig 4a on a two-dimensional *x, y* plane. The side histograms show the frequency distributions of the population’s constituent alleles, *e*_1_, *e*_2_, and *e*_3_, on the *x, y*, and *z* axes.

In the ecological selection phase of the flowchart in Fig 1, individuals acquire differential ecological fitness based on how effectively their inherited genotypes can extract food resources from niches in the ecoscape. Individuals whose genotypes cannot allow them to attain a minimal amount of fitness, such as hybrids in the 2-niche ecosystem in Fig 4a, are eliminated. In contrast, individuals whose genotypes are close to the center of a high-carrying-capacity niche are able to acquire more ecological fitness. This translates into higher reproductive fitness by using a multiplier formula that awards high ecological fitness individuals with a higher mating frequency.

In the sexual selection phase of the flowchart, all the individuals that survive ecological selection are collected in a mating pool. They are allowed to encounter each other randomly in matching rounds and pair up to become successful mating pairs based on the compatibility of their mating-bias alleles. After a matching round, the successfully matched individuals are taken out of the mating pool, and the unmatched individuals go on to the next matching round. The exception is for the high-fitness individuals, who can duplicate themselves according to their awarded mating frequencies to increase their chance and number of successful matches in matching rounds, thereby reproducing more offspring. At the end of a specified number of matching rounds *n*, all the unmatched individuals are eliminated. After that, all the successfully matched mating pairs undergo random assortment of their alleles at the ecological and mating-bias loci to produce offspring for the next generation. The variable *off* specifies the number of offspring each matched pair of parents can produce.

As the computer simulation runs through *i* mating generations, dynamic changes in the population’s genotype composition after each generation are displayed visually. Fig 4a and 4b show an example of the population dynamics of a simulation after three generations. The simulation commenced with 1000 individuals uniformly distributed throughout the genoscape in Fig 4a. By the third generation, all the surviving individual genotypes had moved into the two shaded niche ellipsoids. The figures show the offspring genotypes and allele frequencies produced after the third generation. Because the two niches were positioned diagonally in Fig 4a, the offspring genotypes produced by individuals in the two niches mapped out the eight corners of a cube. As these offspring went through ecological selection in the fourth generation, all the offspring genotypes outside of the niches were eliminated.

The graph in Fig 4a represents a genotype-to-phenotype map. Interactions among alleles in an individual’s genotype produce phenotypes. If those phenotypes are advantageous in a niche and allow the individual to gain ecological fitness, the individual can reproduce more offspring carrying the same alleles and increase their allele frequencies in the gene pool. This, in turn, makes the emergence of the same genotype more likely in the population. So, in effect, the genotypes in Fig 4a, represented as dots, exhibit swarm behavior [62, 63]. The genotypes are always exploring the three-dimensional genotype/phenotype fitness landscape by creating variants with combinations of their alleles in each generation, and once they find a resourceful niche, they will draw the entire population there for subsistence.

The computer simulation also showed that, with the passing of generations, only the fittest individuals in a niche survive, i.e., those with genotypes near the center of a niche in Fig 4a. Because all those individuals have essentially the same ability to extract the niche’s resources, there is no relative fitness advantage, and they all survive with a bare-minimum viable amount of fitness as the population reaches the niche’s carrying capacity. If the population in a niche is small—because the starting population in the niche is small and/or because the matched pairs of parents in the niche are not able to reproduce enough offspring to saturate the carrying capacity of the niche—then the smaller population in the niche runs the risk of being eliminated by a larger population in another niche because of the effect of random mating and recombination. We term such a phenomenon “incumbent selection.” Incumbent selection occurs when a larger population is able to use its numerical advantage to suppress or eliminate a smaller population, such as, in our example, through gene flow and recombination. This can occur even in cases when the individual fitness in the larger population is lower than the individual fitness in the smaller population. In our computer simulation, a larger, established group of genotypes in a saturated niche, where individuals only possess minimal survivable fitness, can suppress or “cannibalize” a smaller genotype group in a mostly empty niche. This occurs even when the individuals in the smaller group are fitter and more fertile because more resources are available for a smaller number of individuals to extract.

The effect of incumbent selection can be mitigated by limiting the niche carrying capacity of the larger group or by increasing the number of offspring produced by the smaller group. Once the smaller group can achieve a large enough population size to counter the suppressive effect of incumbent selection from the larger group, the smaller group can then grow rapidly in population size to reach its niche’s carrying capacity. We can define a fecundity index *F* as the product of an individual’s mating frequency (*freq*) and the number of offspring that it can produce (*off*): *F* = *freq* × *off*. Given that an individual’s mating frequency is proportional to its ecological fitness, *F* is a measure of the gain in reproductive fitness as an individual’s ecological fitness increases. If the average value of *F* in a smaller population in an incompletely filled niche is below a certain threshold *F*_*min*_, the smaller population is liable to be eliminated by a larger population through incumbent selection. Conversely, if the average value of *F* is above a certain threshold *F*_*max*_, the smaller group can free itself from the suppressive effect of incumbent selection and rapidly expand its population size to reach its niche’s carrying capacity. Any values of *F* between *F*_*min*_ and *F*_*max*_ maintain a stable equilibrium state where a proportionally small but survivable population can exist in its niche.

## IV. Mathematical Model of Two Mating-Bias Alleles Without Viable Hybrids

A mathematical model to illustrate the nonlinear dynamic behavior of a 2-mating-bias-allele, 2-ecological-niche sympatric population is shown in Fig 5. Assume that there are two niches, *A* and *B*, in a 2-niche ecoscape. Suppose all the unmated individuals in the model meet randomly to find mates, and they are allowed to go through at most *n* matching rounds in a generation to find compatible mates and produce offspring for the next mating generation. Once an individual finds a matching mate, it is taken out of the eligible mating pool. At the end of *n* matching rounds, all the unmatched individuals die without offspring. We are interested in how the population composition changes after *i* generations. At the start of each generation and at the beginning of each matching round, the mating populations are normalized in the following way: *NA* is the normalized population ratio of niche *A*, and *NB*, the normalized population ratio of niche *B*, so that *NA* + *NB* = 1. *Ax* is the ratio of a mating-bias allele *X* in *NA*, and *Ay* is the ratio of a mating-bias allele *Y* in *NA*, so that *Ax* + *Ay* = 1. *Bx* and *By* are similarly defined for the normalized population *NB* in niche *B*, so that *Bx* + *By* = 1. Then *NAx* = *NA* × *Ax, NAy* = *NA* × *Ay, NBx* = *NB* × *Bx*, and *NBy* = *NB* × *By* represent the four different groups of normalized populations shown in Fig 5, such that *NAx* + *NAy* + *NBx* + *NBy* = 1. Because individuals in the sympatric population meet randomly, the probabilities of encounter, or the matching probabilities, among individuals of different groups in a matching round are the products of the normalized population ratios of the groups. These are represented by the arrowed lines in Fig 5 with their associated matching probability values.

**Fig 5.**
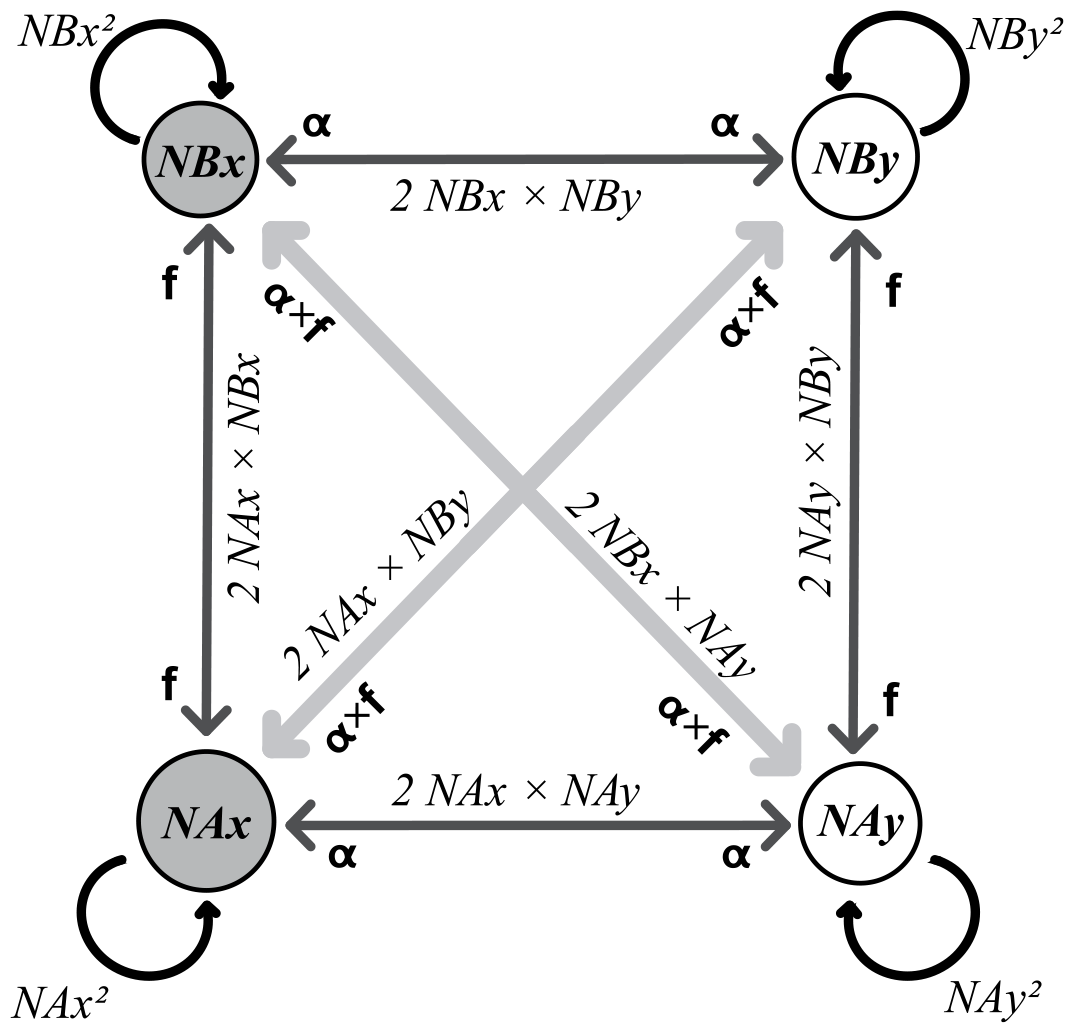
Mathematical model of a 2-mating-bias-allele, 2-ecological-niche sympatric ecosystem. The schematic diagram illustrates the interactions among the different niche groups. Reproductive isolation requires *NAx* and *NBy* to be relatively large, and *NBx* and *NAy* to be relatively small, or vice versa.

The symbol *f* at the heads of the arrowed lines specifies the ratio of offspring that a group *A* or a group *B* parent can expect to recover from a matching encounter, assuming that a matched individual from each group produces only one offspring, i.e., a”unit offspring,” to replace itself. For our 3-ecological-locus individuals in the computer simulation, if there are no viable hybrids (meaning all the hybrid offspring are eliminated because there is no niche for their ecological genotypes), *f* = 1/8, and the unit offspring ratio produced from the encounter between *NAx* and *NBx* is ⅛ × 2 × *NAx* × *NBx*. In the model, *f* is a measure of the strength of ecological selection; the smaller the *f*, the stronger the ecological selection.

Similarly, *α* in Fig 5 denotes the ratio of matched individuals from an encounter, based on a prespecified matching compatibility table (Fig 3). For instance, the unit offspring ratio that group *NAx* and group *NAy* can expect to produce from their encounter is 2 × *α* × *NAx* × *NAy*. For groups that belong to different ecological niches and possess different mating-bias alleles (e.g., *NAx* and *NBy*), the expected ratio of unit offspring that will be produced from their encounter is a multiple of *α* × *f* (e.g., 2 × *α* × *f* × *NAx* × *NBy*). In the model, variable *α* determines the strength of sexual selection.

The sum of all the matching probabilities is equal to 1: *NAx*^2^ + *NAy*^2^ + *NBx*^2^ + *NBy*^2^ + 2 *NAx NBx* + 2 *NAx NAy* + 2 *NBx NBy* + 2 *NAy NBy* + 2 *NAx NBy* + 2 *NBx NAy* = 1. Without the influence of the multipliers *α* and *f* (i.e., when *α* = 1 and *f* = 0.5), the sum of unit offspring ratios produced by all the matched encounters equals 1.

We can analyze the model in Fig 5 to gain an intuitive sense of how ecological selection (post-zygotic RI, as determined by *f*) and sexual selection (premating RI, as determined by mating-bias variable *α* and assortative-mating-cost variable *n*) may interact to produce reproductive isolation. If we let *NAx* represent the largest of the four population groups, then to have maximum reproductive isolation, we want *NAx* and *NBy* to be as large as possible, and *NBx* and *NAy* to be as small as possible, so that, in the limiting case, all the individuals in niche *A* possess the *X* allele and all the individuals in niche *B* possess the *Y* allele. In the first matching round, if each matched individual is only allowed to reproduce one offspring to replace itself (a unit offspring), then the presence of an *f* value that is less than 0.5, will increase the offspring ratio of the larger group more than the offspring ratio of the smaller group. For example, for groups *NAx* and *NBx*, given that *NAx* > *NBx*, the following ratio relationship holds:

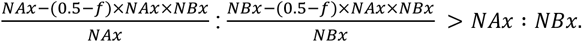

The net effect is that whenever there is loss (in this case, the loss of nonviable hybrid offspring) in the interaction between two groups, the interaction will make the larger group get larger and the smaller group get smaller. The smaller the *f* value, the greater the amount of loss, and the larger the change in the relative ratios of the two groups. The same applies to interactions mediated by the mating-bias factor *α*. If all the unmatched individuals never get another chance to mate after the first matching round, they perish without offspring. In this case, the loss will also make the larger group larger and the smaller group smaller. For example, if *α* < 1, since *NAx* > *NAy*,

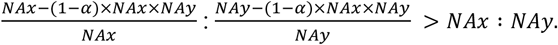

The loss of potential unit offspring is most severe in matches between groups that belong to different niches and possess different mating-bias alleles, i.e., *NAx* × *NBy* and *NBx* × *NAy*, where the loss multiplier is (1 − *α*) + (0.5 – *f*). However, the unit offspring ratios produced from these matches are distributed equally among the four groups. For example, the match between *NAx* and *NBy* adds ¼ × (*α* × *f*) × (2 × *NAx* × *NBy*) unit offspring to each of the four groups *NAx, NAy, NBx*, and *NBy*. This is because random assortment and inheritance of the ecological alleles and mating-bias alleles of the matched parents distribute offspring equally among the four groups. When individuals are matched with individuals from the same group, such as *NAx* × *NAx*, there is no loss of unit offspring. Therefore, only the relative ratio of the group becomes larger. Given our understanding of how the matching interactions among the different groups may affect their offspring’s relative proportions, if the initial normalized populations *NAx* and *NBy* are greater than *NBx* and *NAy*, ecological selection and sexual selection will act in concert to make *NAx* and *NBy* bigger and *NBx* and *NAy* smaller to achieve maximum reproductive isolation. The smaller the values of *f* and *α*, the stronger the selection pressures are to accelerate this process.

Next, we investigate the effect of having multiple matching rounds *n* on the relative unit-offspring ratios of the four groups. After the first matching round, the ratio of unmatched individuals, represented by *P*_1_, is determined by the following equation:

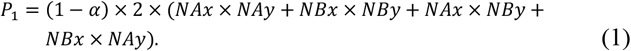

Because *P*_1_ is derived from unsuccessful matches between individuals that possess different mating-bias alleles, and because each of the two groups that participated in an unsuccessful encounter contributed equally to the probability of encounter, the ratio of *X* alleles in *P*_1_ is 1/2, the ratio of *Y* alleles in *P*_1_ is 1/2, and the ratios of *NAx*: *NBx* and *NAy*: *NBy* are preserved in the unmatched mating pool (see Appendix for a more rigorous mathematical derivation).

After *n* matching rounds, the ratio of unmatched individuals that are eliminated by sexual selection without producing offspring is:

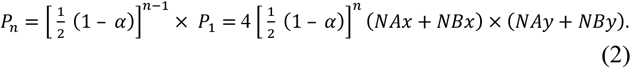

When *α* is less than 1, having multiple matching rounds reduces the number of unmatched individuals. Thus, the variable *n* can be a measure of assortative mating cost. The smaller the number of *n*, the more severe the sexual selection due to mating bias *α*, and more unmatched individuals will die without leaving offspring. However, even for *n* = *∞* (i.e., every individual gets to mate and produce offspring, or no sexual selection in the matching process), differential elimination of the mating-bias alleles in the niche groups can still occur in ecological selection. This is because the large number of matching rounds can alter the offspring ratios of the population groups in Fig 5 and cause uneven elimination of the mating-bias alleles in the niche offspring by ecological selection (as determined by *f*) and the limited niche carrying capacities (as determined by *NA*). Consequently, for large values of *n*, mating-bias allele assortment between niche ecotypes is still possible even though no individuals are eliminated (unmatched) because of mating-bias-allele incompatibility (low *α*) in the matching rounds.

Nevertheless, so far the system in Fig 5 is inherently unstable. Consider the situation when *NAx* and *NBy* have become the remaining dominant groups. Unless their sizes are perfectly equal, the slightly larger group is bound to eliminate the smaller group through their attritive interactions. This is because the larger group always loses a smaller ratio of its offspring through selection than that of the smaller group, and eventually only one population group will remain in the ecoscape. Such a process creates incumbent selection, through which the larger group exerts negative selection pressure on the smaller group. Incumbent selection occurs when a larger group of individuals can use its numerical advantage, established through historical legacy or incumbency, to exert negative selection pressure on the fitness of a smaller group of individuals that it interacts with. This can occur even in cases when the ecological fitness of the individuals in the smaller group, as measured by their abilities to extract local resources, is higher than the ecological fitness of the individuals in the larger group.

The effect of incumbent selection can be limited if the size ratios of the groups can be constrained by the carrying capacities of their respective niches. Suppose niche *A* only has enough resources to support a maximum population ratio *PAmax*, and niche *B* only has enough resources to support *PBmax*. Let variable *off* specify the number of offspring that an individual can reproduce in a generation. Then the number of offspring that a matched individual can produce in a generation is the product of its unit offspring ratio and *off*. If *off* is sufficiently high, such that each mating generation produces enough offspring to saturate the carrying capacities of all niches, then the normalized niche population ratios at the beginning of each mating generation are fixed by *NAmax* = *PAmax*/(*PAmax* + *PBmax*) and *NBmax* = *PBmax*/(*PAmax* + *PBmax*). Such a constraint makes it possible for a stable polymorphism to exist in the system, instead of having a dominant group that will eventually cannibalize the rest of the groups.

Fig 6 shows how the sexual selection and offspring reproduction portions of the life-cycle diagram in Fig 1 are computed in our mathematical models. The mating population is normalized at the beginning of each matching round. The probabilities of encounters of the different normalized groups (*NAx, NAy, NBx, NBy*) can be represented by the elements of a matching probability matrix *M* (Fig 7), which is the product of a vector [*NAx NBx NAy NBy*]^T^ and its transpose. The probability of a successful match of two individuals in an encounter is determined by the compatibility of their mating-bias alleles, specified by a matching compatibility table, as shown in Fig 3. At the end of each matching round, all the pairs of successfully matched individuals are collected. Let us specify a matrix *ΣM*_(*n*)_ the elements of which are the cumulated sums of all the different types of matched pairs of individuals at the end of matching round *n*, represented in a probability matrix format similar to *M*. Next, we can use a unit offspring matrix *U* (Fig 8) to calculate the composition of unit offspring that are produced by each type of matched pairs of individuals in *ΣM*_(*n*)_, according to independent assortment at each locus of the parents’ ecological alleles and mating-bias alleles.

**Fig 6.**
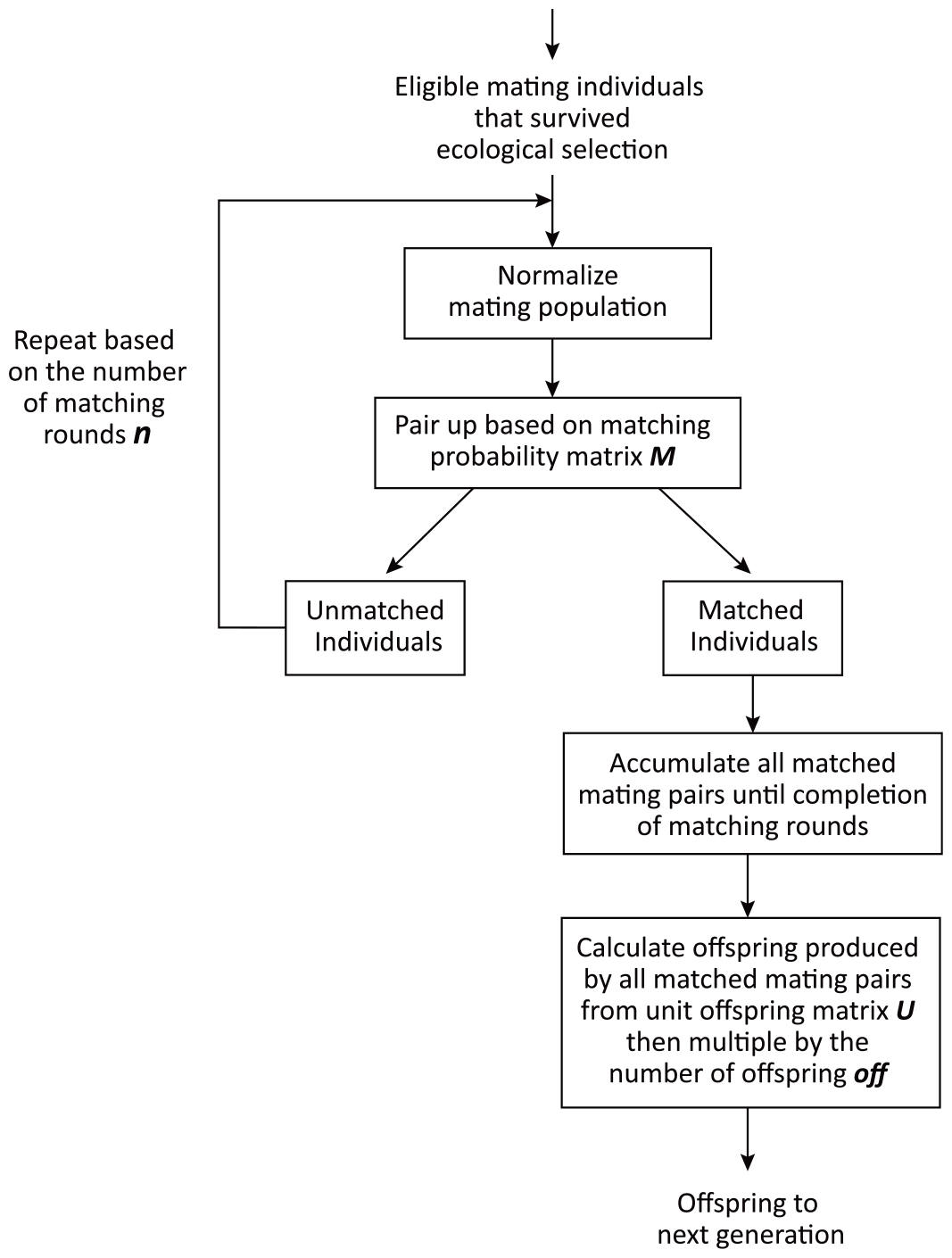
A flowchart showing the algorithm used to match eligible mating individuals and reproduce offspring.

**Fig 7.**
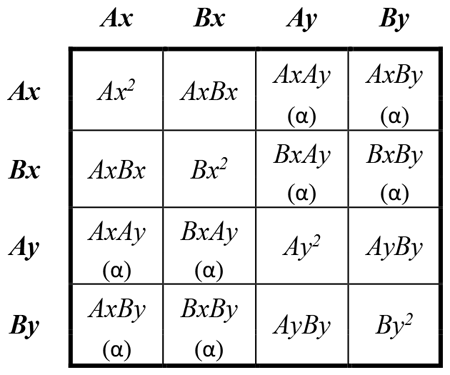
Matching probability matrix *M* for a sympatric model with no viable hybrids. The value in each cell denotes the probability of encounter as a product of its corresponding normalized population groups on the horizontal and vertical headers. Because *Ax* + *Bx* + *Ay* + *By* = 1, all the values of the cells add up to 1. Variable *α* specifies the probability of a successful match between two individuals that have different mating-bias alleles.

**Fig 8.**
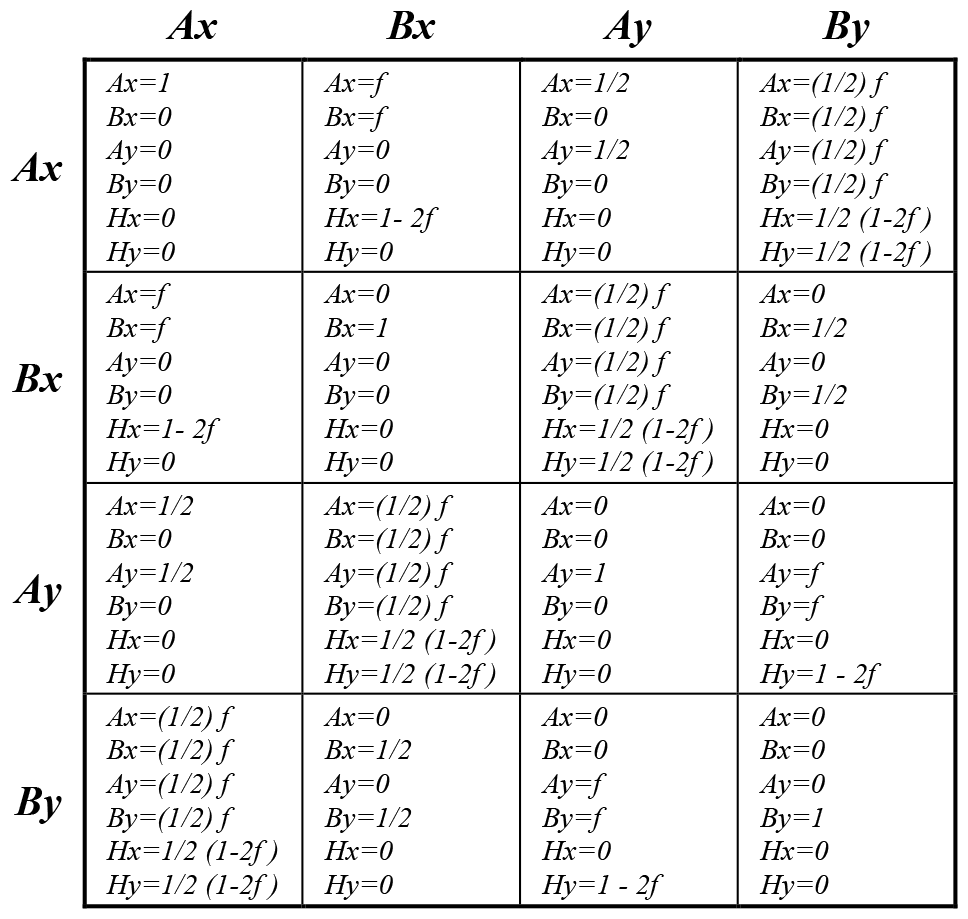
Unit offspring matrix *U*. Each cell of the matrix specifies the types and ratios of offspring that each matched pair of parents, shown on the horizontal and vertical headers, will produce. *Hx* and *Hy* are the ratios of hybrid offspring with the *X* and *Y* alleles. All the ratios in a cell add up to 1.

Having “unit offspring” means that each matched individual is only allowed to reproduce one offspring to replace itself. As shown in Fig 8, each cell of the unit offspring matrix *U* specifies the ratios of the different types of offspring (*Ax, Bx, Ay, By, Hx*, or *Hy*) that each combination of matched parents will produce, based on independent assortment of alleles and the ecological selection variable *f*. Notice that all the ratios in each cell of *U* add up to 1. Therefore, if *α* = 1 or *n* = ***∞***, that is, everyone in the population gets to mate, then a normalized mating population of 1 will also produce a total unit-offspring population of 1. The unit offspring produced may be expressed mathematically as *ΣM*_(*n*)_ ↦ *f* (*U*), where the symbol ↦ *f* (*U*) denotes the element-wise multiplication of the elements of the matrix *ΣM*_(*n*)_, which specify the probability ratios of the different types of parental matches, by the offspring proportions in each corresponding cell of *U* to produce different types of offspring. The different types of offspring are then added up to produce output vectors 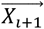 and 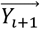 that show the total unit offspring (unnormalized) produced from the *i*^*th*^ generation. For instance, we can decompose *U* into submatrices that only contain the ratios of individual offspring types: *U* = *U*_*Ax*_ + *U*_*Hx*_ + *U*_*Bx*_ + *U*_*Ay*_ + *U*_*Hy*_ + *U*_*By*_. Then

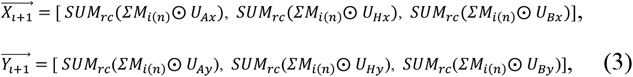

where ⨀ is the Hadamard product that performs element-wise multiplication of two matrices of the same size. The operation *SUM*_*rc*_(*A*) sums all the elements of matrix *A. ΣM*_*i*(*n*)_ = *ΣM*_(*n*)_ in generation *i*.

Expanding the equations and using the formula of geometric series, a set of difference equations can be derived that describe the nonlinear dynamic behavior of the 2-mating-bias-allele, 2-ecological-niche sympatric population with no viable hybrids (see Appendix for detailed mathematical derivation):

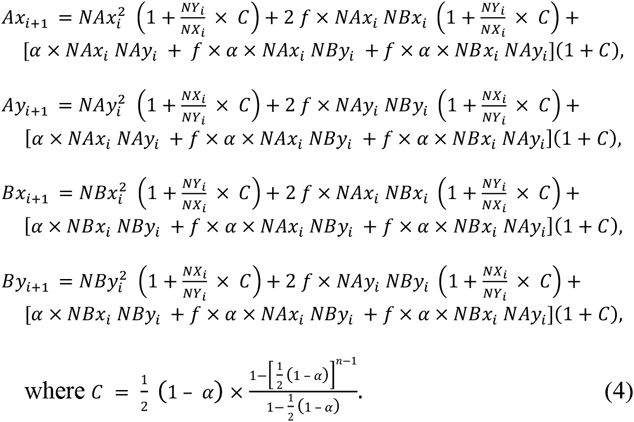

The variables *NAx*_*i*_, *NAy*_*i*_, *NBx*_*i*_, and *NBy*_*i*_ are the normalized population ratios at the start of mating generation *i*. They are the normalized offspring ratios from generation *i* − 1, and they represent genotypes that have survived ecological selection and sexual selection from the previous generation. In the equations, *NX*_*i*_ = *NAx*_*i*_ + *NBx*_*i*_ and *NY*_*i*_ = *NAy*_*i*_ + *NBy*_*i*_. The variables *Ax*_*i*+1_, *Ay*_*i*+1_, *Bx*_*i*+1_, and *By*_*i*+1_ are the ratios of unit offspring (not normalized) that are produced after *n* matching rounds in generation *i*.

For special cases when *NAx*_*i*_ + *NBx*_*i*_ = *NX*_*i*_ = 0 or *NAy*_*i*_ + *NBy*_*i*_ = *NY*_*i*_ = 0, i.e., when the *X* or the *Y* allele is absent:

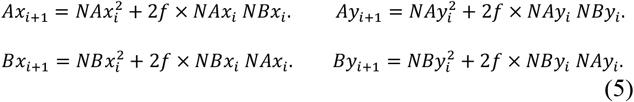

If *off* is large enough so that the number of offspring produced in each generation is able to completely saturate the carrying capacities of the two niches, *NAmax* and *NBmax*, then the normalized *NAx, NAy, NBx*, and *NBy* ratios going into the next mating generation can be calculated as follows:

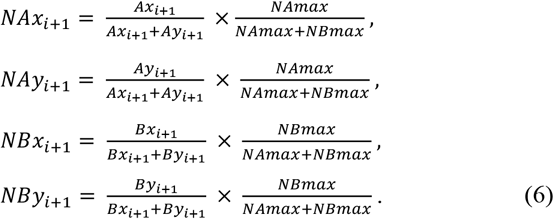

Finding the analytical solution to the set of nonlinear difference equations (4)-(6) has proven to be difficult. For this reason, we developed a user-friendly computer program to numerically calculate and display the solution. Fig 9 shows the general layout of an interactive program created using MATLAB App Designer. It displays a phase portrait (vector field) solution of the equations based on various input parameters. The graph plots *Ax*, the ratio of individuals with *X* alleles in niche *A*, against *Bx*, the ratio of individuals with *X* alleles in niche *B*. As shown, the vector lines trace the change in *Ax* and *Bx* values of different populations as they progressed through 100 mating generations, starting from different initial coordinate points (*Ax, Bx*) that are spaced at 0.1 intervals apart on the *Ax* and *Bx* axes. Three basic patterns of dynamic behavior emerge as the equation parameters are varied.

**Fig 9.**
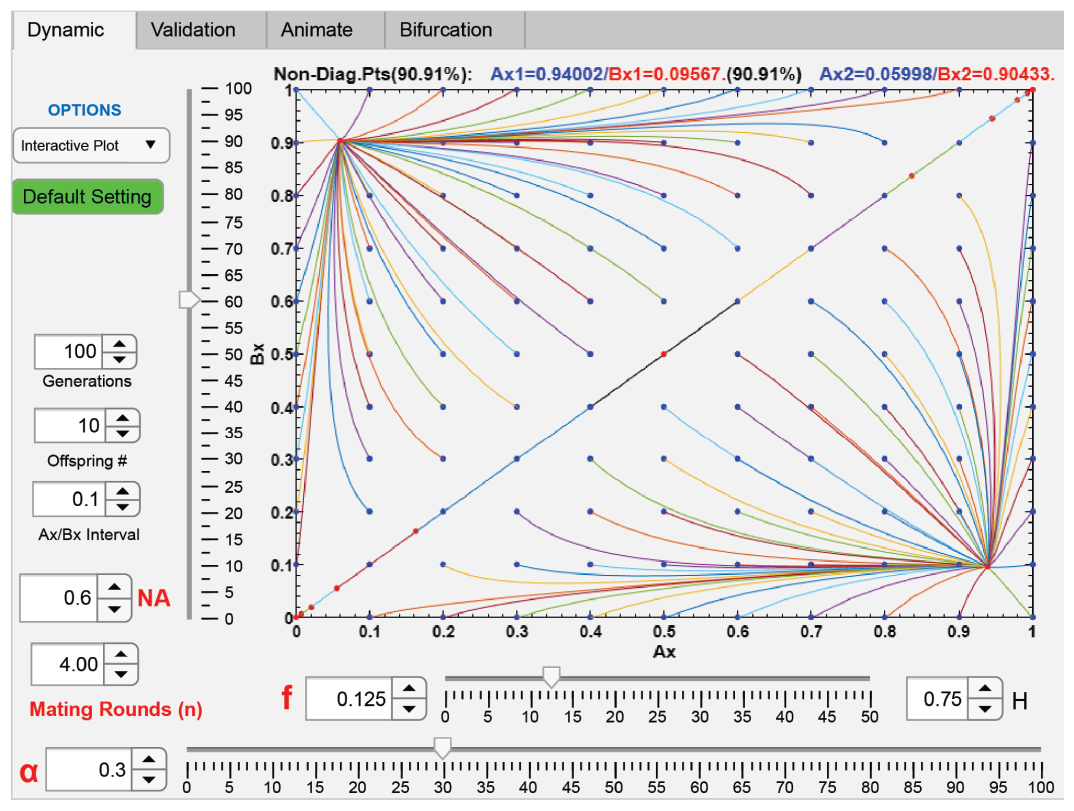
Phase portrait solution of a 2-niche sympatric system without viable hybrids. A computer screenshot shows a phase portrait (vector field) solution of the difference equations derived from a 2-niche model without viable hybrids. The vector lines begin at specified initial values of *Ax* and *Bx*, which in this example are spaced at intervals of 0.1. The lines converge to two fixed points on either side of the diagonal line where *Ax* = *Bx*. Above the plot, the two fixed-point values for *Ax* and *Bx* are displayed. The proportion of the initial values of *Ax* and *Bx* in the vector field that eventually converge to fixed points is expressed as a percentage (convergence percentage). The values of the various input parameters used in the equations, *NA, f*, α, *n, i*, and *off*, are also displayed. *H* is the ratio of hybrid offspring (1 − 2 *f*).

First, when selections are relatively strong with low values of *α* and *f*, all the non-diagonal initial conditions converge to a fixed point (a sink node) on either side of the diagonal line (see Fig 9). In this case, we can call the system globally convergent, i.e., convergent to fixed points, or simply “convergent.” The coordinates of the fixed points are solved numerically and displayed above the phase portrait in Fig 9. The two fixed points are mirror images of each other across the diagonal line. In our model, to have maximum reproductive isolation between the two niche groups, the lower fixed point needs to be as close to the far-right lower corner as possible (where *Ax* = 1 and *Bx* = 0), which can only be reached if either *α* = 0 or *f* = 0. Any nonzero combinations of *α* and *f* will produce stable mating-bias-allele polymorphism at the fixed points.

The program has an animation feature that displays the changes in *Ax* and *Bx* coordinates on the population vector lines in real time (Fig 10). It turns out that *Ax* = *Bx* = 0.5 is an unstable source node in the dynamic landscape. With the passing of generations, diagonal-line points on either side of this source node will be driven solely by sexual selection to diverge toward opposite ends of the diagonal line, to the corners where *Ax* = *Bx* = 0 and *Ax* = *Bx* = 1. (In our model, ecological selection, as determined by the value of *f*, has no effect on points on the diagonal line.) This is because sexual selection favors the most numerous mating-bias allele in the population. The most numerous type of mating-bias allele will have more chance to match and out-reproduce the less numerous types of mating-bias alleles. From the computer animation, it can be seen that the speed of this divergence is inversely proportional to the values of *α* and *n*, which jointly determine the strength of sexual selection. In Fig 9, the diagonal line, as defined by *Ax* = *Bx*, is an unstable ridge. Any point that deviates ever so slightly from the diagonal line will be drawn into one of the two fixed-point attractor basins on either side of the ridge and converge to a fixed point with time.

**Fig 10.**
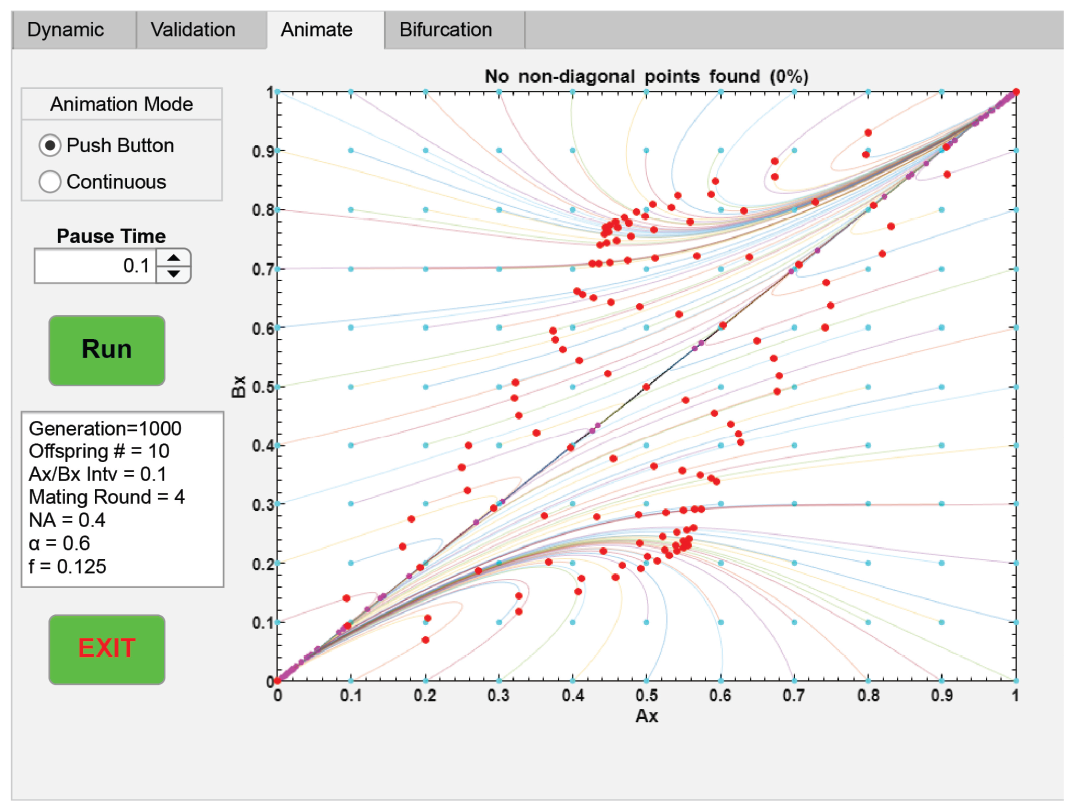
A computer screenshot showing an animation feature of the program. The dots, which represent the normalized ratios of *Ax* and *Bx* in different populations, are programmed to move along the vector lines in real time with the passing of each generation. The input parameters are displayed in the table insert.

Second, if we gradually reduce the ecological and sexual selection pressures by increasing the values of *α* and *f*, eventually we will reach a point where no fixed points exist in the dynamic vector field (see Fig 11). In this case, all the non-diagonal points move toward the diagonal line and are driven by sexual selection to diverge toward the ends of the diagonal line, where all niche groups possess the same mating-bias allele. (In the special case that *α* = 1 or *n* = ***∞***, no sexual selection exists on the diagonal line, and the points will come to rest at stationary positions on the diagonal line.) We can call such a system globally divergent, or simply “divergent.”

**Fig 11.**
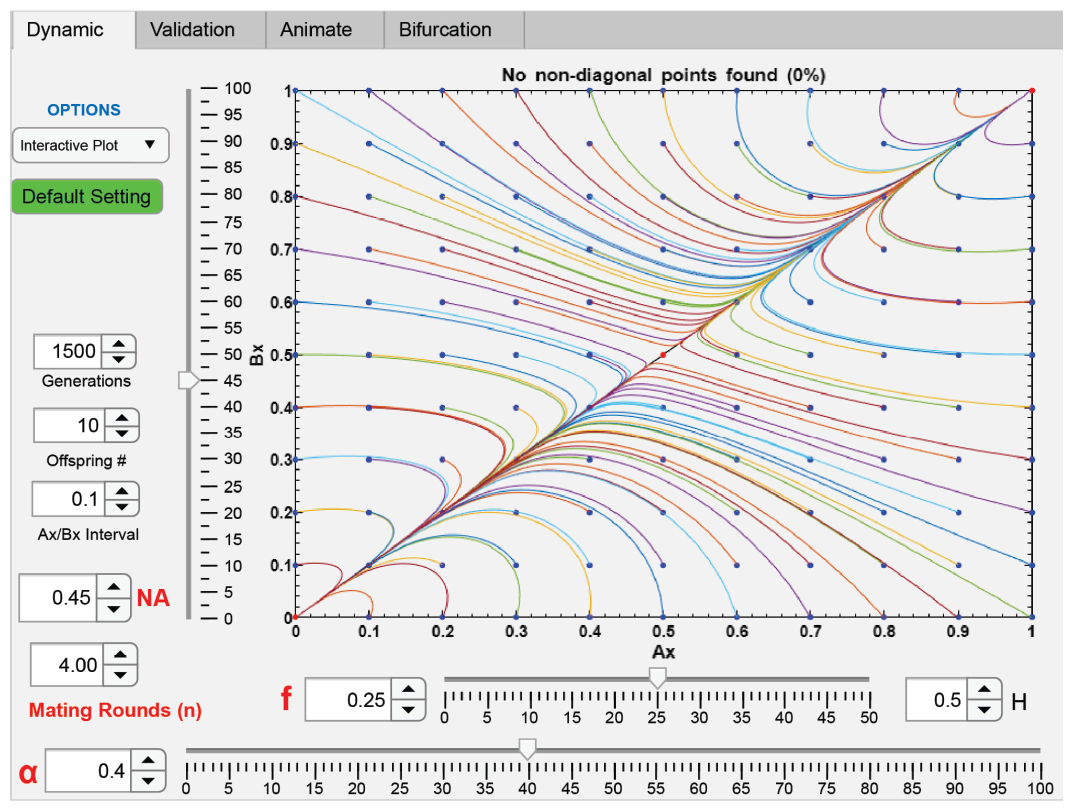
A computer screenshot showing nonconvergence. No fixed points exist. *Ax* = *Bx* = 0.5 is an unstable equilibrium point. Sexual selection drives all points on the diagonal line toward the ends of the line, and eventually, only one type of mating-bias allele remains in the population.

Third, we can find the bifurcation thresholds when our nonlinear dynamic system transitions from being globally convergent to being globally divergent. We can apply numerical methods to plot a curve that traces all the fixed points of the equations over the range of a parameter. Fig 12 shows such a curve as we vary *α* from 0 to the bifurcation threshold *α* = 0.59895 (displayed at the top of the graph) beyond which the system becomes globally divergent and no fixed points are possible. As the fixed point moves toward the origin with decreasing degrees of reproductive isolation, the system can become “partially convergent,” with only a portion of the non-diagonal starting points converging to a fixed point (shown as solid lines in Fig 12) and the rest moving to the diagonal line (shown as faint lines). The percentage of non-diagonal points that converge to a fixed point, or the convergence percentage, is displayed at the top of the graph. In general, the transition from global convergence to partial convergence to global divergence is quick, and it tends to occur over a small interval of change in parametric value. If we zoom in at the bifurcation threshold by changing the plot axes and the starting-point intervals (*Ax*/*Bx Interval*), we can see a subcritical Hopf bifurcation just before the fixed point disappears (Fig 13).

**Fig 12.**
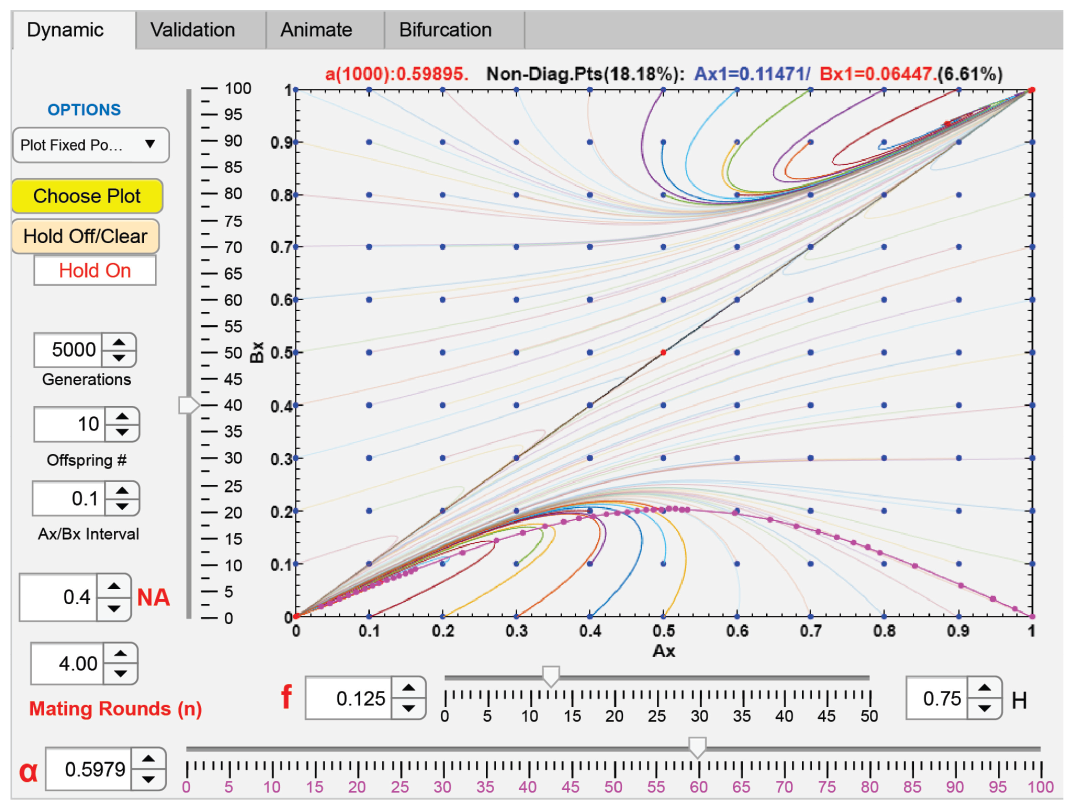
A computer screenshot showing a fixed-point curve as the value of *α* is varied. The input parameters are *NA* = 0.4, *f* = 0.125, and *n* = 4. The dots on the curve are the solved iteration stops during the numerical calculation. The upper limit of *α* (i.e., *α* = 0.59895) beyond which the algorithm could no longer find a fixed point is displayed at the top of the graph. The value in the parentheses is the maximum number of generations that the algorithm used to find the fixed points. The vector field is partially convergent. The solid lines show vector populations that eventually converged to a fixed point (at *Ax* = 0.11471 and *Bx* = 0.06447 in this example) near the upper limit of *α* (at *α* = 0.5979). The faint lines represent vector populations that never converged to a fixed point. The convergence percentage (18.18%) is also displayed.

**Fig 13.**
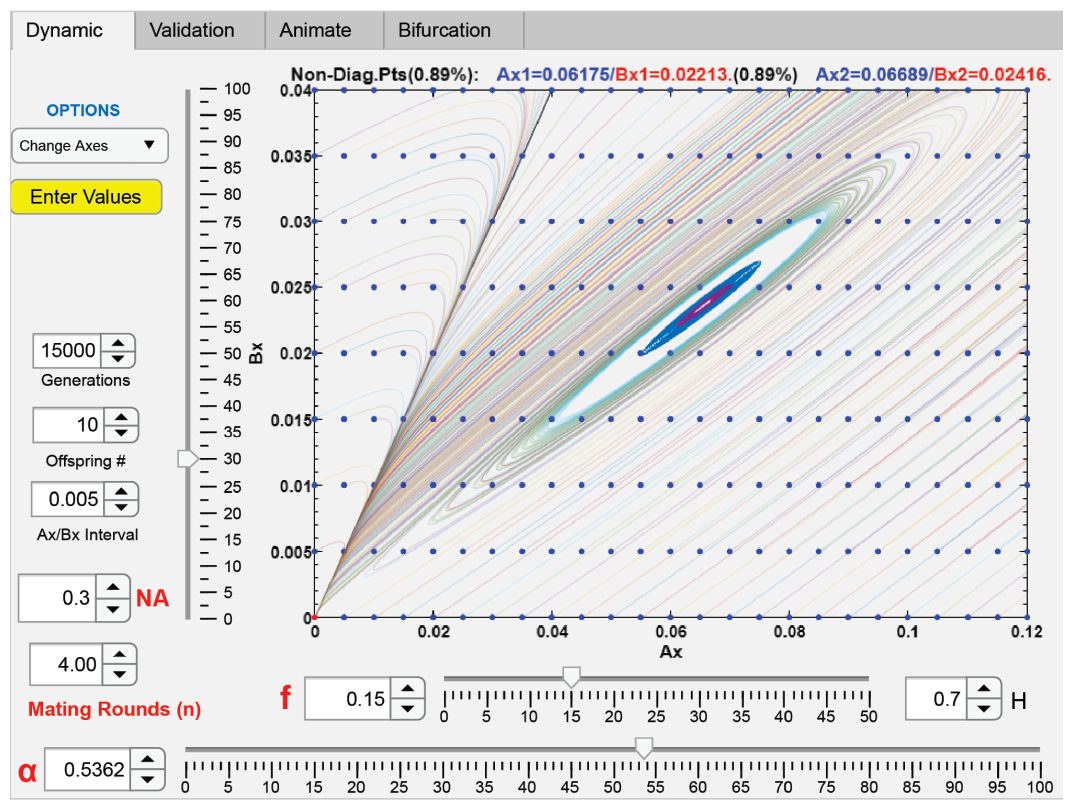
A computer screenshot showing a subcritical Hopf bifurcation. A stable fixed point and an unstable limit cycle appear near the origin (*Ax* = *Bx* = 0) when the values of *α* and *f* are relatively large. All the initial values of *Ax* and *Bx* outside the unstable limit cycle diverge (shown as faint vector lines).

We can obtain similar fixed-point plots for parametric variables, *f, NA*, and *n*, and their bifurcation thresholds. Fig 14 shows the general shapes of such curves for parametric variables *NA* and *NA*. The fixed-point curves for variable *α* are very similar to those for variable *f*. When plotted with different values of *NA*, all the fixed-point curves for *f* (or for *α*) seem to terminate at approximately the same bifurcation value of *f* (or *α*). As previously described, convergence percentages tend to decrease rapidly at the tail ends of the curves. The shape of the fixed-point curves for the variable *NA* is a parabola, and the mid-point of the parabola, where *NA* = 0.5, is the point closest to the coordinate of maximum reproductive isolation at *Ax* = 1 and *Bx* = 0. If everything else is equal, when *NA* equals 0.5, or when the populations of groups *A* and *B* are equal (therefore have maximum matching interaction between them and maximum hybrid loss in the system), the system is most likely to converge to a fixed-point polymorphism. As the value of *NA* moves away from 0.5, and the fixed points move toward the ends of the parabolas, the percentage of convergence rapidly vanishes.

**Fig 14.**
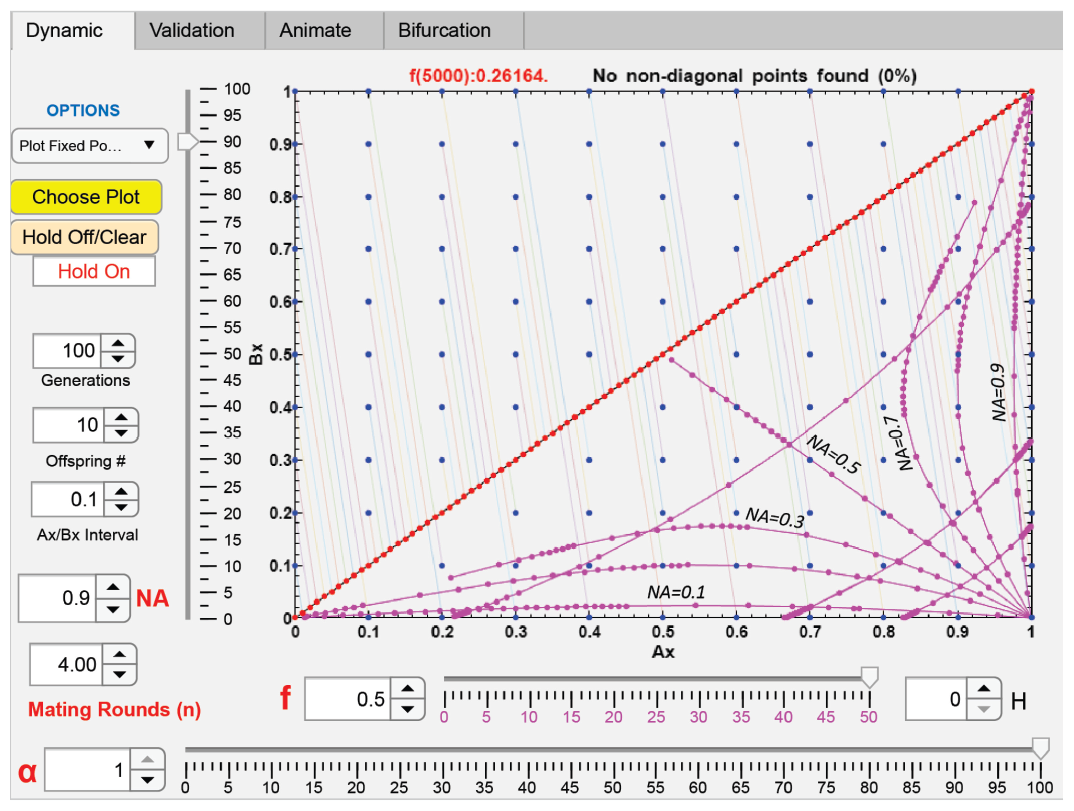
A labeled computer screenshot showing representative shapes of fixed-point curves. Fixed-point curves obtained by varying the value of *f* tend to be similar in shape to the fixed-point curves obtained by varying the value of *α*. Here, they are shown for various values of *NA*. The curves originate at the coordinate *Ax* = 1 and *Bx* = 0, when the value of *f* (or *α*) is zero, and radiate outward as the values of *f* (or *α*) increase. The curves seem to vanish at approximately the same upper values of *f* (or *α*) Crisscrossing these curves is a set of parabolas. These parabolas represent the typical shapes of fixed-point curves that are calculated by varying the value of *NA*. The percentage of convergence diminishes as the value of *NA* increases or decreases away from 0.5 and the fixed points move toward the endpoints of the parabolas. As the values of *α* and *f* increase, the fixed-point parabola for *NA* moves toward the diagonal line.

The fixed-point curves for the variable *n* (the number of matching rounds) tend to be short and invariant. This is because, with each successive matching round, the portion of unmatched individuals 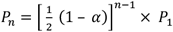 decreases rapidly as an exponential function of *n*, so does its effect on the dynamics of the system. Notably, for many parametric values, the system will not converge if *n* = 1, when sexual selection is strongest, but will converge when *n* is greater than 1 (e.g., for *NA* = 0.6, *f* = 0.4, and *α* = 0.1, the system only converges for *n* ≥ 6). Therefore, very strong sexual selection caused by high assortative mating cost (very low values of *n*) and very weak sexual selection caused by low mating bias (high values of *α*) can both inhibit the development of mating-bias-allele polymorphism and premating RI. These findings align with prior research suggesting that intermediate values of sexual selection are most conducive to the development of reproductive barriers [14, 19, 22, 28, 45, 70, 71].

The ability to plot fixed-point curves for all parametric variables enabled us to numerically solve all the fixed points and bifurcation thresholds in the normalized sympatric model. Fig 15 shows an illustrative 3-D surface and contour plot of all the fixed points as we specify *n* and *NA* and vary *f* and *α* on the *x* and *y* axes. The *z* axis shows the absolute magnitude *abs* (*Ax* − *Bx*) at each fixed point, and the color scale of the surface plot shows the percentage of non-diagonal initial (*Ax, Bx*) points in the vector field that will converge to the fixed point. Fig 16 shows a 3-D volume plot of the fixed points by specifying *n* and varying the normalized values of *f, α*, and *NA* on the *x, y*, and *z* axes. The color scale of the volume plot indicates the percentage of convergence at the fixed points.

**Fig 15.**
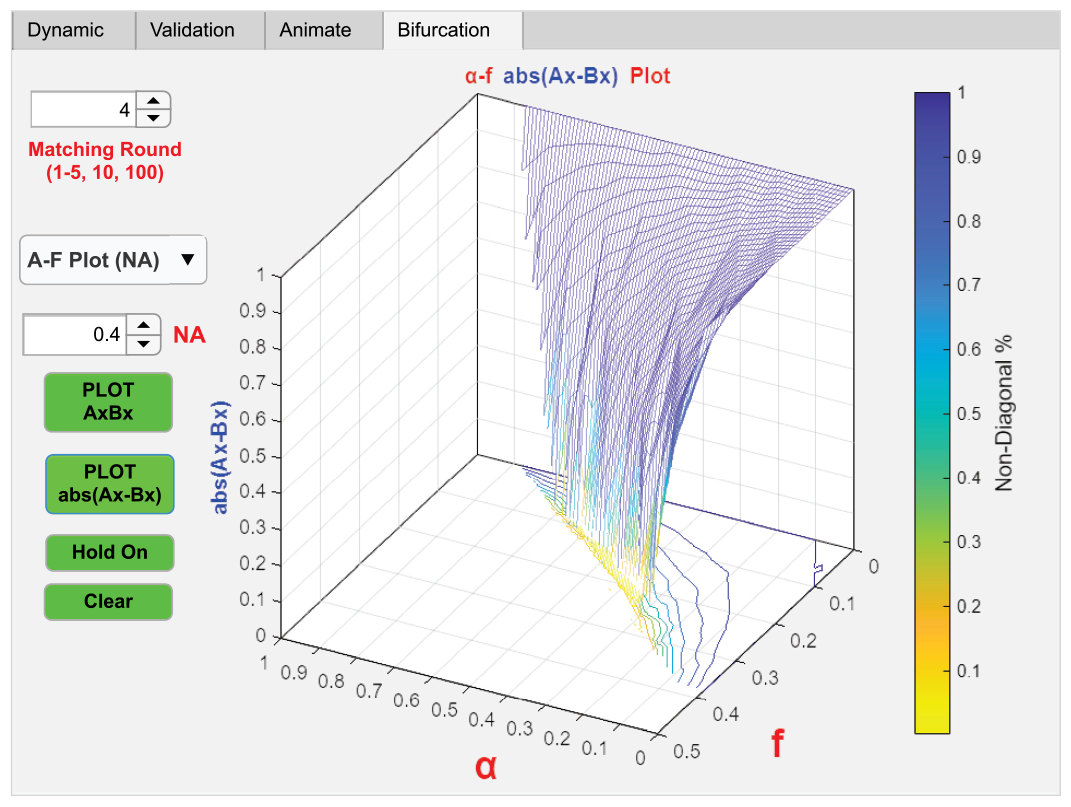
A surface and contour plot of the difference between *Ax* and *Bx* at fixed points. The value of *abs* (*Ax* − *Bx*) is a measure of the degree of RI as the values of *α* (the degree of mating compatibility) and *f* (an ecological-selection factor) are varied. The values of *NA* (the ratio of niche A population) and *n* (the number of matching rounds) are specified. The percentage of convergence at each fixed point is denoted by the color on the side colorbar.

**Fig 16.**
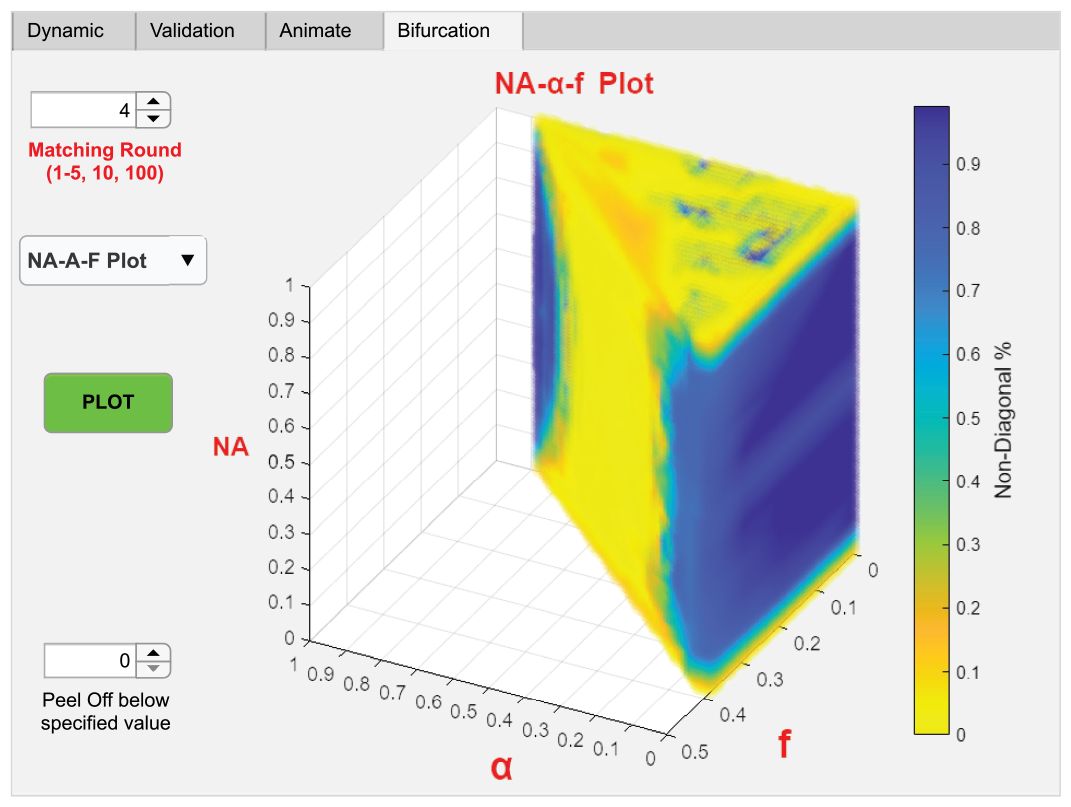
A computer screenshot showing a three-dimensional volume plot of fixed points. The axis variables are the normalized parameters *NA* (the ratio of niche *A* population), *α* (the degree of mating compatibility), and *f* (an ecological-selection factor). To create the plot, the only input variable that needs to be specified is *n*, the number of matching rounds. The percentage of convergence at each fixed point is denoted by the color on the side colorbar. The computer application allows for the peeling away of fixed points that have convergence percentages below a specified value in the 3-D volume plot.

The results from these plots (Fig 15 and 16) indicate that when the number of matching rounds *n* is sufficiently high so that further increase in *n* will not significantly affect the dynamic behavior of the system, a diagonal line on the *α*-*f* plane, connecting the coordinates (*α* = 0, *f* = 0.5) and (*α* = 1, *f* = 0), demarcates two regions, one where fixed points can exist and another where fixed points cannot exist. The condition most conducive to achieving maximum RI is at the point where *NA* = 0.5, *α* = 0, and *f* = 0. As we move toward the diagonal demarcation border, the system tends to become partially convergent. The convergence time lengthens as we move toward the corners at (*α* = 1, *f* = 0) and at (*α* = 0, *f* = 0.5), and it becomes infinite when *α* = 1 (i.e., no sexual selection) or *f* = 0.5 (i.e., no ecological selection) at those corner locations. The system also tends to become partially convergent as we move toward the extreme ends of *NA*, i.e.,where *NA* = 0 and *NA* = 1. As the number of matching rounds *n* decreases toward 1, the volume of the triangular prism in Fig 16 shrinks in size toward the coordinate point *NA* = 0.5, *α* = 0, and *f* = 0. Therefore, very low values of *n* (less than 4 in this example) seem to impede the development of fixed-point polymorphism and reproductive isolation.

The origin (at *Ax* = 0 and *Bx* = 0) acts like an unstable saddle point when fixed points exist in the system and the origin falls within their basins of attraction (Fig 9 and 12). Any slight deviation from zero in the initial *Ax* or *Bx* values will draw the population toward a fixed point, which acts like an attractor (a sink) in the vector field. This means that in a 2-niche ecosystem that only has one type of mating-bias allele *Y* in its populations, a mutant mating-bias allele *X* that has a sufficiently strong mating bias can invade the system and drive it toward a fixed-point polymorphism of two mating-bias alleles.

Fig 17 shows a generational plot that tracks the change in *Ax* and *Bx* values given specified input parameters. In this illustrative example, if we set the initial *Ax* = 0.0001 and *Bx* = 0, the *X* allele will invade and achieve fixed-point polymorphism after 35 generations. Fig 18 shows a semi-log plot of the generation time needed for an invading mutant allele *X* to reach fixed-point polymorphism as a function of *α* and *NA*, as well as the corresponding fixed-point *Ax* and *Bx* values (expressed as percentages by multiplying their ratios by 100). As shown, the generation time to fixed-point polymorphism is shortest with low *α* and *NA* values. For a given value of *α*, the invasion generation time increases exponentially with *NA* until it reaches a transitional *NA* value, *NAt* (which is approximately 0.84 for *α* = 0.3 in our example), where it becomes infinite. After *NAt*, the population converges to the system’s mirror-image fixed point across the diagonal line in Fig 9, and the generation time gradually stabilizes at a high value.

**Fig 17.**
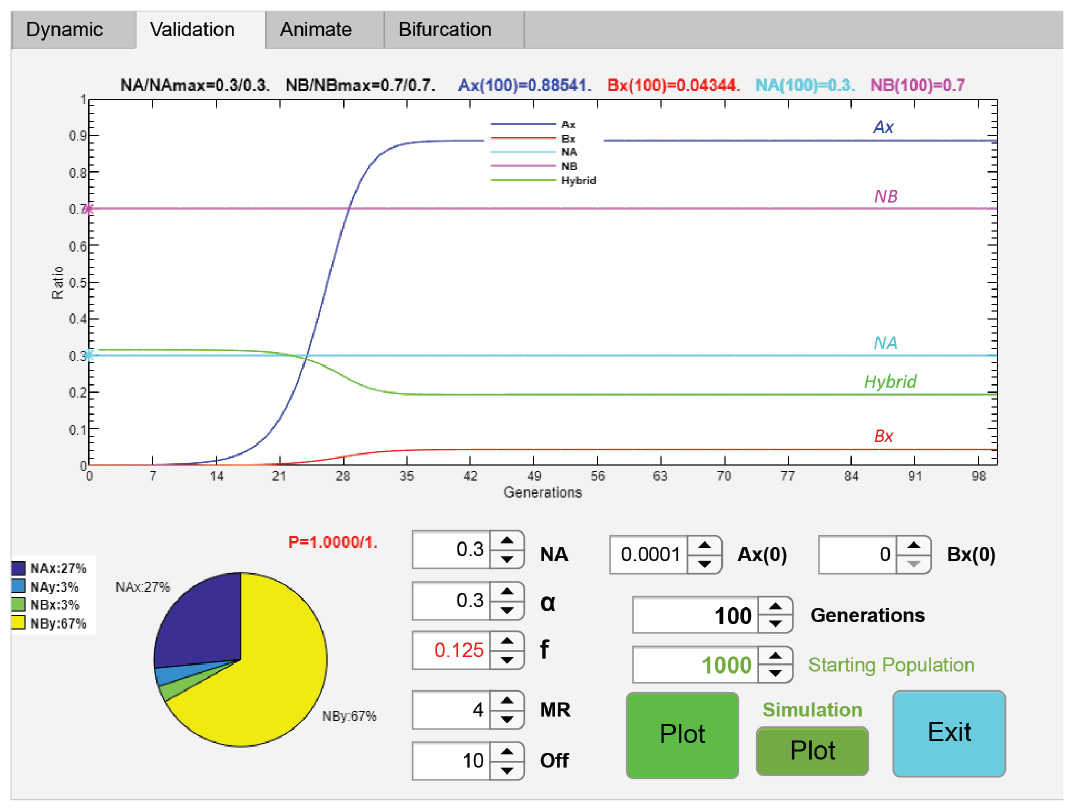
Change in population composition and allele ratios with generation time. A labeled computer screenshot shows how the composition of a sympatric population may change with generation time for a specified set of input parameters. *Ax*(0) and *Bx*(0) are the starting *Ax* and *Bx* values. The graph plots the changes in *Ax* (the ratio of *X* allele in niche *A*), *Bx* (the ratio of *X* allele in niche *B*), *NA* (the ratio of niche *A* population), *NB* (the ratio of niche *B* population), and the ratio of hybrid offspring (a measure of the degree of RI) produced in each generation. The fixed-point values of *Ax* and *Bx* are displayed at the top of the graph. *NA*/*NAmax* and *NB*/*NBmAx* are the ratios of final niche populations relative to their niche carrying capacities. If the origin (*Ax* = *Bx* = 0) is within the basin of attraction of a fixed point in the system’s vector field, then an initially small mutant population (specified as *Ax* = 0.0001 and *Bx* = 0 in this example) will invade a homogenous population that only has the *Y* allele and, with time, reach equilibrium polymorphism at the fixed point.

**Fig 18.**
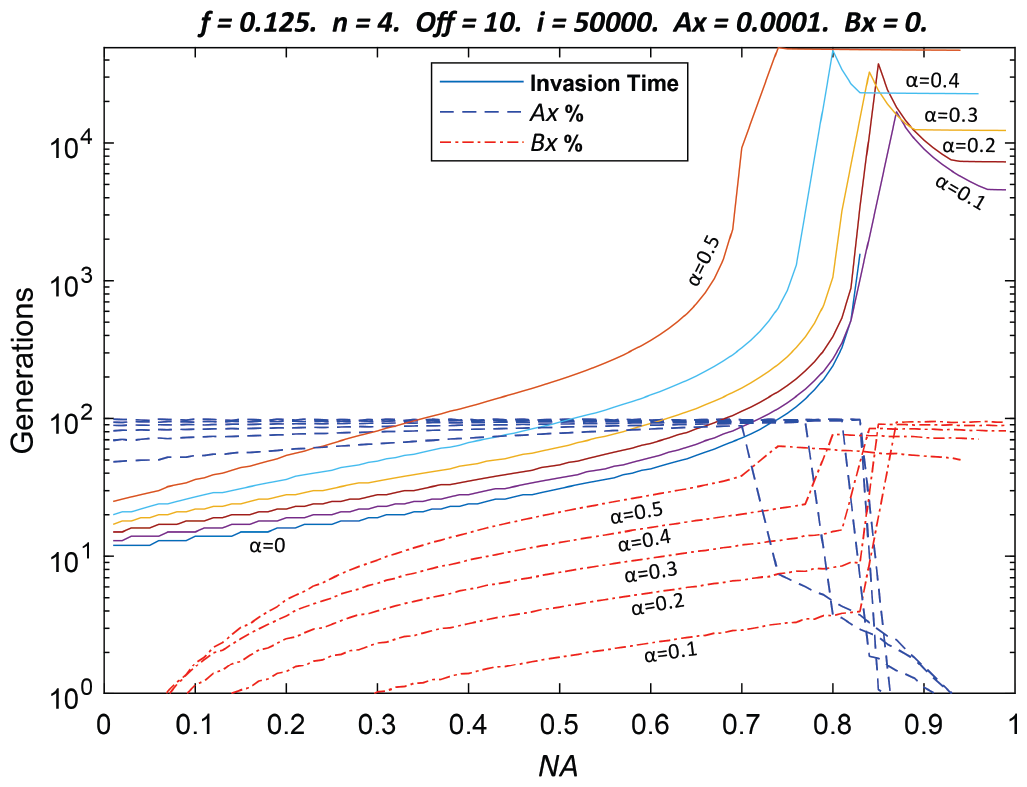
Invasion time as determined by mating bias and niche size. A labeled semi-log plot shows the generation times needed for mutant alleles with different mating-bias values to invade as a function of niche population ratios. Variable *α* specifies the mating-bias value. *NA* is the normalized population ratio of niche *A*. Displayed above the graph are the input parameters, specified using the variables in the mathematical model without viable hybrids. The generation time is displayed using a log scale on the vertical axis. The niche population ratio *NA* (which is the same as *NAmax* because the number of offspring produced in each generation is able to fill the niche’s carrying capacity) is displayed on a linear scale on the horizontal axis. The values of *Ax* and *Bx* at the equilibrium fixed points, achieved after an invading mating-bias allele *X* reached stable polymorphism with the existing *Y* allele, are plotted as percentages against different values of *NA* in the graph. For *α* > 0, the generation time increases exponentially with *NA* until it reaches a point when it becomes infinite (e.g., around *NA* = 0.84 for *α* = 0.3). Subsequently, the equilibrium fixed point flips to its mirror-image fixed point on the other side of the diagonal line in the vector field and the generation time stabilizes at a high value (e.g., at approximately 12,530 generations for an invading *X* allele with *α* = 0.3).

A potential explanation for the shape of the invasion-time curves in Fig 18 is that as *NA* decreases, the ratio of the smaller niche-*A* population that is lost to nonviable hybrids is much greater than that of the niche-*B* population (the “minority cytotype principle” [72, 73]), so that any beneficial high-mating-bias mutant allele *X* that emerges in niche *A* will be under greater positive selection pressure to multiply and stem this hybrid loss. As *NA* becomes substantially larger, and the population of niche *A* becomes the dominant group, then it will take time for the much larger niche-*A* group to transfer the mutant allele *X* to the smaller niche-*B* group through their limited matching interactions. Contrary to the situation when *NA* is small, a high-mating-bias allele *X* that emerges in a large population of *NA* does not have the benefit of a higher positive selection pressure through its linkage disequilibrium and runs a higher risk of being eliminated. A smaller niche-*A* population also means that there is a smaller probability for a mutant beneficial allele to emerge in the group. When the population in niche *A* is large, however, there is a higher chance of producing a mutant high-mating-bias allele in niche *A*, but it takes time to seed the now smaller population in niche *B* to produce fixed-point polymorphism.

Because individuals in our model are monogamous and matched individuals are taken out of the mating pool, when *α* is small, the chance of a mutant allele *X* in niche *A* matching with itself (the matching probability *NAx*^2^) is greatly increased after the first matching round, thus avoiding the offspring loss that comes from mating with the *Y* allele. In the limiting case when *α* = 0, there are no matches between the *X* and *Y* alleles, and the ratio of *NAx* in the unmatched population after the first matching round is 0.5 (versus *NAx* ≤ 0.0001 during the first matching round in the Fig 18 example). Furthermore, because *α* = 0, there is no way to introduce any *X* allele into niche *B*; therefore, *Bx* = 0 for all values of *NA*.

In the Fig 18 example, there is very little assortative mating cost for a small invading population of high-mating-bias (*X*) alleles after the first matching round. This is because virtually all the low-mating-bias (*Y*) alleles match in the first matching round and are removed from the mating pool. This phenomenon suggests that in our particular matching scheme, the cost of assortative mating, as determined by the number of matching rounds *n*, appears to be context- or frequency-dependent. When the invading mutant population is small, all individuals can mate even with low values of *n*, and the assortative mating cost is low. However, as the mutant population grows in size, the number of matching rounds begins to exert a more pronounced effect on the assortative mating cost in the system.

We can use a search-cost variable *θ* to extend the variable range of the assortative mating cost. In this way, we can create more graduated amounts of assortative mating cost to analyze its effects on the invasion dynamics of the system. In the matching compatibility table in Fig 3, we define a variable *θ* (similar to *α*) that specifies the ratio of successful matches between individuals that possess the same mating-bias alleles. The ratio of successful matches between individuals possessing different mating-bias alleles is then *α*^*’*^ = *α* × *θ*. Therefore, in a system that almost exclusively contains *Y* mating-bias alleles, not all the individuals possessing the *Y* allele will match in the first matching round. This is because a ratio of (1 − *θ*) unmatched individuals remains in the mating pool going into the next matching round. In real-world examples, increased search cost, as defined by *θ*, could be caused by an expanded geographic range or the turbidity of lake water that reduces the probability of matching encounters among individuals of a sympatric population.

Fig 19 shows an illustrative example of a computer application that plots the dynamic phase portrait of a 2-mating-bias-allele, 2-niche sympatric system without viable hybrids as the value of *θ* is varied. The rest of the parametric values are the same as those used in Fig 9. If *θ* = 1, there is no search cost, and the phase portrait is identical to that in Figure 9. If *θ* = 0, there is no matching encounter among individuals, no offspring is produced, and the population becomes extinct. If *α* = 1, there is no sexual selection, and all starting populations move to stationary positions on the diagonal line regardless of the value of *θ* (except for *θ* = 0). For values of *θ* between 0 and 1, the assortative mating cost is determined by the variables *θ, α*, and *n*. Fig 19 shows the phase portrait solution when *θ* = 0.8. As we gradually increase the search cost by lowering the value of *θ* from 1, starting populations near the ends of the diagonal line begin to converge toward the origin and its opposite end on the diagonal line, which means that invasion by a high-mating-bias mutant allele becomes more difficult.

**Fig 19.**
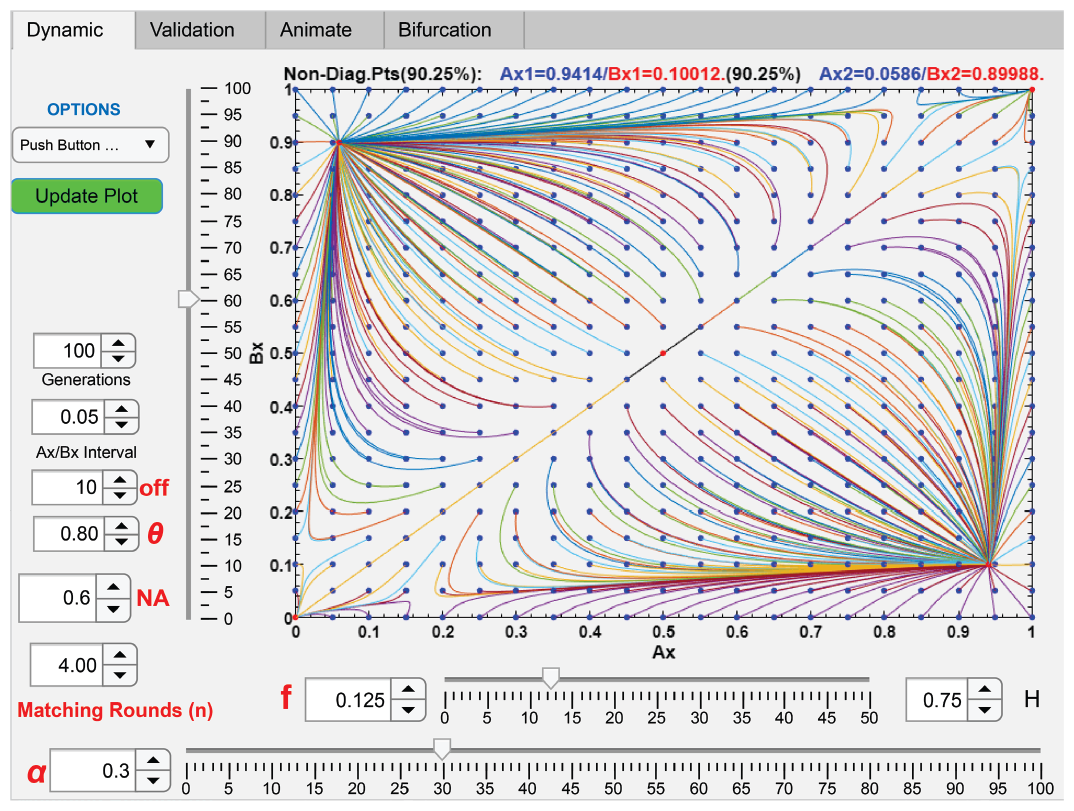
The effect of search cost on the population dynamics of a 2-niche sympatric system. A computer screenshot displays a phase-portrait solution of a 2-mating-bias-allele, 2-niche sympatric ecosystem without viable hybrids, as the value of a search-cost variable *θ* is varied. This example uses the same parametric values as those in Fig 9, except for the value of the new variable *θ*, and an *Ax*/*Bx* interval of 0.05. If *θ* were 1, there would be no search cost and the phase portrait would appear identical to that in Fig 9. If *θ* were 0, there would be no matching encounters among individuals and the population would go extinct. When the value of *θ* is decreased from 1 to 0.8, population vectors near the ends of the diagonal line are driven toward the origin and its opposite diagonal corner, which makes invasion by high-mating-bias alleles more difficult.

Therefore, increased search cost (*θ*) can make invasion by a high-mating-bias mutant allele more difficult in a single-mating-bias-allele sympatric population undergoing ecological disruptive selection. This effect can be mitigated by increasing the invading population ratio *NAx*, decreasing the value of *α* (increasing the mating bias of the mutant allele), or increasing the value of *n* (increasing the number of matching rounds). In the example shown in Fig 19, if *θ* = 0.9, an initial invading mutant population *NAx* = 0.0001 (*NBx* = 0) cannot invade even if we increase the number of matching rounds *n* to 100. However, for the same value of *θ*, if we increase *NAx* to 0.001 (*NBx* = 0), invasion becomes successful for values of *n* ≥ 9. For *NAx* = 0.01 (*NBx* = 0), the mutant *X* allele can also invade with values of *n* ≥ 6.

## V. Mathematical Model of Two Mating-Bias Alleles With Viable Hybrids

Analysis of the mathematical model without viable hybrids can be extended to investigate how the presence of viable hybrids affects the nonlinear population dynamics in our 2-mating-bias-allele, 2-niche sympatric ecosystem. The presence of viable hybrids in the mating pool should weaken the effect of *f* and thus the strength of ecological selection on *NA* and *NB*. This is because the mating of *NA* and *NB* ecotypes with hybrids, as well as the mating of hybrids among themselves, can produce additional *NA* and *NB* offspring that add to their respective groups and increase the effective value of *f* for each niche group. Using similar mathematical derivations as we did for the system without viable hybrids, the nonlinear dynamics of a 2-mating-bias-allele, 2-niche sympatric system with viable hybrids can be described by the following set of difference equations (see Appendix for detailed mathematical derivation):

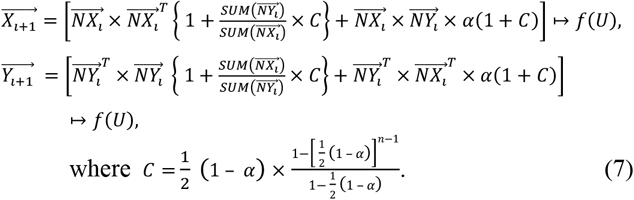

For the special case when 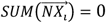 or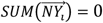, i.e., when the *X* or the *Y* allele is absent:

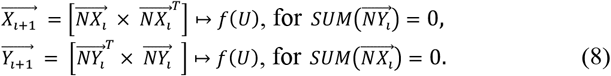

The vectors 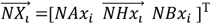 and 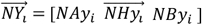 specify the normalized population ratios at the start of generation *i*. They represent offspring genotypes that have survived ecological and sexual selection from the previous (*i* − 1) generation. *NAx*_*i*_ is the ratio of *X* allele in niche *A, NBx*_*i*_ is the ratio of *X* allele in niche *B*, and 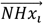 is a vector that specifies the normalized ratios of *X* allele in the hybrid niches. In our 3-ecological-locus model, 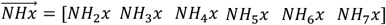, which specifies the *X*-allele ratios in the six hybrid niches (out of a total of eight possible niches from permutations of the three ecological loci). Similarly, *NAy*_*i*_, *NBy*_*i*_, and 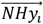 specify the ratios of the *Y* allele in their respective niches. The sum of all the elements in 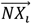 and 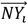 is equal to 1, i.e.,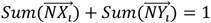. The probabilities of encounters among the various groups can be represented by elements of a matching probability matrix *M* shown in Fig 20. In the equations, *U* is a unit offspring matrix analogous to the one used in the model without viable hybrids (see Fig 8). Except in this case, because there are viable hybrids, *U* also needs to include offspring compositions for matches that include hybrid parents. The operation *ΣM* ↦ *f*(*U*) has been defined previously in the system without viable hybrids. It performs element-wise multiplication of the elements of a matrix *ΣM* (which specifies the accumulated probability ratios of different types of parental matches) by the offspring proportions in each cell of *U* to produce different offspring types.

**Fig 20.**
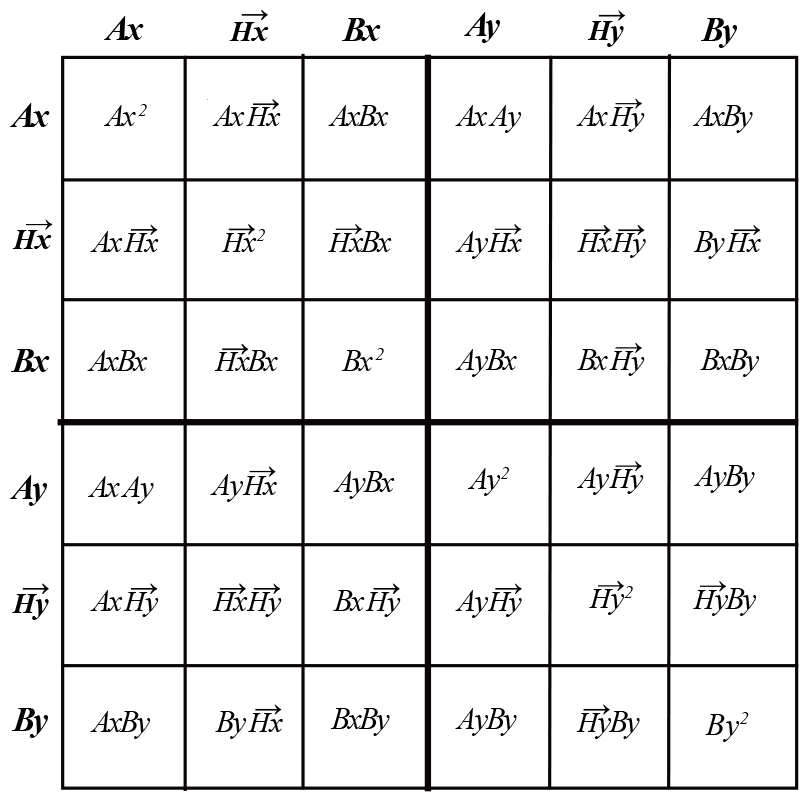
Matching probability matrix *M* for a sympatric model with viable hybrids. 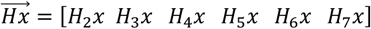 and 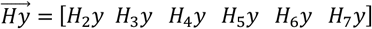 specify the ratios of *X* and *Y* mating-bias alleles in the hybrid niches. *Ax, Ax, Bx*, and *By* specify the ratios of mating-bias alleles, *X* and *Y*, in the *A* and *B* niches. The cells of the matrix show the products of their respective variables on the horizontal and vertical headings.

If *off* (the number of offspring that a matched individual can produce) is sufficiently large so that enough offspring are produced in each generation to completely saturate the carrying capacities of all the niches, *NAmax*, 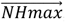, and *NBmax*, then the normalized *NAx*, 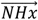, *NAy, NBx*, 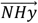, and *NBy* groups going into the next generation can be calculated as follows:

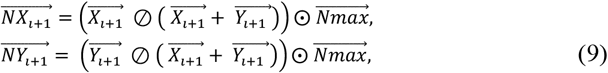

Where 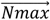 is a vector with elements that are the normalized ratios of all niche carrying capacities: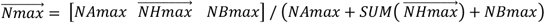. The Hadamard operators ⨀ and ⊘ perform element-wise multiplication and division of two same-sized vectors or matrices.

A computer program was developed to numerically solve these difference equations (7)–(9) for a 3-ecological-locus, 2-mating-bias-allele, 2-niche, sympatric ecosystem with viable hybrids and plot their phase-portrait solutions (Fig 21). Here, in additional to niches *A* and *B*, we also have six hybrid niches *H*_2_ to *H*_7_, defined by the genetic proximity of their ecotypes to those in niches *A* and *B*. If ecotype *A* has genotype *aaa* at its three ecological loci, and ecotype *B* has *bbb*, then the ecological genotypes of the hybrids are *H*_2_ = *aab, H*_3_ = *baa, H*_4_ = *aba, H*_5_ =*bab, H*_6_ =*abb*, and *H*_7_ =*bba*. 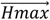 specifies the carrying capacities of the hybrid niches, and if they are not all zero, then the system can produce viable hybrid offspring that participate in the next mating generation. Similar to Fig 9, Fig 21 shows a vector-field phase portrait of the 3-ecological-locus, 2-mating-bias-allele ecosystem with viable hybrids, according to various input parameters. In addition to the usual inputs, *Hx* specifies the beginning ratio of *X* allele in a hybrid niche, so the beginning ratio of *Y* allele in the same niche is by default 1 − *Hx*. When a hybrid niche is empty, i.e., when *Hx* = *Hy* = 0, its *Hx* value is set to be equal to −1. *Hfx* specifies the final *X* allele ratio in a hybrid niche after *ii* generations if a fixed point exists.

**Fig 21.**
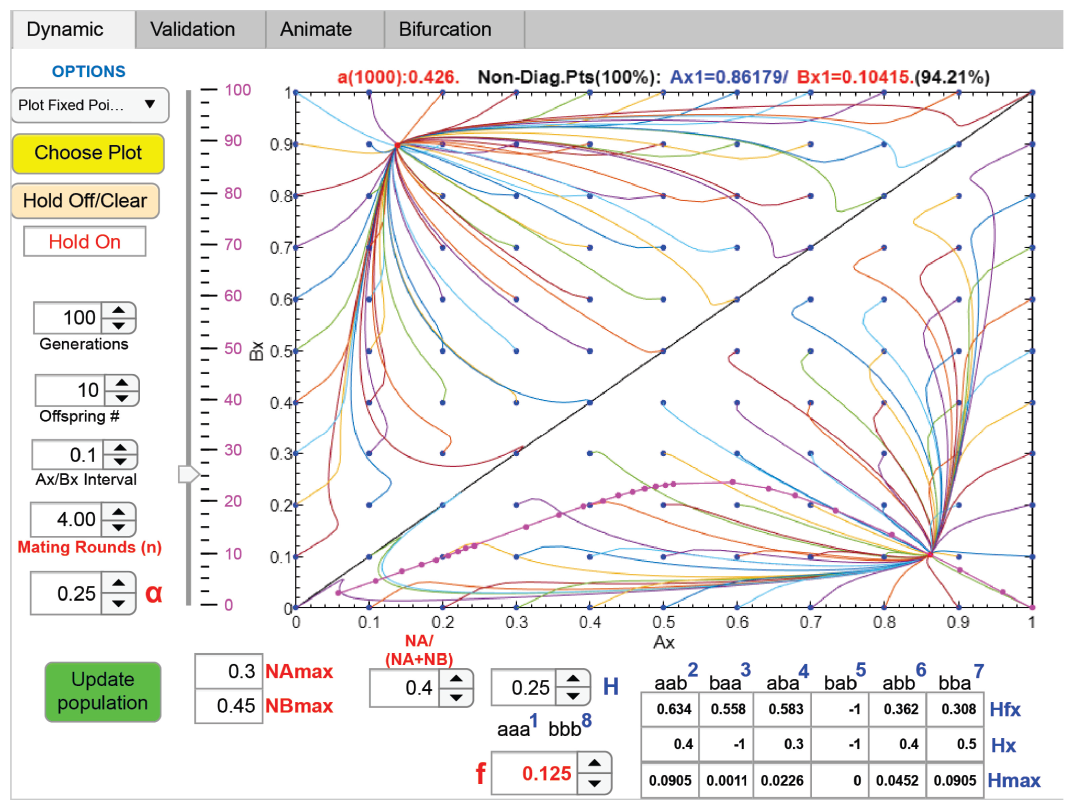
Phase portrait solution of a 2-niche sympatric system with viable hybrids. A computer screenshot shows a phase-portrait solution of the difference equations derived from a 2-niche model with viable hybrids. It uses and displays the same parametric variables as those for the model without viable hybrids (Fig 9). However, this model includes hybrid variables *H*, which represents the ratio of hybrid niches in the ecoscape (i.e., the sum of 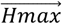, the hybrid niche carrying capacities), and *Hx*, the ratio of *X* mating-bias allele in each hybrid niche. The ratio of the *Y* allele in each hybrid niche is *Hy* = 1 − *Hx*, except in the case when *Hx* is expressed as −1, which is used to signify *Hx* = 0 and *Hy* = 0. The variable *Hfx* represents the final *X* allele ratio in each hybrid niche when the system reaches dynamic equilibrium at a fixed point. Also shown in the graph is a fixed-point curve (magenta line) that traces all the fixed points in the vector field as the variable *α* is varied.

Fig 21 shows the nonlinear system with viable hybrids converging to a fixed point. Also plotted is a fixed-point trajectory curve for the parameter *α*. In general, the computation results are in line with our expectation that the presence of viable hybrids in the mating pool will weaken ecological selection and increase the effective value of *f*, as compared to the model without viable hybrids. In contrast to what has been observed in the model without viable hybrids, many of the vector lines in Fig 21 make sharp-corner turns and cross over each other. This likely occurs because we are projecting an eight-dimensional plot with eight independent parameters (*Ax, H*_2_*x, H*_3_*x, H*_4_*x, H*_5_*x, H*_6_*x, H*_7_*x, Bx*) onto a two-dimensional surface (defined by *Ax* and *Bx*). At the crossing points of vector lines, although their *Ax* and *Bx* values are the same, the 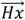 values of the crossing vector lines are different. Besides these differences, the nonlinear dynamic behaviors of the two models, with and without viable hybrids, are qualitatively similar, especially when *Hmax* is small (i.e., when there is strong ecological selection).

Fig 22 shows a schematic diagram that can be used to visualize how the presence of viable hybrids affects the population dynamics at niche *A* and niche *B*. Let *Ax*, 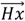, *Bx, Ay*, 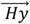, and *By* represent normalized population ratios in the system at the beginning of each mating generation. Let 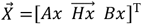 and 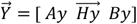, so that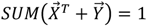. *NAx, NAy, NBx*, and *NBy* are the original normalized population ratios of our model in Fig 5. In Fig 22, the 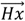 and 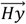 populations are represented by two sidebars in the graph, and they are arranged in order of their ecological genotype proximity to the ecotypes in *NA* or *NB*. For instance, in our 3-ecological-locus example, *H*_2_*x* and *H*_3_*x* (genotypes *aab* and *baa*) are positioned closer to the bottom of the sidebar than *H*_4_ *x* and *H*_5_ *x* (genotypes *aba* and *bab*) because of their closer genotype proximity to *NA* (genotype *aaa*). The gray scale shading in the sidebars gives a qualitative indication of the relative population ratios in the hybrid niches.

**Fig 22.**
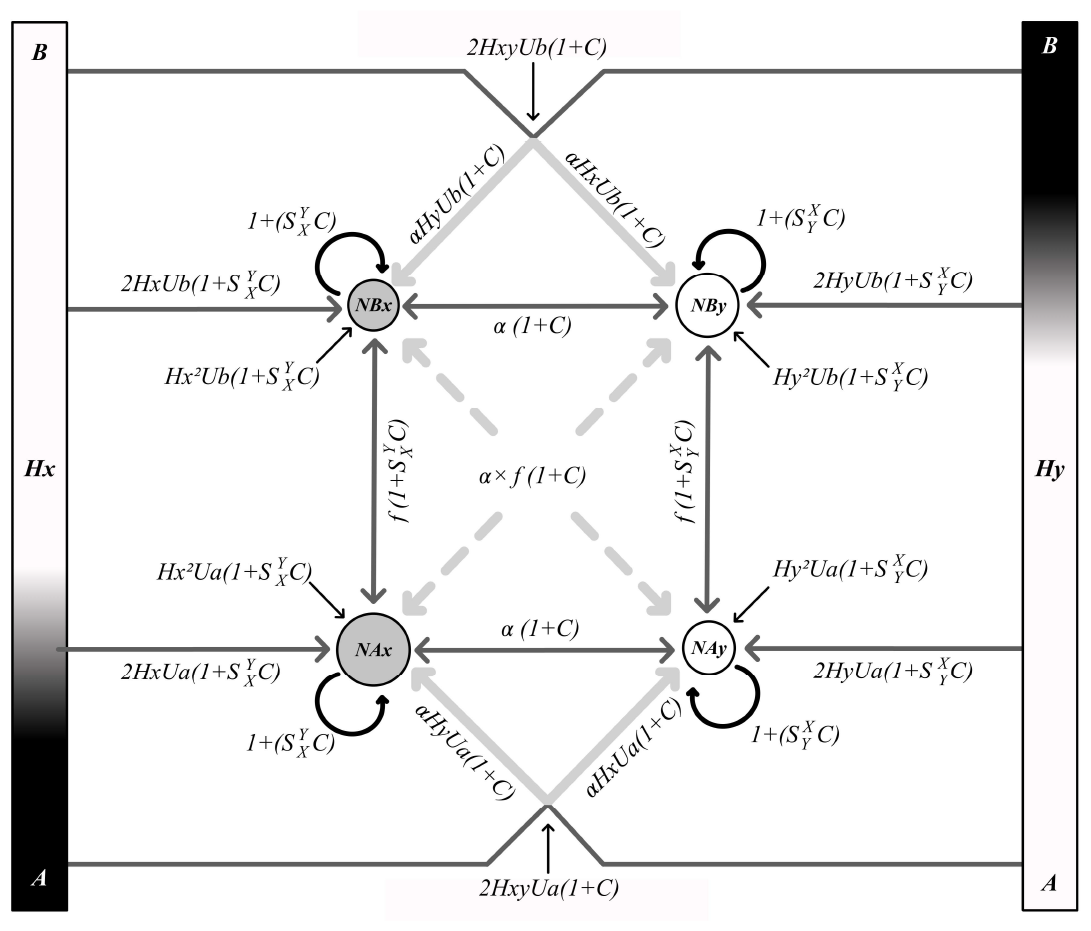
The effect of viable hybrids on a 2-niche mathematical model. A schematic diagram illustrates how the presence of viable hybrids affects the interactions among niche populations in a mathematical model of a 2-mating-bias-allele, 2-ecological-niche sympatric ecosystem. The two sidebars represent the populations of *X* and *Y* alleles in the hybrid niches, arranged in order of their genotype proximity to the genotypes in the *A* and *B* niches. The sidebars are shaded to indicate the population density distribution in the hybrid niches. The solid arrows pointing from the sidebars to the center niche groups, along with their associated multipliers, depict how the various hybrid groups can affect the respective niche *A* and *B* populations. The gray arrows signify that the unit offspring produced from the hybrid matches are evenly distributed among the specified *A* and *B* niche groups, because the matched parents have different mating-bias alleles.

Also shown in the diagram are various multipliers that supply additional offspring to the *NAx, NAy, NBx*, and *NBy* niche groups due to the presence of viable hybrids. For example, the multiplier 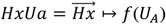 specifies the additional unit offspring that *NAx* receives because of mating with the hybrid group 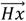. Similarly, *Hx*^2^*Ua* specifies the additional unit offspring that *NAx* gains because of the presence of matched mating pairs in 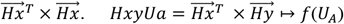 specifies the additional unit offspring that are contributed to groups *NAx* and *NAy* because of the presence of matched mating pairs in 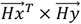. In the diagram, 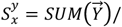 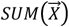 and 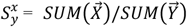. Therefore, the interaction between *NAx* and the multiplier 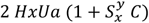 supplies an additional 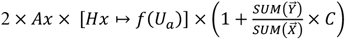 unit-offspring ratio to the population *NAx* after a mating generation (see Appendix for detailed derivations).

If the viable hybrid populations are small, then the influences of multipliers *Hx*^2^, *Hy*^2^, *Hxy*, 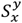, and 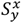 are relatively minor. Likewise, offspring produced by the mating of niche *A* or niche *B* groups with hybrids that carry different mating-bias alleles— e.g., *α* × (*Ax* × *HyUa*)(1 + *C*) and *α* × (*Ay* × *HxUa*)(1 + *C*)—are distributed equally to the two different mating-bias-allele groups in the same niche (e.g., *NAx* and *NAy*) and do not contribute much to the difference between the groups. Therefore, the major players in the model are the multipliers 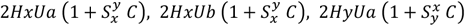, and 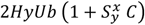, and they have the largest impact on the niche *A* and *B* population ratios. Consequently, to achieve maximum reproductive isolation, it is desirable for the *Hx* population to congregate near niches closest to the *NA* genotype (near the bottom of the *Hx* sidebar), and the *Hy* population, closest to the *NB* genotype (near the top of the *Hy* sidebar).

## IV. Mathematical Model of Three Mating-Bias Alleles

Employing computer algorithms similar to those used in the 2-mating-bias-allele case, computer programs were developed to simulate and numerically solve the nonlinear dynamic behavior of a 3-mating-bias-allele system in a 2-niche sympatric ecoscape. (See Appendix for a description of the algorithms.) To model the 3-mating-bias-allele system, we used the same variables and nomenclatures as those used in the 2-mating-bias-allele case. However, in addition to the two mating-bias alleles, *X* and *Y*, we specified a third mating-bias allele, *Z*, with variables *α, β*, and *γ* representing the matching probability ratios among the three mating-bias alleles. Fig 23 shows the matching compatibility tables for the three mating-bias alleles.

**Fig 23.**
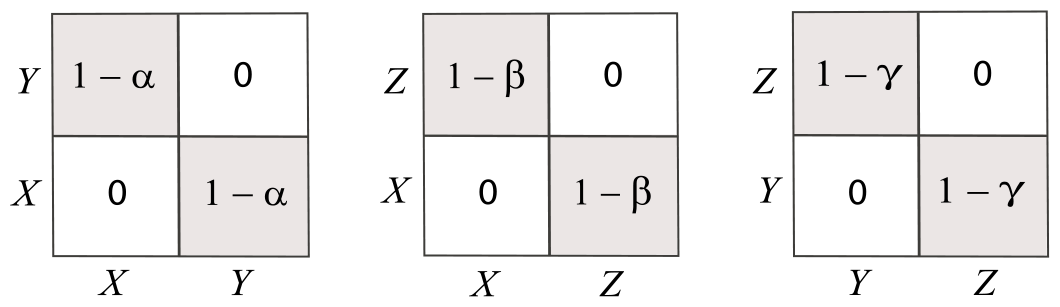
Matching compatibility tables for mating-bias alleles, *X, Y,* and *Z*. When two individuals possess the same mating-bias alleles, the mating barrier between them is zero, i.e., they are perfectly compatible. When they possess different mating-bias alleles, their compatibility is determined by variables *α, β*, and *γ*.

Fig 24 shows a representative screenshot that displays the nonlinear dynamic behavior of our 3-mating-bias-allele, 2-niche ecosystem without viable hybrids. Another program that models the same ecosystem with viable hybrids also yielded similar expected results (not shown). Let *Ax* be the ratio of *X* allele in niche *A, Ay* be the ratio of *Y* allele in niche *A*, and *Az* be the ratio of *Z* allele in niche *A*, and define similarly for *Bx, By*, and *Bz*, so that *Ax* + *Ay* + *Az* = 1 and *Bx* + *By* + *Bz* = 1. The three plots in Fig 24 show the trajectories of different lineages of populations starting from all possible permutations of *Ax, Ay, Az, Bx, By*, and *Bz* at intervals of 0.2. (The starting values are shown as blue dots, and the ending values after 1000 generations are shown as red and magenta dots.) Displayed at the bottom of the figure are parametric values for variables *NA, NB, f*, and *n*, which are the same variables as those in the 2-mating-bias-allele case, as well as the values for *α, β*, and *γ*. In this example, the computer algorithms calculated four fixed points based on these parametric values. The fixed-point values are color-coded and displayed above the plots. The trajectories that converge to these fixed points are also displayed in the same corresponding colors. Only trajectories that converge to fixed points are displayed. Divergent trajectories that end up on the diagonal line are not displayed, except for their endpoints, which are shown as magenta dots on the diagonal line.

**Fig 24.**
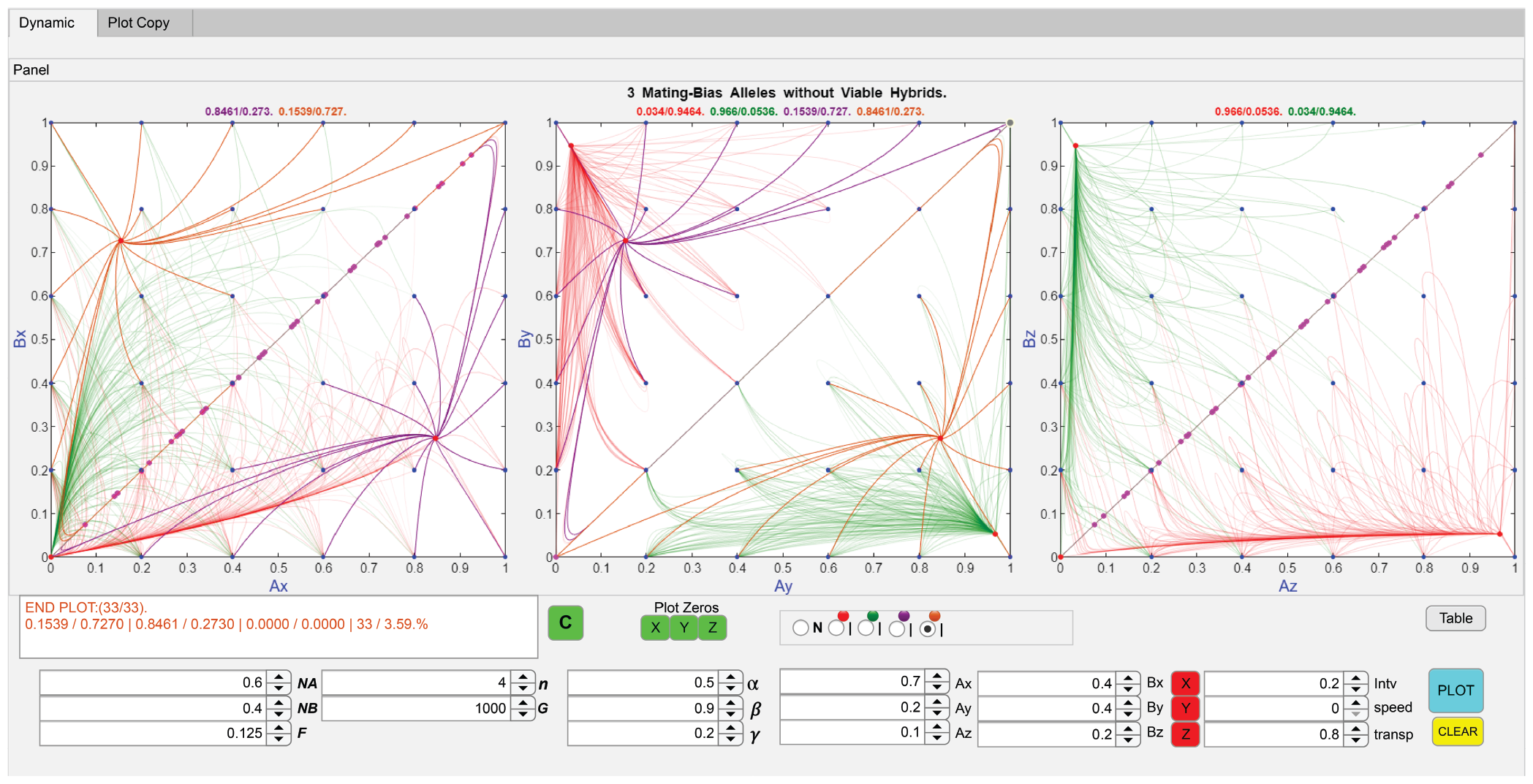
A computer screenshot showing the phase-portrait solutions of a 3-mating-bias-allele, 2-niche sympatric ecosystem. The three plots show the ratios of mating-bias alleles *X, Y, Z* in niche *A* and niche *B*. The lineages that converge to different fixed points are identified by different colored lines. The fixed-point values are displayed above the plots. The parametric values used in the calculations are displayed at the bottom of the figure.

Numerical computation using different parametric values revealed that it is impossible to have more than two mating-bias alleles coexist in fixed-point polymorphism in a niche. One of the three mating-bias alleles inevitably becomes extinct first, resulting in either *Ax* = *Bx* = 0, *Ay* = *By* = 0, or *Ax* = *Bz* = 0. The remaining two mating-bias alleles then interact according to the nonlinear dynamics described by the 2-mating-bias-allele model, and the system either converges to fixed-point polymorphism, partially converges, or diverges (Fig 9, 11, and 12). In Fig 24, when the *X* allele is eliminated after 1000 generations, the remaining *Y* and *Z* alleles converge to fixed-point polymorphism in the two niches, as shown by the red and green trajectories. When the *Y* allele is eliminated, the trajectories (not shown) of the *X* and *Z* alleles diverge and end up on the diagonal line. When the *Z* allele is eliminated, the brown and purple trajectories map out the fixed points created by alleles *X* and *Y*. Because not all three mating-bias alleles can coexist in fixed-point polymorphism in a given niche, the 3-mating-bias-allele, 2-niche system seems to exhibit the phenomenon of competitive niche exclusion—in other words, disruptive ecological selection results in hybrid loss which creates a new niche for suitable pairs of mating-bias alleles to explore and eliminate the less-fit variants.

## VII. Invasion by Mutants

In a 2-mating-bias-allele, 2-niche sympatric system, if the parametric values are such that the system becomes convergent or partially convergent (Fig 9 and 12) and the convergent trajectories involve the origin (i.e., at *Ax* = *Bx* = 0), then a mutant allele *X* with the specified mating-bias value *α* will be under selective advantage to invade a population that consists entirely of *Y* alleles and be swept into fixed-point polymorphism with the *Y* allele. Assuming an infinite population size, the lower the mating-bias value *α* and the smaller the normalized niche ratio *NA*, the faster such an invasion will take place (Fig 18).

If the mating-bias value *α* in the 2-mating-bias-allele model is controlled by a meta-allele at a separate genetic locus, then a mutant meta-allele producing a lower value of *α* (a higher mating bias) can invade a 2-mating-bias-allele, 2-niche system with fixed-point polymorphism engendered by a meta-allele associated with a higher value of *α* (a lower mating bias). Fig 25 displays the result of a computer simulation that shows a high-mating-bias meta-allele invading and completely replacing a low-mating-bias meta-allele when both variants of the meta-allele can produce fixed-point polymorphism. This occurs because a lower value of *α* increases the reproductive isolation, reduces nonviable hybrid loss, and is beneficial for individuals in both niches. Mutant meta-alleles producing higher values of *α* (*α* > 0.5 in the Fig 25 example) will not be able to invade such a system. This process can be considered a “one-allele model” of incremental RI because such a high-mating-bias mutant meta-allele is favored by individuals in both niches that possess either the *X* or the *Y* mating-bias allele, and only a single allele change is needed to increase RI.

**Fig 25.**
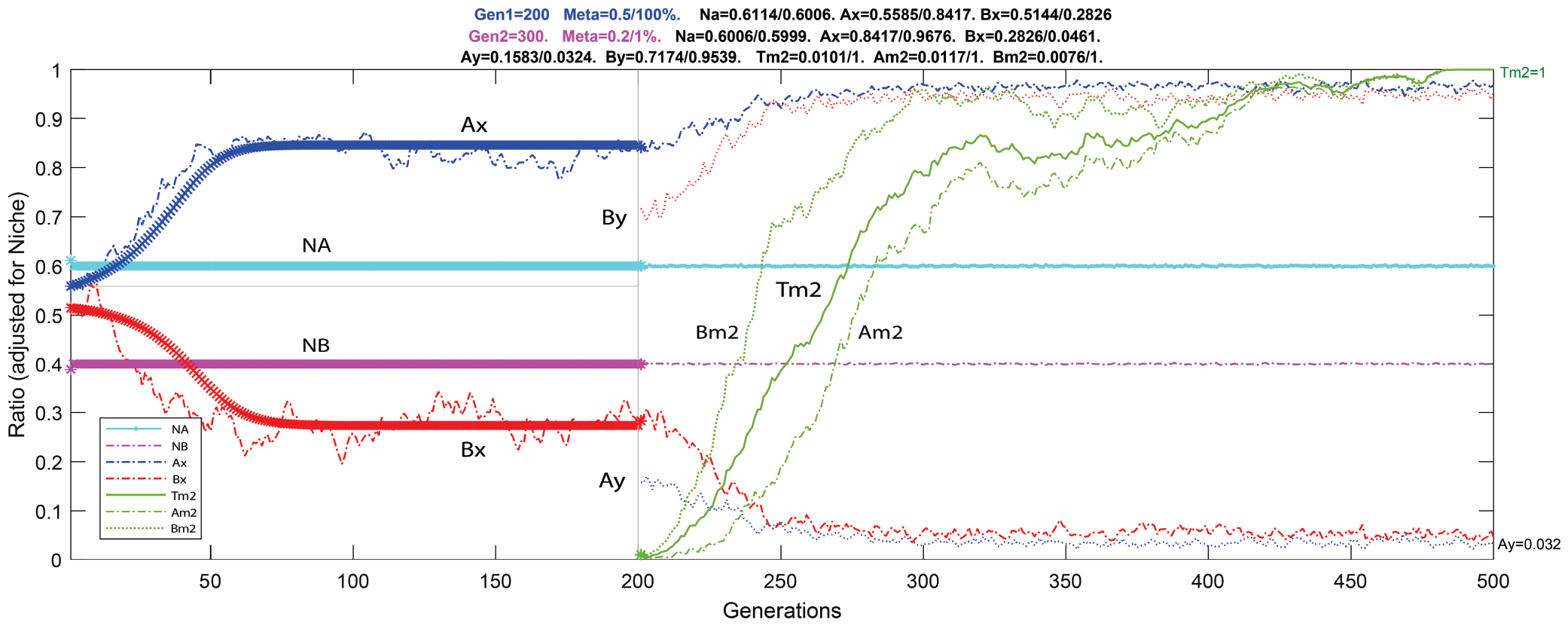
Invasion by a meta-allele in a sympatric system. The plotted results of a computer simulation show that a mutant meta-allele with a higher mating bias can invade and replace an existing meta-allele with a lower mating bias in a 2-mating-bias-allele, 2-niche sympatric system with fixed-point polymorphism. The system initially had a meta-allele with *α* = 0.5. It was invaded by a meta-allele with a higher mating bias *α* = 0.2. The simulation ran for 500 generations. At generation 200, the higher-mating-bias meta-allele was introduced at a ratio of 0.01 of the total population. *Tm*2 is the ratio of all mutant meta-alleles in the population, *Am*2 is the ratio of mutant meta-alleles in niche *A*, and *Bm*2 is the ratio of mutant meta-alleles in niche *B*. The solid lines before generation 200 are the calculated results, and the dotted lines are the results from the computer simulation. The rest of the variables are the same as those used in the 2-mating-bias-allele model. The parametric values and the numerical results of the simulation are displayed above the graph. The system assumes no viable hybrids. *n* = 4. *f* = 0.125. *Off* = 10. Simulation population size = 3000.

The situation becomes less clear when a mutant meta-allele associated with high mating bias (low *α*) capable of producing fixed-point polymorphism tries to invade a divergent system caused by a meta-allele with low mating bias (high *α*)—in other words, the invading mutant tries to convert a divergent system without fixed points into a convergent system with fixed points. In general, it is more difficult for a rare mutant meta-allele to invade a divergent system than a convergent system and produce fixed-point polymorphism. Our computer simulation has shown that for this to happen, the invading meta-allele needs to produce higher mating bias (lower values of *α*) and have a larger initial invading population ratio than those required to invade a comparable convergent system (see discussion on invasion thresholds.). This scenario is more likely to occur via gene flow in parapatric speciation or in secondary contact following allopatric or peripatric divergence.

In a divergent system, sexual selection is the dominant selection force that favors low-mating-bias variants, and it tends to drive the population toward homogeneity, with just a single type of mating-bias allele remaining, given long enough time. (For example, in Fig 11, the points on the diagonal line will move faster toward the line’s endpoints given lower values of *α* and *n*.) Initially, sexual selection in a divergent system favors the invasion of a lower-mating-bias meta-allele, one that produces an *α* greater than the existing *α* in the system. However, when there is only a single type of mating-bias allele left in the population, the matching barrier no longer exists, and meta-alleles no longer have an effect on the mating-bias alleles and cannot invade.

Next, we examined how a 2-mating-bias-allele, 2-niche sympatric system could be invaded by a third mutant mating-bias allele. Fig 26 shows a representative example. Initially, the system only had two mating-bias alleles, *X* and *Y*, with an *α* value of 0.5. The system was allowed to converge and reach fixed-point polymorphism, until at generation 500, when a third mating-bias allele *Z* with mating-bias values *β* = 0.9, *γ* = 0.2, and initial population ratios *Az* = 0.01 and *Bz* = 0 was able to invade the system and completely eliminate allele *X*. After that, allele *Z* and allele *Y* reached a new fixed-point polymorphism that had a higher mating bias *γ* and resulted in greater reproductive isolation between the two niches. This process can be considered a “two-allele model” of incremental RI because divergent species need to fix different mating-bias alleles to increase RI.

**Fig 26.**
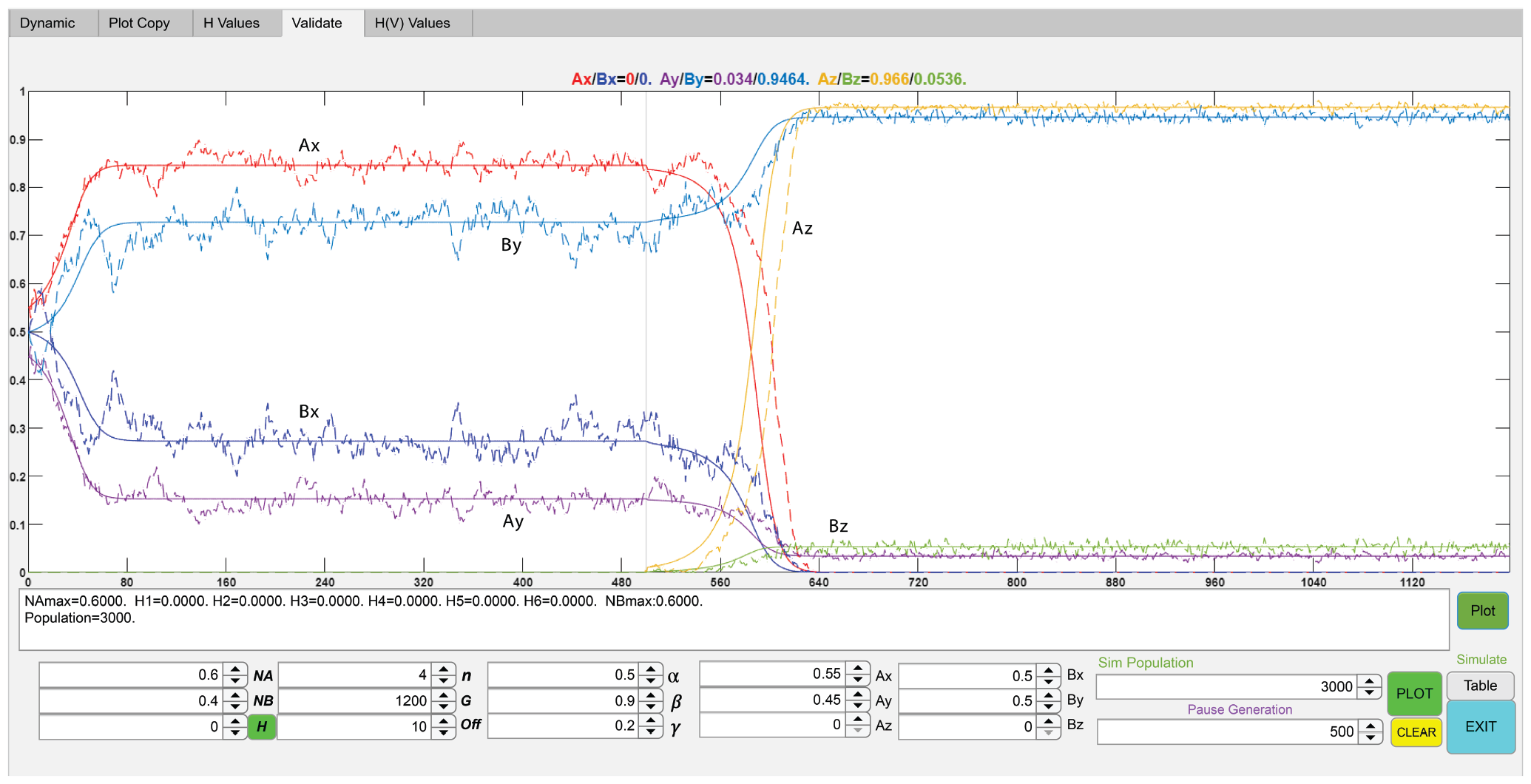
Invasion by a third mating-bias allele in a sympatric system. A labeled computer screenshot shows how a 2-mating-bias-allele sympatric system in stable fixed-point polymorphism can be invaded by a third mating-bias allele to achieve increased reproductive isolation. Initially, a 2-niche sympatric system with two mating-bias alleles *X* and *Y* and a mating-bias value *α* = 0.5 was allowed to converge to a fixed point at *Ax* = 0.8461 and *Bx* = 0.273. At generation 500, a third mating-bias allele *Z* with mating-bias values *β* = 0.9 and *γ* = 0.2 was introduced in niche *A* at a population ratio of 0.01 (1%), i.e., *Az* = 0.01 and *Bz* = 0. The mutant *Z* allele was able to invade and eliminate the *X* allele by generation 640 and drive the system toward a new fixed point at *Ay* = 0.034 and *Az* = 0.966, resulting in increased reproduction isolation between the two niche populations. The color-coded new fixed-point values are shown above the plot. The input parametric values are displayed at the bottom of the figure. The solid lines are the calculated results of mathematically derived computer algorithms. The dotted lines are the results of an individual-based computer simulation using a population size of 3000.

How such an invasion can occur is explained by analyzing the system dynamics in Fig 24, which uses the same input parameters as those used in Fig 26 and displays the phase-portrait results for all initial values of *Ax, Ay, Az, Bx, By*, and *Bz*. Initially, when *Az* = *Bz* = 0, the system has two fixed points at *Ax*/*Bx* = 0.8461/0.273 and *Ax*/*Bx* = 0.1539/0.727, marked by the purple and brown trajectories in the plots. After the invasion by the *Z* allele, the new fixed points at *Az*/*Bz* = 0.9666/0.0536 and *Az*/*Bz* = 0.034/0.9464 are marked by the red and green vector trajectories respectively. Because the origin of the *Az*-*Bz* plot (i.e., at *Az* = *Bz* = 0) is included in the areas of starting *Az*/*Bz* values of the red and green trajectories, and because the initial fixed points in the *Ax*-*Bx* and *Ay*-*By* plots (marked by the purple and brown trajectories) are also within the areas delineated by the starting points of the red and green trajectories, a small mutant population of the *Z* allele will be driven by the system’s nonlinear dynamics away from the origin and toward the converging fixed points in the *Az*-*Bz* vector field. Concurrently, populations at the initial fixed points in the *Ay*-*By* plot are driven to their new, higher-mating-bias fixed points marked by the red and green trajectories; and populations at the initial fixed points in the *Ax*-*Bx* plot are driven toward the origin (extinction).

It is easy for the *Z* allele to invade and replace the *X* allele because *β* is high (i.e., there is a low mating barrier between the *X* and *Z* alleles) and *γ* is lower than *α* (i.e., there is a higher mating barrier between the *Z* and *Y* alleles that confers greater selective advantage by minimizing hybrid loss). In the limit when *β* = 1, an invading *Z* allele will just act like a mutant *X* allele with a higher mating-bias value *γ*. However, if we keep the rest of the parametric values the same, and gradually decrease the value of *β*, for *β* < 0.7748, the invasion will fail, and the *Z* allele will be eliminated instead. Similarly, in the example in Fig 26, if we change just one variable at a time and keep the rest of the input parameters the same, the *Z* mutant allele in niche *A* will also fail to invade and will be eliminated if *NA*< 0.24, *γ* > 0.3917, or *n* < 3. Otherwise, in our computer calculations, the *Z* allele can invade with a starting population *Az* as low as 1× 10^−15^, which is probably the accuracy limit of our program’s numerical approximation. We refer to these limiting parametric values as the “invasion threshold,” which represent the initial barrier that a population of mutant alleles must overcome for a successful invasion.

Similar to the meta-allele case, it is harder for a mutant third allele with a higher mating bias to invade a divergent 2-mating-bias-allele system and convert it into a convergent system with fixed-point polymorphism. Fig 27 shows such an example. Even with a *β* value of 1 (i.e., the *X* and *Z* alleles are perfectly compatible) and *γ* = 0.2, a mutant *Z* allele with an initial population ratio of *Az* = 0.1 (*Bz* = 0) cannot invade a divergent system with two mating-bias alleles *X, Y* and *α* = 0.7, and the *Z* allele is eliminated. However, if the initial population ratio of the *Z* allele is increased to *Az* = 0.1343 or greater, then invasion becomes successful, the *X* allele is eliminated, and the system becomes convergent with fixed points at *Az*/*Bz* = 0.966/0.0536 and *Ay*/*By* = 0.034/0.9464. If we change just one variable and keep the rest of the parametric values the same in Fig 27, the *Z* allele will invade if *γ* ≤ 0.064 or *n* ≥ 5. The *Z* allele cannot invade with any value of *NA* or *β* with the rest of the given parametric values unchanged. If we increase *n* (reduce the cost of assortative mating), the initial *Az* population ratio that is required for a successful invasion can be reduced to a certain degree. For instance, for *n* = 10, invasion is successful for an initial *Az* population ratio ≥ 0.0667; and for both *n* = 25 and *n* = 100, invasion is successful for an initial *Az* population ratio ≥ 0.0658. Note that if the pre-invasion system only contains a single mating-bias allele *Y* (i.e., *Ax* = *Bx* = 0), a mutant *Z* allele with *γ* = 0.2 can invade and establish fixed-point polymorphism with an initial *Az* population ratio as low as 1× 10^−15^ (e.g., the invasion scenario in Fig 9). Therefore, the presence of an *Ax* population in our divergent system seems to act to increase the invasion threshold of a compatible mutant *Z* allele with high *β* and low *γ* values that are capable of making the system convergent.

**Fig 27.**
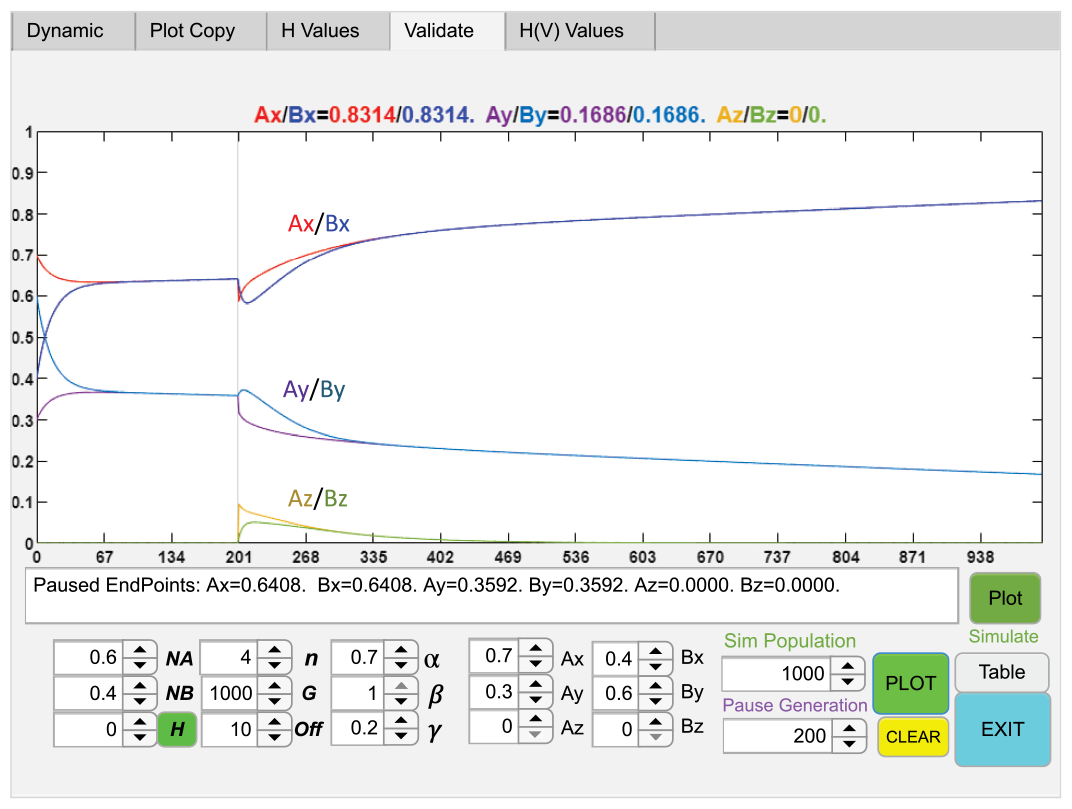
A labeled computer screenshot showing the existence of invasion thresholds. A mutant allele *Z* with mating-bias values *β* = 1 and *γ* = 0.2 and an initial population ratio of 0.1 (10%) in niche *A*, i.e., *Az* = 0.1 and *Bz* = 0, was introduced at generation 200, but failed to invade a divergent system of 2-mating-bias alleles, *X, Y* with a mating-bias value *α* = 0.7. The color-coded final allele ratios are displayed above the plot. The parametric input values are displayed at the bottom of the figure.

## VIII. Alidation

An individual-based computer program was used to simulate the population dynamics of a 3-ecological-locus, 2-mating-bias-allele, 2-niche sympatric ecosystem to verify the accuracy and validity of our derived nonlinear equations (4)-(6) and (7)-(9). The results are shown in Fig 28. The solid lines show the changes in *Ax, Bx, NA, NB*, and *NH* values obtained by solving, using numerical methods, the set of nonlinear difference equations that describe the mathematical model with viable hybrids, with the specified input parameters. The dashed lines are the results from the individual-based computer simulation using the same input parameters and with a starting population of 3000. The results from the numerical solution and the results from the individual-based simulation show a high level of correspondence.

**Fig 28.**
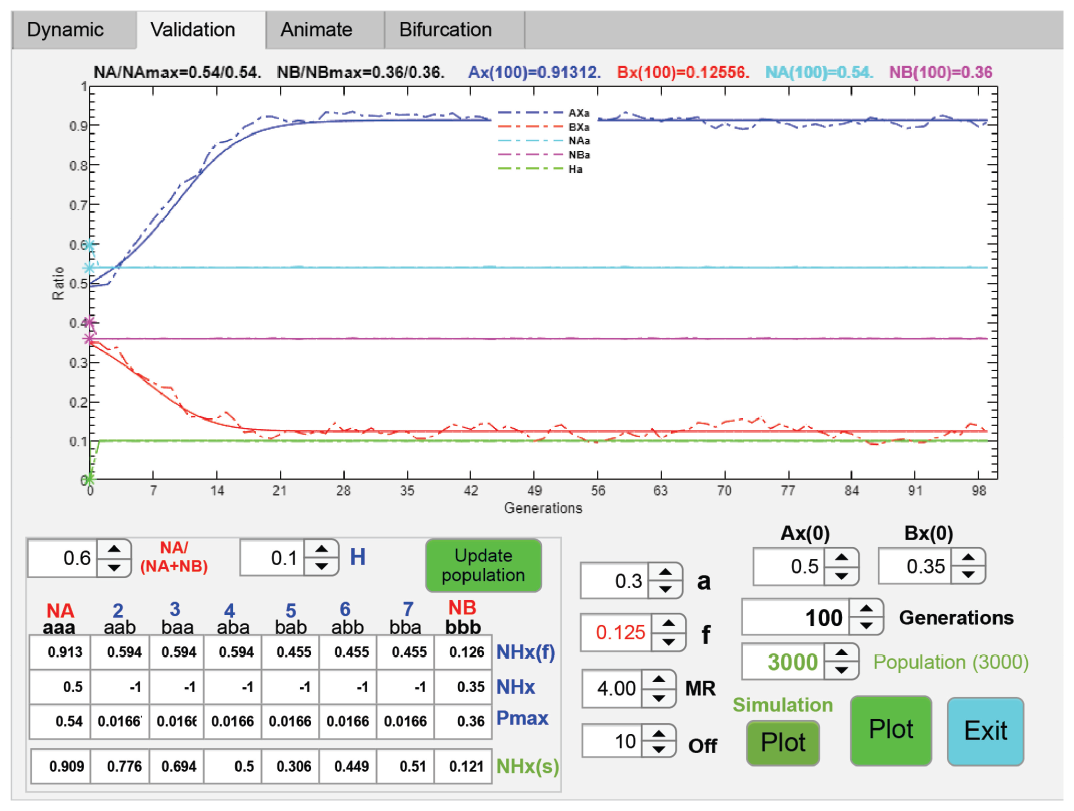
Validation of mathematical models with computer simulations. The graph in this computer screenshot plots the results of an individual-based computer simulation against the calculated solutions of its mathematical model’s difference equations. In this example, the mathematical model is a 2-mating-bias-allele, 2-niche sympatric system with viable hybrids. The parametric variables are defined in the same way as those used in Fig 21. The starting population for the computer simulation is 3000. The simulation uses individuals that have three ecological-allele loci and one mating-bias-allele locus. *Pmax* specifies a niche’s carrying capacity, and *NHx* specifies a niche’s starting *X*-allele ratio. *NHx*(*f*) is the calculated *X*-allele ratio of a niche at the fixed point in the mathematical model. *NHx*(*s*) is the *X*-allele ratio of a niche at the fixed point in the computer simulation. The results of the individual-based simulation are plotted as dotted lines in the graph, and the solutions of its mathematical model’s difference equations are plotted as solid lines. The two sets of results show a high level of correspondence.

A similar individual-based computer program was also used to simulate the population dynamics of a 3-ecological-locus, 3-mating-bias-allele, 2-niche sympatric system to verify the accuracy and validity of the computer algorithms used in Fig 24. The results also show a high level of correspondence (Fig 26).

## IX. The Effects of Contact Barriers Between Niches

Thus far, individuals in our sympatric systems could encounter one another freely and randomly to find compatible matching mates. Next, we extended our model to investigate how the existence of contact barriers between niches can affect the development of mating-bias-allele fixed-point polymorphism to produce premating RI. Such contact barriers could be the result of geographical barriers or habitat preferences that decreases the probability of matching encounters between ecotypes in different niches.

Fig 29 shows a representative example of the calculated phase-portrait solution of such a model. It uses the same input parameters as those in Fig 9 for a 2-mating-bias-allele, 2-niche sympatric system without viable hybrids. However, in this case, we have added a migration variable *m* that restricts matching encounters between niche *A* and *B* populations. The value of *m* can vary between 0 and 1. When *m* = 1, there is no contact barrier, and the system is purely sympatric. When *m* = 0, there is complete contact isolation, and we have two separated allopatric niche populations. Values of *m* between 0 and 1 allow for varying degrees of contact isolation between the niche populations and can be used to simulate different degrees of parapatric gene flow. In the matching probability matrix shown in Fig 7, variable *α* specifies the matching probability between population groups that have different mating-bias alleles. Similarly, variable *m* can specify the matching probability between groups that belong to different niches. In the model’s computations, the value of *m* only affects the result of the accumulated matching probability matrix *ΣM*_(*n*)_; otherwise, the rest of the algorithmic calculations remain the same.

**Fig 29.**
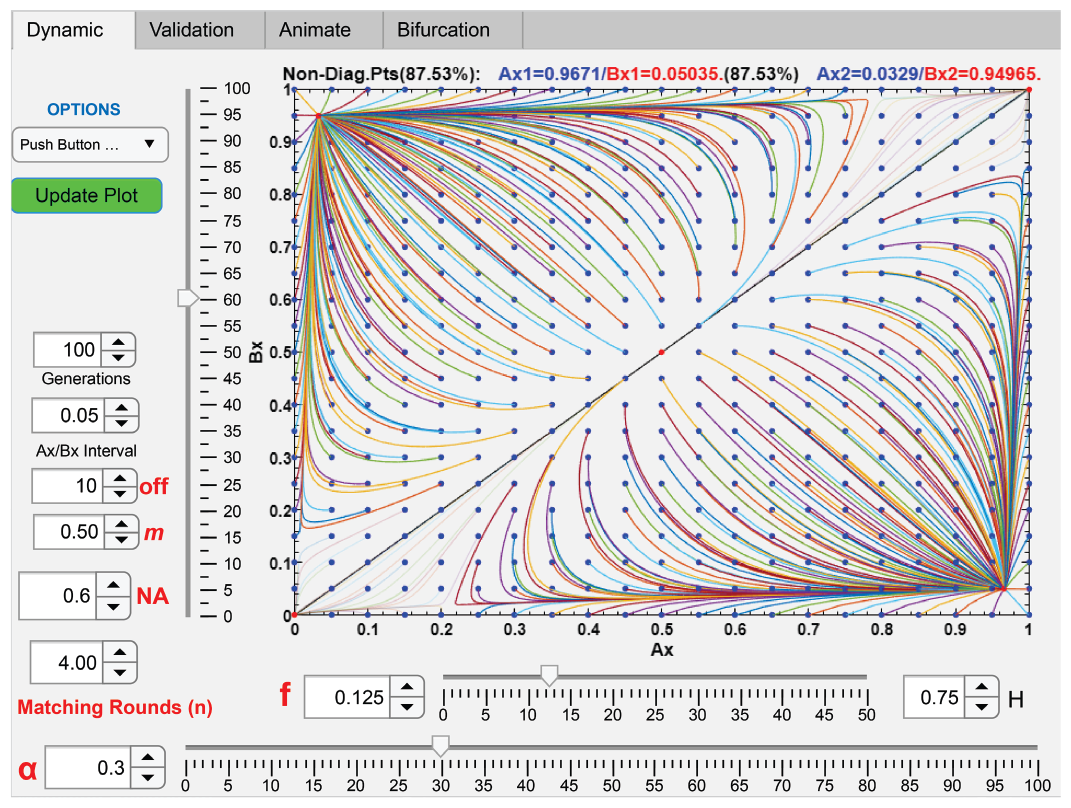
The effect of contact barriers in a 2-niche ecosystem. A computer screenshot shows the phase-portrait solution of a 2-mating-bias-allele, 2-niche ecosystem with no viable hybrids and with restricted contact between the two niche populations. *Ax*/*Bx* interval = 0.05. The system uses the same input parametric values as those in Fig 9, except a new migration variable *m* is added. When *m* = 1, there is no contact barrier between the two niches, and the system is sympatric. When *m* = 0, there is no contact between the two niches, gene flow is zero, and the system is allopatric. When compared to the sympatric system in Fig 9, Fig 29 shows that reducing the value of *m* to 0.5 moves the fixed points closer to the corners (*Ax* = 1, *Bx* = 0) and (*Ax* = 0, *Bx* = 1), resulting in increased premating RI. Meanwhile, sexual selection drives population vectors near the ends of the diagonal line toward the origin and its opposite diagonal corner (shown as faint lines), where only a single type of mating-bias allele remains in the system.

When *m* = 1, there is no contact restriction between the two niche populations, and we obtain the same vector-field phase portrait as in Fig 9. As we gradually reduce the value of *m* and increase the contact barrier, the fixed point in the lower vector field moves closer to the corner where *Ax* = 1 and *Bx* = 0, resulting in stronger premating RI, as shown in the example in Fig 29. Similarly, its mirror-image fixed point across the diagonal line in Fig 9 moves closer toward the corner where *Bx* = 1 and *Ax* = 0. Below a certain value of *m*, however, sexual selection in the separated niches begins to exert its influence, and this can drive niche populations with a single, predominant type of mating-bias allele toward mating-bias-allele homogeneity. This effect is shown in Fig 29. When *m* = 0.5, populations near the origin (where *Ax* = 0 and *Bx* = 0) of the vector field in Fig 29 are driven toward the origin, where only the *Y* alleles exist; and populations near the opposite corner, where *Ax* = 1 and *Bx* = 1, are also driven to that corner, where only the *X* alleles exist. When *m* = 0, niche *A* and niche *B* are completely isolated from each other, and the population dynamics in each niche are purely driven by sexual selection. This is shown in Fig 30. When *m* = 0, the more numerous type of mating-bias allele in each niche will completely eliminate the less numerous type of mating-bias allele in the same niche.

**Fig 30.**
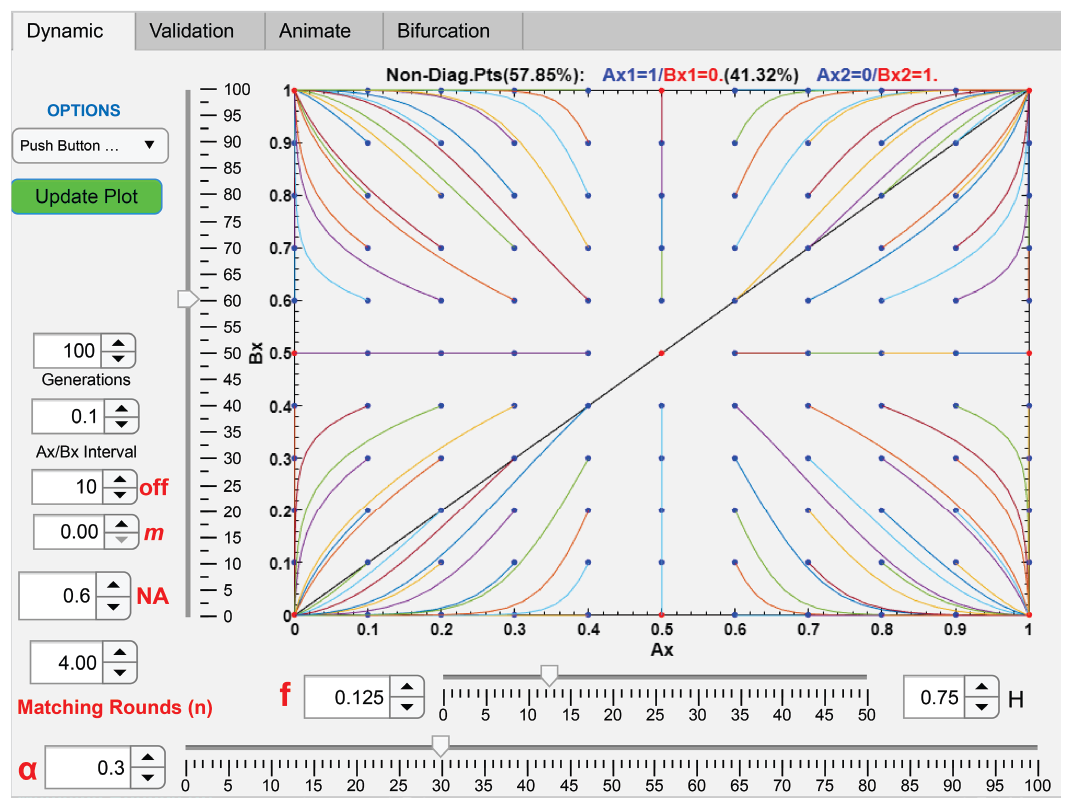
The effect of complete contact isolation in a 2-niche ecosystem. A computer screenshot shows the phase-portrait solution of a 2-mating-bias-allele, 2-niche ecosystem with no viable hybrids and with complete contact isolation between the two niches. The plot uses the same input parameters as those in Fig 29. However, the value of *m* has been changed to zero, resulting in complete contact isolation and no gene flow between niche *A* and niche *B* populations. In this case, sexual selection will drive the more numerous mating-bias allele in each niche to fixation by completely eliminating the less numerous mating-bias allele in the same niche.

These results suggest that when contact barriers first arise in a 2-niche sympatric population under disruptive ecological selection, the contact barriers can facilitate and strengthen premating RI by mating-bias-allele fixed-point polymorphism. However, as the contact barriers become greater, the effect of sexual selection in each niche increases, the percentage of fixed-point convergence in the phase portrait begins to shrink, and it becomes more difficult for a high-mating-bias mutant allele to invade a homogeneous system with only one type of mating-bias allele to produce fixed-point polymorphism.

## X. Mating-Bias Alleles with multiple gene loci

Using computer algorithms similar to those employed in the model with viable ecological hybrids (see Fig 21), we developed computer applications to investigate how mating-bias traits that are produced by the interactions of multiple gene loci could affect the emergence of premating RI in our 2-niche sympatric ecosystem.

Thus far, our models have used two mating-bias alleles, *X* and *Y*, that reside on a single gene locus. Now, assume *X* and *Y* are mating-bias traits or super-alleles that are the results of the interactions of three gene loci, and each gene locus has two alleles, *x* and *y*. Following the same approach used for categorizing ecological hybrids, we can arrange the eight permutations of the three gene loci based on their genetic proximity to one another: *h*1 = *x*_1_*x*_2_*x*_3_, *h*2 = *x*_1_*x*_2_*y*_3_, *h*3 = *y*_1_*x*_2_*x*_3_, *h*4 = *x*_1_*y*_2_*x*_3_, *h*5 = *y*_1_*x*_2_*y*_3_, *h*6 = *x*_1_*y*_2_*y*_3_, *h*7 = *y*_1_*y*_2_*x*_3_, *h*8 = *y*_1_*y*_2_*y*_3_. Let us designate *X* as the genotype of *h*1 and *Y* as the genotype of *h*2. Consequently, *h*2 to *h*7 represent the hybrid offspring produced by *X* and *Y* parents. If we denote *α* as the mating-bias value between *X* and *Y*, we can construct a matching compatibility table, shown in Fig 31, for these eight mating-bias genotypes based on their genetic similarity to one another.

**Fig 31.**
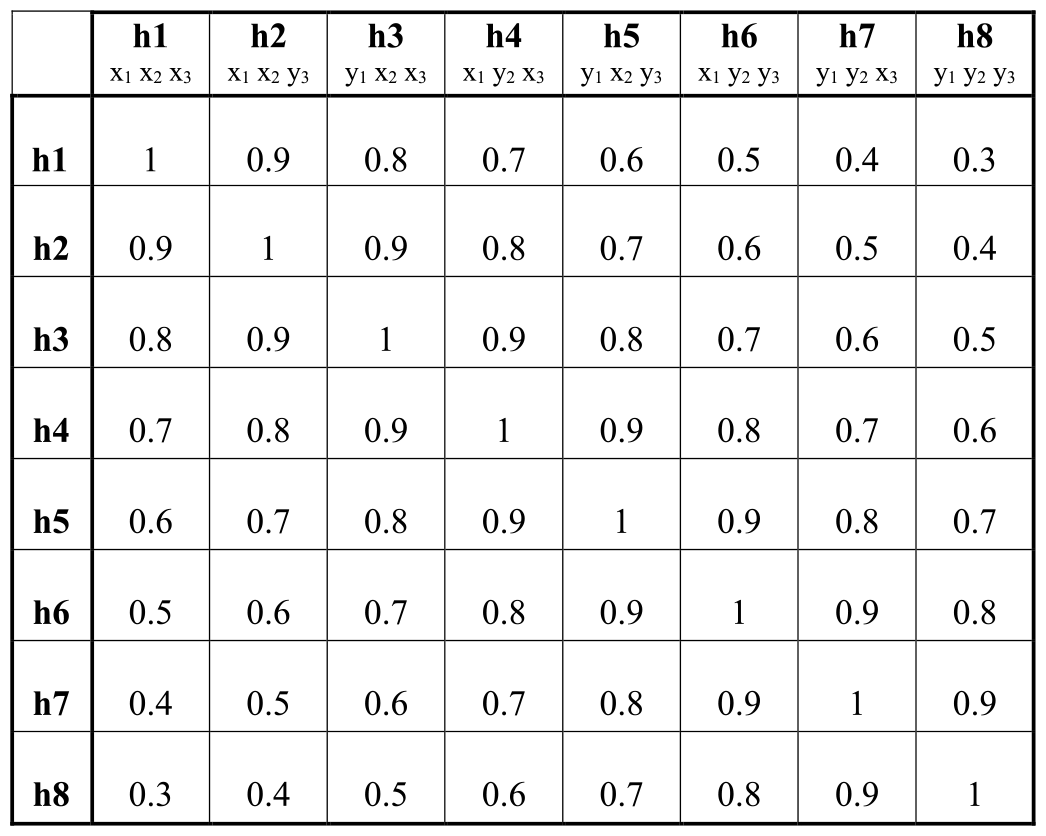
A matching compatibility table for mating-bias genotypes with three gene loci. The value in each cell specifies the probability of a successful mating match between two parental genotypes on the vertical and horizontal axes. This example uses an *α* value of 0.3. The difference in mating-bias values between adjacent genotypes is (1 − *α*)/7. Same genotypes are perfectly compatible, so their probability of a successful match is 1. The probability of a successful match between genotypes *h*1 and *h*8 is α. Subtracting the value in each cell from 1 converts the table to the format shown in Fig 3.

We developed a computer application that followed the algorithms in Fig 1 and Fig 6 to compute the phase-portrait solutions of a 2-niche sympatric system with 3-locus mating-bias genotypes (see Fig 32). For comparison, we also developed an application to solve a system with 2-locus mating-bias genotypes (not shown). The results revealed that as the number of gene loci for the mating-bias genotypes increases, the conditions required for the system to converge become more stringent. Lower values of *α* and *f*, higher values of *n*, and *NA* values closer to 0.5 are needed for the system to converge and produce fixed points. The reason for this is that, contrary to the case for ecological hybrids, there is no disruptive ecological selection against mating-bias hybrids. Consequently, all the mating-bias hybrids are ecologically viable. In our mathematical model (see Fig 5), just as the presence of viable ecological hybrids increases the effective values of *f* and diminishes ecological selection, the presence of viable mating-bias hybrids increases the effective value of *α* and diminishes sexual selection. As the gene loci for mating-bias alleles increase, more mating-bias hybrid offspring are produced, making mating-bias alleles assortment and premating RI less likely. Our results are consistent with the findings of previous researchers [32, 74].

**Fig 32.**
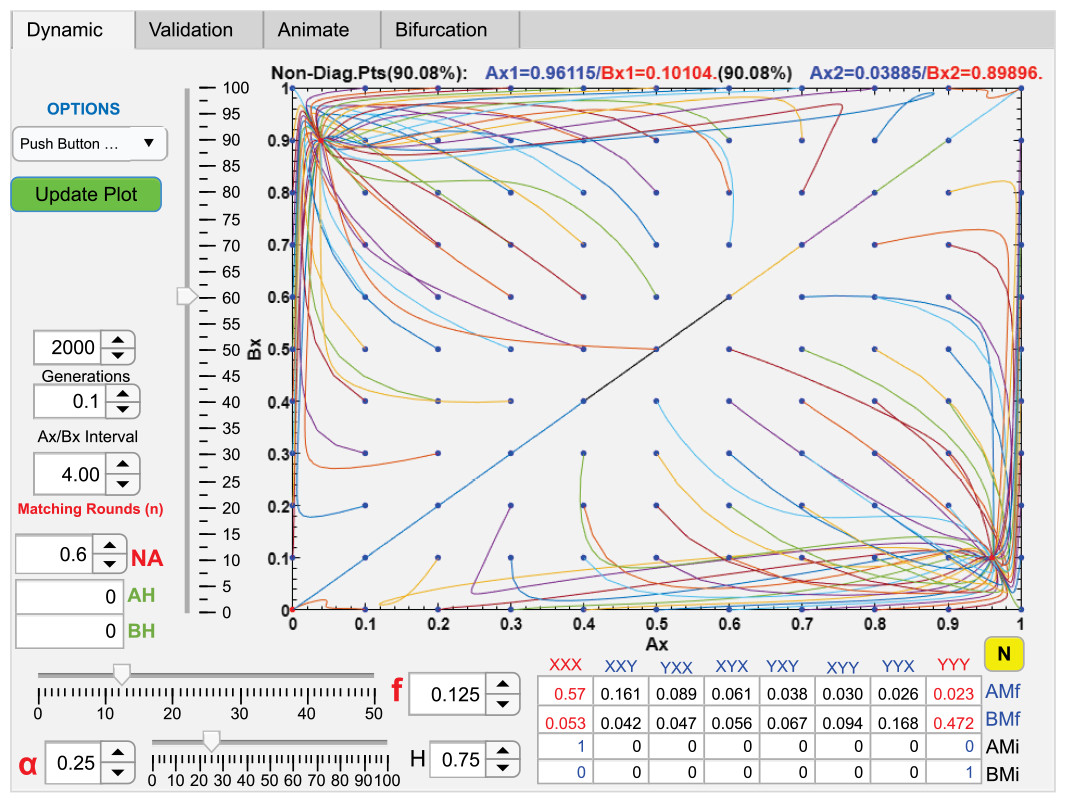
Phase portrait solution of a 2-niche sympatric system with 3-locus mating-bias genotypes. A computer screenshot shows a phase-portrait solution of a 2-niche sympatric model with 3-locus mating-bias genotypes. The model assumes no viable ecological hybrids. *AMi* and *BMi* specify the starting hybrid genotype ratios in niche *A* and niche *B. AH* and *BH* specify the total starting hybrid ratios in niche *A* and niche *B*. If the system converges, *AMf* and *BMf* show the final mating-bias genotype ratios at the fixed point in niche *A* and in niche *B*. If we let *X* represent the ratio of genotype *xxx* and *Y* represent the ratio of genotype *yyy*. Then *Ax* is the ratio of *X*/(*X* + *Y*) in niche *A*, and *Bx* is the ratio of *X*/(*X* + *Y*) in niche *A*. The rest of the parametric variables are the same as those used in Fig 9. As shown, the system converges to two fixed points at *Ax*/*Bx* = 0.96115/0.10104 and *Ax*/*Bx* = 0.03885/0.89896.

Fig 32 shows a 2-niche system with 3-locus mating-bias genotypes converging to fixed-point polymorphism given the specified set of parametric values. *AMf* and *BMf* display the ratios of mating-bias genotypes in niche *A* and *B* at the fixed point. As we create parametric conditions more favorable for generating stronger RI—by decreasing the values of *α* and *f*, increasing the value of *n*, or having *NA* closer to 0.5—we observe a greater assortment of the most incompatible mating-bias genotypes in the two different niches. In the example shown in Fig 32, improving the condition for stronger RI increases the ratio of *X* (genotype *h*1) in niche *A* and increases the ratio of *Y* (genotype *h*8) in niche *B*. The increased ratios of *X* and *Y* reduce the ratios of the intermediate hybrids (genotypes *h*2 to *h*7) and shift the frequency distribution of the hybrids toward the dominant genotype in each niche. This results in a dominant cluster of mating-bias genotypes around *X* in niche *A* and another dominant cluster around *Y* in niche *B*, producing greater premating RI between the two niches. In the limit, when *α* = 0, only the *X* genotype remains in niche *A* and only the *Y* genotype remains in niche *B*, and all the intermediate mating-bias hybrids are eliminated. Premating RI is complete and there is no gene flow between niche ecotypes. When *f* = 0, all the offspring of inter-niche matches are killed. Because of intra-niche sexual selection, only the *X* genotype or the *Y* genotype remains in each niche, and all the other mating-bias genotypes are eliminated.

When a system converges, it allows for a wide range of initial mating-bias genotype ratios in its phase portrait to converge to the same fixed point. In other words, the final, fixed-point mating-bias genotype ratios in the niches are independent of the initial mating-bias genotype ratios. This implies that the system has the ability to select the most divergent pair of mating-bias genotypes (*i. e*., the pair with the greatest mating incompatibility, such as genotypes *X* and *Y* in our example) for mating-bias assortment to produce the maximum premating RI between the two niches.

In a convergent system with multi-locus mating-bias genotypes, the origin of the phase portrait often lies within the basin of attraction of a fixed point. Consequently, a mutant allele at a locus that can create a new pair of mating-bias genotypes with greater mating incompatibility than what exists in the current system can invade and increase premating RI between the niches (see analogous example in Fig 17).

## XI. Computer Simulation of a Gonochoric Two-allele Model in a Multi-niche Ecoscape

Our individual-based computer simulation was used to study a gonochoric “two-allele model” of sympatric speciation [3, 16] in a multi-niche ecoscape, in which individuals encountered one another randomly to find mates. In the simulation, an individual in a gonochoric population has either the male sex or the female sex, and its genome is shown in Fig 33. In addition to the three gene loci for ecological alleles, the genome also contains three loci for male-trait alleles, three loci for female-preference alleles, and one locus for sex identity alleles. The sex identity locus has two alleles, *S*_*m*_ for male and *S*_*f*_ for female, which specify the sex of an individual and whether the individual’s male-trait genotype or female-preference genotype is expressed. Assume all 3-locus genotypes have alleles at each locus that are numbered from 1 to 60, so their respective genotype-to-phenotype map can be represented by a three-dimensional graph with allele axes ranging from 1 to 60. Fig 34a shows an example of such a genotype-to-phenotype map for the ecological alleles. It plots a 3-D ecoscape with 1000 randomly distributed ecological genotypes, displayed as dots, and six ecological niches, shown as shaded ellipsoid volumes.

**Fig 33.**
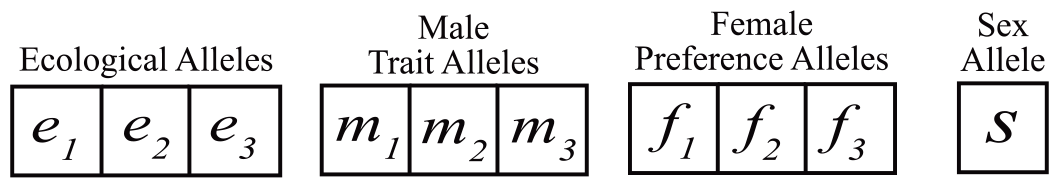
Genome of a gonochoric individual.

**Fig 34a.**
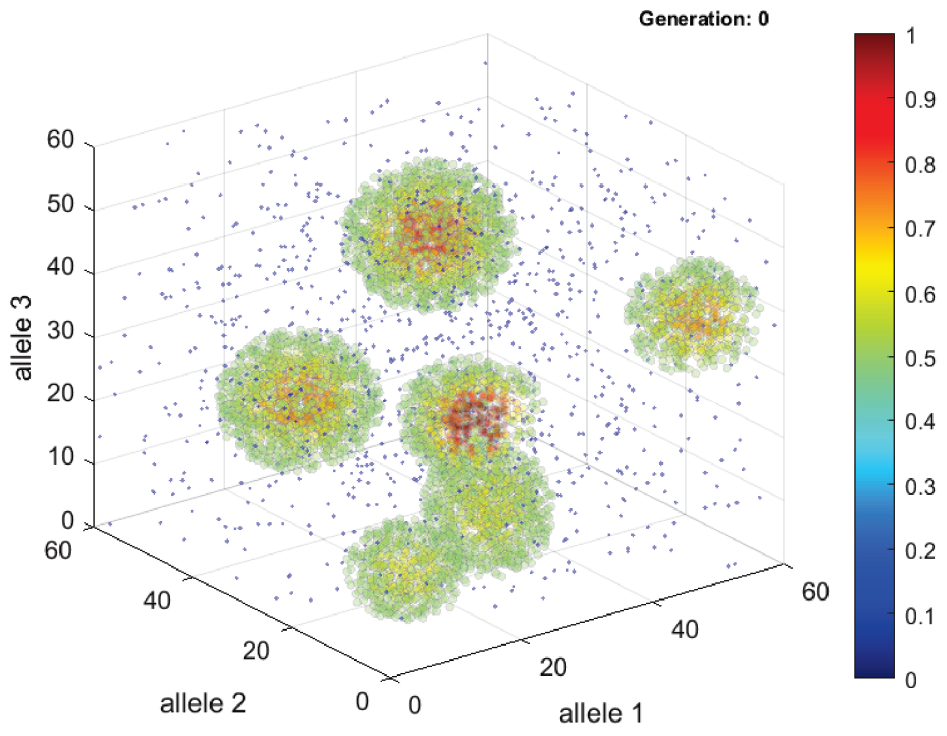
A genotype-to-phenotype fitness landscape in a multi-niche ecosystem. A computer screenshot shows the three-dimensional genotype-to-phenotype map used in the simulation of a “two-allele model” of sympatric speciation. Similar to Fig 4a, the dots represent phenotypes, the *x, y*, and *z* coordinates of which are specified by alleles in an individual’s 3-locus genotype. However, in this case, the alleles on the axes could be the *e*_1_, *e*_2_, and *e*_3_ alleles from an individual’s ecological genotype; the *m*_1_, *m*_2_, and *m*_3_ alleles from its male-trait genotype; or the *f*_1_, *f*_2_, and *f*_3_ alleles from its female-preference genotype. The graph shows the initial genotypes of 1000 individuals randomly distributed in a 3-D space. The ellipsoid probability densities of six ecological niches are also shown.

**Fig 34b.**
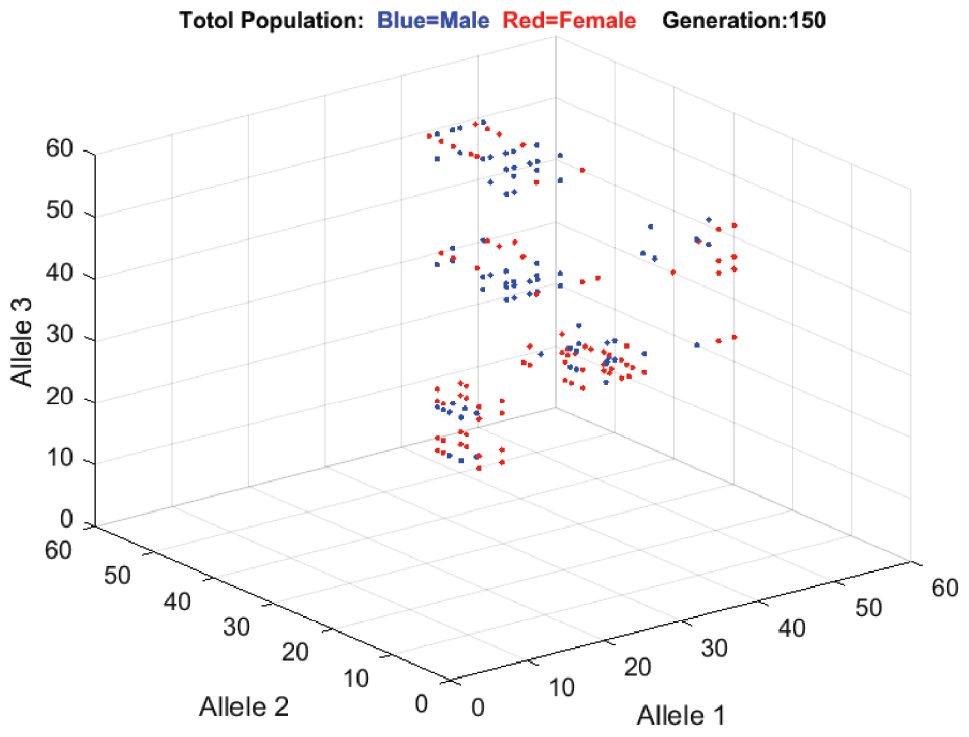
Spontaneous emergence of linkage disequilibrium among male-trait, female-preference, and ecological alleles. A computer screenshot shows the emergence of linkage disequilibrium among male-trait alleles, female-preference alleles, and ecological alleles to produce premating RI. After 150 generations, the randomly distributed male-trait and female-preference genotypes in Fig 34a spontaneously aggregated into six assorted mating groups. The genotypes of each mating group are in linkage disequilibrium with the ecological genotypes of a random niche group. This spontaneous emergence of linkage disequilibrium among random groups of male-trait alleles, female-preference alleles, and ecological alleles in separate niche populations is able to produce premating RI among all six niche groups. The blue dots represent individual male-trait genotypes, and the red dots represent individual female-preference genotypes. Because of overlaps, each dot may represent more than one individual possessing the specific genotype. The size of each mating group gradually shrank with each passing generation because of runaway selection. In this example, the female-preference variance value σ is 60, and the number of matching rounds is 10.

Similarly, alleles from the three male-trait loci can be used as axis coordinates to specify a male’s position in a 3-D genotype-to-phenotype map for male traits; and alleles from the female-preference loci, a female’s position in a 3-D genotype-to-phenotype map for female preferences The female-preference-allele coordinates specify the center of a spherical volume of probability density distribution which is constructed from three normal distribution functions along the *x, y*, and *z* axes. The means of the normal distribution functions are at the center position of the spherical volume and their variance, σ, specifies the variance of the probability density distribution in the sphere. In a matching encounter, where a male’s male-trait phenotype position lands in a female’s female-preference probability density distribution determines their chance of a successful mating match. When a male’s trait position is at the center of a female’s preference sphere, their probability of a successful match is 1. The probability of a successful match diminishes as the male’s trait position moves toward the periphery of the female’s preference sphere. After *n* matching rounds in a generation, all the unmatched individuals die without leaving offspring.

Fig 34b shows the simulation result of a gonochoric, sympatric population using the two-allele model of sympatric speciation. Initially, alleles were randomly assigned to the loci of individual genotypes without any linkage associations among them. Following the same life-cycle algorithm as in Fig 1, the male and female individuals were allowed to compete for niche resources, become mating pairs based on the compatibility of their male-trait and female-preference genotypes, and reproduce offspring by random assortment of their alleles. Given favorable parametric values, the initially uniformly distributed male-trait and female-preference genotypes in the graph were seen to congregate into six random clusters of matched male-trait/female-preference genotypes that were associated with the six niche ecotypes, and the niche groups became reproductively isolated. The genotype clusters in Fig 34b tended to become more compact with the passing of generations because of runaway selection and intraspecies sexual selection. Such ecotype, male-trait, female-preference linkage disequilibrium tended to emerge when the variance σ of the female-preference sphere was reduced below a certain value. This outcome of spontaneous linkage disequilibrium between adaptive ecological alleles and mating-bias alleles in a two-allele model of sympatric speciation is in line with the findings of previous researchers [31, 32, 45].

We can call the genotype clusters in Fig 34b the “MF clusters” (male-trait/female-preference genotype clusters that are in linkage disequilibrium with a specific ecological genotype). In our previous cases of unisex, isogamous populations, the probabilities of matching success among individuals are determined by alleles (e.g., *X, Y*, and *Z*) of various mating-bias values (e.g., *α, β*, and *γ*) at a single gene locus. Here, in the gonochoric multi-gene-locus example, the probability of a successful match between two opposite-sex individuals is determined by two variables: the distance between the male-trait position and the female-preference position (as defined by the coordinates of their 3-gene-locus genotypes) in the 3-D genotype-to-phenotype map (Fig 34a) and the variance σ of the female’s preference sphere

Computer simulations of the example in Fig 34a using just two ecological niches (see Fig 4a) showed that the system would converge with a female-preference-sphere variance value σ between 3 and 580. When the system converged, two distinct MF clusters appeared in the 3-D genotype-to-phenotype map that were as far apart from each other as possible (based on the specified value of σ) to produce the maximum RI between the two niche populations. When σ was greater than 380, the system was divergent, and only a single MF cluster associated with both ecological niches appeared. The unimodal MF cluster shrank in size with time because of runaway selection and sexual selection. When σ was less than 3, the population was prone to crashing and becoming extinct, because the small female-preference variance made it difficult for individuals to find compatible mates with a limited number of matching rounds.

Despite the higher dimensionality of its mating-bias genotypes, qualitatively, the gonochoric population exhibits the same kind of population dynamics as those of its unisex lower-dimension counterpart. Therefore, we can expect to observe the same mechanisms of invasion by higher-mating-bias mutants in the gonochoric population as those in the unisex population— such as invasion by meta-alleles that can produce higher mating biases (“one-allele model” of incremental RI) and invasion by higher-mating-bias mutants in a convergent or divergent system or in a system with a singular type of mating bias (“two-allele model” of incremental RI).

## XII. Discussion

### We can draw the following conclusions based on the results of our study

#### 1. Evolution could be viewed as swarm behavior of combinatorial genotypes and phenotypes in a multi-dimensional fitness landscape

Because of the difficulty of traveling back in evolutionary time to study the origin of species, computer simulation remains an attractive way to discover and validate the theoretical mechanisms and conditions that could have brought about a speciation event. The individual-based computer simulation presented in this study was created based on the idea that all phenotypic features of an organism are the results of interactions among its genes in gene networks. The genes that make up a network could be structural genes or genes that are responsible for interactions or communications among other genes or gene networks. The simplified genotype-to-phenotype map in Fig 4a illustrates how alleles of three different gene loci can combine to create a unique genotype. As the genotype flourishes or perishes, depending on the fitness of its associated phenotype in an environment, so do its constituent alleles increase or decrease in frequency in the gene pool. The more prevalent the constituent alleles of a genotype are in the gene pool, the more likely the genotype will reappear in the genoscape due to random mating and recombination. This solves the credit assignment problem in a merit-based system. If an allele contributes negatively to a genotype, it tends to get eliminated with the unfit genotype, and vice versa for an allele that contributes positively to a fit genotype. The alleles in Fig 4a could themselves represent unique genotypes or phenotypes that are the results of the interactions of other lower-level allele complexes, and the hierarchical process could continue all the way down to the DNA level.

This combinatorial approach to genotype and phenotype creation is attractive, because assuming each gene has several alleles, the number of possible genotypes and phenotypes that could be created by their permutational combinations is enormous [60, 75]. Adding, changing, or eliminating a constituent allele, through mutation, neutral drift, or random chance events, could create an exponential change in the number of possible combinatorial variations.

The modular and hierarchical architecture of genomes also enhances the evolvability of this combinatorial approach [76, 77]. In genomes, gene networks could be loosely grouped together based on the distinct functions that they implement and viewed as separate functional modules. Because genomes tend to be modular, functional gene networks are separated, so changes in genes within a module can create functional variants of the module (a unit of selection) for evolution to act on, but they do not change the functions of other modules. (The alternative is to have a tangled web of inseparable gene circuits, a mutation within which will make the entire genome the unit of selection in evolution.) Because genomes tend to be hierarchical, high-level genes—such as regulatory genes or genes in charge of communications and interactions among gene networks—can control the functions of large networks of genes or modules. Mutations in such high-level genes (or high-level modules) can then create novel combinations of sophisticated modular functionalities to produce new functional variants. This Lego-like shuffling and reshuffling and recombining of building-block gene networks of varying degrees of complexity and at different levels of hierarchy can potentially produce large changes in phenotypes that allow a species to jump across low-fitness valleys in a fitness landscape to reach other unexplored niches. Once a population lands and establishes a beachhead in a new niche, then microevolution can take over to find the most optimal genotype/phenotype to exploit the niche’s resources. Such a mechanism of evolution by jumping from niche to niche in the fitness landscape could help to explain the saltatory pace of speciation observed in punctuated equilibrium [34, 35]. Nonetheless, this model of evolution does not contradict examples of gradual speciation observed in the fossil record [78] as the speed of speciation often depends on the shape and slopes of a dynamic fitness landscape.

The shapes of the fitness landscape for biological organisms have been the focus of many research studies in the past three decades. The results seem to confirm that a climbable Mount Fuji type of landscape (or its equivalent high-dimensional holey landscape [79]), assumed in our study, rather than a rugged landscape resembling the Madagascar stone forest, is essential for the evolvability of species [60, 80-83]. Like sexual reproduction, the hierarchical, modular architecture of the genome and a genotype fitness landscape conducive to hill-climbing adaptation are likely evolved features in biological evolution that ensure a species’ adaptability and chance for long-term survival [83, 84].

Because phenotypes are created by interactions among nested and hierarchical networks of genes, any beneficial change in the properties of a gene in a network that increases the phenotype’s fitness will create opportunities (or positive selection pressure) for genes elsewhere in the networks to effect their own beneficial adaptive changes in response (by creating, eliminating, or substituting alternative allele variants). In our simulations, this dynamic drove a population of genotypes along a fitness gradient, pulling them from the peripheries to the centers of the ellipsoids in Fig 4a, where they reached peak fitness levels.

Evolution of a species could therefore be viewed as a cloud of swarming individual genotypes that are always sending out tentacles through mutation, drift [60], chance events, or mostly just by recombination of existing allele variants to search for niche resources in a multi-dimensional genoscape. Once it finds an available niche, it will then draw the rest of the population there, similar to what a foraging ant or bee would do [62, 63].

#### 2. Incumbent selection could allow a larger, more established population to suppress or eliminate a smaller minority population of higher fitness

The phenomenon of incumbent selection is closely tied to the concept of demographic swamping [85]. It describes how a more established, larger group of individuals could use its numerical advantage to suppress or eliminate a smaller group through their attritive interactions, even when the individuals in the larger group have lower fitness than those in the smaller group. This could happen in biological evolution. Suppose a group of birds have long beaks because it is the optimal phenotype for foraging food in an environment. Over time, the females come to prefer males with long beaks because a long beak is associated with fitness. Suppose now the environment changes, and short beaks become the better phenotype for obtaining food. Still, a mutant male with a short beak will not be able to find any female to mate with because they all prefer long beaks, even though the short-beaked male now has higher ecological fitness.

When a species becomes established in a niche, over time, its population becomes dominated by a few genotypes that are most adapted to its environment, and its gene pool consists mostly of the constituent alleles of those dominant genotypes. Nevertheless, there are ways for other rare and fringe alleles to hide and survive in the gene pool. Because of their small numbers, the rare alleles—which when combined create “mutant” gene networks and phenotypes that are unfit for the current environment—have less chance to meet and be eliminated by selection pressures. The situation changes, however, if the environment changes and makes the current niche unlivable. In this case, the dominant phenotypes and their constituent alleles are decimated, creating genomic instability, which then gives the rarer alleles opportunities to meet and combine in new gene networks to produce novel and unusual phenotypic variants that the species can use to explore other niches in the genoscape.

It has been argued that as a population becomes smaller, harmful recessive diseases are more likely to emerge [78]. Still, we must remember that fitness is relative. A genotype that is unfit (harmful) in the current environment may be fit (adaptive) when the environment is different. While it is true that most random point mutations are harmful [78], hopefully, through purifying selection, rare, lethal alleles that are incompatible with the majority of the genomic circuits are quickly eliminated, and only those rare alleles that are more benign and compatible with the rest of the genome can hide and survive in the gene pool, thus preserving genetic variability. A changing environment continuously culls the gene pool for the most adaptive alleles to construct the most optimal genotypes and phenotypes. Even though those rare alleles can create less-fit phenotypes when combined with the dominant gene circuits in the current environment, when the dominant alleles are decimated in an environmental upheaval, incumbent selection is reduced, ecological selection relaxes as the effects of chance and randomness feature more prominently in the dwindling population, and those rare alleles will have more opportunities to meet and combine to create viable and functional new gene networks that are adaptive to a new environment. It pays for a species in such a dire situation to take a risk and spread out to explore other new territories, when continuing in the same path means certain extinction.

#### 3. The interaction of ecological selection and sexual selection in sympatric speciation can be modeled by nonlinear dynamic systems

Evolution cannot happen without selection, and selection cannot happen without differential elimination of less fit individuals. The mathematical model in Fig 5 shows that ecological selection and sexual selection can act in concert to bring about assortment of two mating-bias alleles, and thus premating RI, in a sympatric 2-niche ecoscape. Their actions appear to be complementary: weak sexual selection can be compensated by strong ecological selection, and vice versa, in producing premating RI (see Fig 16). Assortative mating cost, determined by the number of matching rounds in a generation, increases the effect of sexual selection, while the presence of viable hybrids decreases the effect of ecological selection. Based on our model, stable assortment of the mating-bias alleles is unlikely unless each of the two ecotypes is able to produce enough offspring to saturate its niche’s carrying capacity. This is easily accomplished, for instance, in fish species that can spawn millions of offspring in each generation. In situations when not enough offspring can be produced to saturate a niche’s carrying capacity, stable mating-bias-allele polymorphism is still possible if adaptive individuals in the niche are able to proportionally increase their reproductive fitness by increasing their ecological fitness—because they can acquire more resources in a mostly empty niche and reproduce more offspring. Such a frequency-dependent increase in the fitness of a small group may be sufficient to counter the incumbent selection pressure from a larger group and prevent the extinction of the small group. The small group can then be kept alive at a suppressed, though stable, equilibrium state if it can achieve an average fecundity index *F* between *F*_*min*_ and *F*_*max*_. Such an exceptional scenario can be modeled by adjusting the effective carrying capacity of the smaller group’s niche to be equal to the number of individuals in the group multiplied by their increased mating frequencies. The smaller group may then be treated as one that has saturated its niche’s effective carrying capacity.

The nonlinear dynamic behaviors of our mathematical models are described by sets of difference equations. A computer application then uses numerical methods to solve the equations and displays the results graphically. Phase portrait solutions of the models show that the system either converges to fixed points, which act like attractors in a vector field, becomes partially convergent, or is nonconvergent. The locations of the fixed points are determined by parameters such as niche carrying capacities (*NAmax*,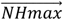, *NBmax*), strength of ecological selection (*f*), mating bias (*α*), and assortative mating cost (*n*), but are independent of the starting mating-bias-allele ratios (*Ax*,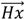, *Bx*) in the niches. In the phase-portrait vector field (Fig 9, 11, and 12), both ecological selection (specified by *f*) and sexual selection (specified by *n* and *α*) interact to determine the nonlinear dynamic behavior of the non-diagonal populations. However, only sexual selection determines the dynamic behavior of the populations on the diagonal line.

Based on our results, premating RI tends to be reversible and less permanent than other forms of RI. Changes in environmental parameters can cause two specialist ecotypes that have not achieved complete premating RI between them to merge into a generalist group with a singular type of mating-bias allele. For instance, when the mating-bias alleles of the two niche groups are at a fixed-point polymorphism in a convergent system that still allows introgression, fusion into a generalist group may occur when environmental change supports the emergence of viable hybrids, which reduces ecological selection and can cause the system to become divergent.

Because it is difficult to have complete sexual and ecological selection (*f* = 0 or *α* = 0) at the outset, most fixed-point solutions tend to create mating-bias-allele polymorphism in the development of premating RI, and a certain degree of introgression is expected between incipient species [65, 86]. Nonetheless, the reduced gene flow between niche groups can decrease the homogenizing effect of recombination and encourage the emergence of other isolating mechanisms, such as drift or differential adaptation, to complete the speciation process [87].

The emergence of premating RI is most favored when the assortative mating cost is low, ecological selection is strong, mating bias is high, and the populations of the two niches are approximately equal, such as when parameter values are close to *NA* = 0.5, *f* = 0, and *α* = 0 in Fig 16. As parameter values move away from this ideal condition, toward a diagonal line connecting the coordinates (*f* = 0.5, *α* = 0) and (*f* = 0, *α* = 1) in Fig 16, the likelihood of fixed-point convergence and premating RI rapidly disappears. In general, convergence to a fixed-point polymorphism is more likely with low values of *f* (high ecological selection), low values of *α* (high mating bias), high values of *n* (low assortative mating cost), *NA* = 0.5 (roughly equal niche sizes), and high enough values of *off* (number of offspring) to saturate all niche carrying capacities (or a fecundity index *F* > *F*_*min*_ for unsaturated niches).

In our dynamic animations, ecological selection usually occurs faster than sexual selection in systems that converge. The total strength of sexual selection in our model appears to be jointly determined by the values of *n* (the number of matching rounds) and *α* (the degree of mating bias). Very strong sexual selection caused by *n* close to 1 (but not the strong sexual selection caused by low values of *α*) or very weak sexual selection caused by *α* close to 1 (but not high values of *n*) may not be optimal for the development of fixed points and RI.

#### 4. Invasion by high-mating-bias mutant alleles can create or increase reproductive isolation

Our computer animation has shown that the origin (at *Ax* = 0 and *Bx* = 0) of a vector-field phase portrait can act like an unstable saddle point when fixed points exist in the system and the origin falls within their basins of attraction (Fig 9 and 12). This implies that when only a single type of mating-bias allele exists in two niche populations that are undergoing disruptive ecological selection, a mutant allele with a suitable high mating-bias value *α* will be under positive selection pressure to invade and establish fixed-point mating-bias-allele polymorphism and thus premating RI between the two populations (Fig 17). Fig 18 shows a representative plot of the generation time needed for a small population of such a mutant mating-bias allele to be swept into a stable fixed-point equilibrium. Invasion by such a small population of high-mating-bias alleles becomes more difficult as search cost (*θ*) increases (Fig 19).

The caveat to be kept in mind is that computer calculations based on infinite population size and idealized mathematical models do not account for the risks of random events that can occur in real life when a starting population is extremely small. In our idealized model of monogamous individuals in Fig 18, there is almost no assortative mating cost associated with very-high-mating-bias mutant alleles for *n* > 1. In real life, however, an invading mutant allele with high mating bias will have difficulty finding mates initially and risks being eliminated by random chance before it can attain a threshold population size, even though it is under greater positive selective pressure to multiply. In the limiting case, a single high-mating-bias mutant with *α* = 0 will not be able to find a matching mate. This may be one of the reasons why incipient sister species tend to have weak RI (caused by low-mating-bias alleles) between them [87]. Because ecological selection and sexual selection tend to be complementary in producing fixed-point convergence (i.e., a weaker ecological selection demands a stronger sexual selection for convergence and vice versa), a system with more viable hybrids will require a mutant with a higher mating bias to invade, which will be more prone to elimination at its inception. How these two opposing forces, random chance and selection pressure, interact to determine fixation time warrants further study.

Our computer simulation has shown that a mutant meta-allele that produces a higher mating bias (a lower value of *α*) can invade and replace an existing meta-allele that produces a lower mating bias (a higher value of *α*) in a convergent 2-mating-bias-allele, 2-niche sympatric system (“one-allele model” of incremental RI). Similarly, computer simulation and numerical solution of a 3-mating-bias-allele, 2-niche sympatric system have also demonstrated that a third mutant allele with a low intra-niche mating bias (high *β*) and a higher inter-niche mating bias (*γ* < *α*) can invade a convergent 2-mating-bias-allele, 2-niche system to increase RI between the two niche groups (“two-allele model” of incremental RI). Analogously, a divergent system can also be invaded by such high-mating-bias meta-alleles and third mating-bias alleles to become a convergent system. However, to accomplish this, the invading mutants usually need to overcome much higher invasion thresholds by having higher mating biases and larger initial population ratios than those required to invade a comparable convergent system—something that is more easily accomplished, perhaps, via gene flow in parapatric speciation, or through introgression or hybridization in secondary contact following allopatric or peripatric divergence.

#### 5. Source of high-mating-bias alleles in sympatric speciation

A major criticism of sympatric speciation models is the lack of high-mating-bias alleles in natural populations that are available to produce premating RI between ecotypes under disruptive ecological selection [3, 43, 88-90]. In sexual selection, a more populous type of mating-bias allele can use its numerical advantage to eliminate a smaller, incompatible type of mating-bias allele, especially when the incompatibility (as measured by 1 − *α*) between them is large. A more sexually discriminative individual will have more difficulty finding mates when the search cost is high and is more likely to end up without offspring. These theoretical considerations argue against the existence of alternative, highly incompatible mating-bias alleles in natural populations. Yet, in nature, genetic and phenotypic polymorphisms appear to be the rule rather than the exception [3, 91], especially when many genes are involved in producing quantitative traits. Many mechanisms—through pleiotropy, spatial and temporal environmental heterogeneity [92-94], heterosis [95], transgressive segregation [96], the transporter process [97], adaptive plasticity [98], hybridization or rearrangement of ancestral genes [75, 99] or standing genetic variations [100, 101], etc.—have been proposed to explain these observed polymorphisms. In an environmental upheaval, disruptive ecological selection could decimate the dominant allele types and allow rare alleles to meet and combine to produce novel phenotypes [75]. Analogously, uncommon lower-level constituent alleles could also meet and combine to generate novel variants of mating-bias alleles. Obviously, such high-mating-bias alleles could always be donated by a parapatric species, through secondary contact, or by other related sympatric species through introgression [99, 102]; but absent such outside help, how could sympatric speciation happen?

Our model suggests several other possible ways to overcome these theoretical objections to sympatric speciation. First, the phase-portrait solutions of our model’s nonlinear difference equations (see Fig 9) demonstrate that a small population of high-mating-bias alleles can coexist with a dominant population of mating-bias alleles in a polymorphic fixed-point equilibrium, as long as there is a small ecological niche available in the ecoscape to support the existence of the high-mating-bias alleles. (For instance, for *α* = 0.2, *NA* = 0.1, *f* = 0.3, *n* = 4, a fixed point exists at *Ax* = 0.364 and *Bx* = 0.0168 in the model without viable hybrids.) The results shown in Fig 18 also suggest that it is easier for a high-mating-bias allele to invade a small niche than a large niche. Later, when ecological disruptive selection opens up a bigger niche, the existing high-mating-bias alleles could then be co-opted and used to achieve premating RI for the bigger niche.

Second, as previously argued (see Fig 17 and 18), when environmental conditions are favorable, a high-mating-bias mutant allele (or a mutant gene network from a combination of rare alleles) will be under positive selection pressure to invade a homogeneous population with a single type of mating-bias allele and be swept into fixed-point polymorphism. The examples in Fig 25 and Fig 26 also demonstrate that a meta-allele or a third allele with high inter-niche mating bias can invade a convergent system of low-mating-bias alleles. A divergent system caused by low-mating-bias alleles could also be invaded by such high-mating-bias mutant alleles, even though the invasion threshold is typically higher.

Third, because it is usually easier for a high-mating-bias allele to invade a convergent system than a divergent system, it might be easier to let a medium-strength mating-bias allele establish fixed-point equilibrium first in two niche ecotypes that are under disruptive ecological selection. The partially reduced gene flow between the ecotype groups could then lower the threshold for other high-mating-bias mutant alleles to invade, which further reduces gene flow between the two niche groups. The process could repeat itself until a polymorphism of high-mating-bias alleles is achieved—e.g., through the one-allele and two-allele models of incremental RI.

Fourth, because in our mathematical model, the two arms of selection (ecological selection and sexual selection) in Fig 5 are qualitatively indistinguishable and complementary, any mechanism that can increase the gain or loss in those two arms will produce the same dynamic behavior in the system. For instance, extrinsic and intrinsic postzygotic barriers such as maladaptive hybrid oviposition or clutch size [103], asynchronous hatchling emergence and transgressive segregation [104], sex ratio distortion, gametic incompatibility, zygotic mortality, hybrid inviability, hybrid sterility, and hybrid breakdown [73, 105] can all decrease the value of *f* in ecological selection. Similarly, parental imprinting [106], habitat preferences, chromosomal inversion and rearrangement [107], and other mechanisms of temporal, behavioral, and mechanical mating isolation [108] can decrease the value of *α* in sexual selection. Different types of premating barriers can also act cumulatively to increase RI [109]. Because premating RI through assortative mating can evolve much faster than intrinsic post-zygotic barriers, premating RI is usually the dominant driver of RI in the initial stage of sympatric speciation. In later stages, ongoing hybrid loss in disruptive ecological selection can fuel the emergence of other RI mechanisms to minimize the loss. In sympatric speciation, these myriad mechanisms are recruited by positive selection pressures in a process called adaptive coupling [66, 67, 110] to produce ever higher reproductive isolation between the niche populations and complete the speciation process [87]. Empirical studies have shown that multiple imperfect isolating barriers usually accumulate and interact late in the speciation process to consolidate and complete RI [87, 105, 111]. The validity of the coupling phenomenon is also supported by a recent study showing that both premating RI and post-zygotic RI accumulate faster in sympatric speciation than in allopatric speciation [112].

Two mechanisms are thought to facilitate coupling [67, 109, 113]. Coupling leads to linkage disequilibrium among the genes responsible for different reproductive barriers. Individuals possessing such coupled genomes can benefit from stronger overall RI when interacting with individuals from a different niche. As a result, the coupled genomes have a greater selective advantage than uncoupled genomes when it comes to invading and multiplying. This is because they are better at extracting the new niche resources created by disruptive ecological selection and/or unfilled niche capacities. In the Fig 4a illustration, if we let each axis of the 3-D plot represent alleles of a gene locus responsible for a RI mechanism and position the newly-created niche resource at the point where the alleles of the three gene loci produce maximum RI, then we should observe the same population dynamics driving individual genotypes to cluster at the location of maximum RI and establish linkage disequilibrium (i.e., coupling) among the different reproductive barrier genes. Alternatively, we can view the *X* and *Y* alleles in our 2-mating-bias-allele system as super-alleles resulting from the interactions of different barrier alleles at a lower level. Selection of the *X* and *Y* alleles in a fitness landscape should then lead to similar selection and linkage disequilibrium of those barrier alleles at the lower level.

Increased selective advantage of coupled loci caused by selective sweep or genetic hitchhiking is another mechanism that can promote coupling [67, 109, 113]. The coincidence of different barrier effects results in a stronger overall barrier than would be caused by selection on any single barrier alone. Because of such linkage disequilibrium, genes of a weak reproductive barrier can experience an increased selective advantage to multiply as a result of their association with the other barrier genes through coupling. As more barrier effects are recruited through coupling (see “cline attraction” in [67]), linkage disequilibrium of barrier loci becomes widespread in the genomes, and the overall RI is increased, making the reversal of RI difficult.

Fifth, because different types of reproductive barriers tend to interact synergistically, multiplicatively, and additively to produce the final RI [23, 109], many weak barriers, when combined, could interact to produce strong overall RI. As long as there are still unused niche resources left in the system— whether from hybrid loss, unfilled ecological niches, or a fluctuating environment that undermines the stability of current barriers—these resources can generate positive selection pressures for additional types of barriers to emerge and increase RI. Consequently, adding a different barrier mechanism may produce an outsized effect in increasing RI, and a low-mating-bias allele in a different barrier mechanism can produce the effect of a high-mating-bias allele in the overall RI.

Lastly, because the strength of mating bias *α* that is needed to produce fixed-point polymorphism is determined by the values of other variables such as *NA* (niche population ratio), *f* (strength of ecological selection), and *n* (the number of matching rounds), changing those other parametric values may relax the requirement for high-mating-bias alleles to achieve RI.

For instance, the prism-shaped fixed-point solutions in Fig 16 show that the actions of *f* and *α* are complementary in bringing about RI; therefore, an environmental change might create such a low value of *f* that the threshold required for *α* to achieve convergence is lowered and thus allow existing low-mating-bias alleles in the population to produce fixed-point polymorphism. As another example, in certain cases during disruptive ecological selection, when a larger niche group is capable of eliminating a smaller niche group through incumbent selection, the smaller group could nonetheless use an increased fecundity ratio *F* to achieve a higher niche population, and thus a higher effective niche carrying capacity, which might then make convergence to a fixed point possible. The ensuing reproductive isolation might be sufficient to free the smaller niche group from the suppressive effect of incumbent selection to reach its actual niche carrying capacity.

The flip side is that a convergent system with fixed-point polymorphism can be changed into a divergent system when parametric variables change. For instance, the convergent system shown in Fig 26 will become divergent if we increase the ratio of viable hybrids *H* (i.e., decrease hybrid loss or increase the effective value of *f*), and the RI between the niche populations will disappear. This reveals the transitory and reversible nature of premating RI by mating-bias-allele polymorphism in sympatry. Hopefully, after an initial reproductive barrier is established and gene flow is reduced, other coupling mechanisms, such as intrinsic postzygotic barriers, could be recruited to make the RI permanent.

#### 6. The effect of having multi-locus mating bias alleles

Our basic 2-niche mathematical models have assumed distinct mating-bias alleles (e.g., *X, Y*, and *Z*) on a single gene locus. Even though the mating biases among the alleles are incomplete (*α, β*, and *γ* values <1), no mating-bias hybrids are produced in the models because the parents transmit the alleles as discrete units to their offspring. This avoids the problems of having mating-bias hybrid offspring in the sexual selection arm of the model in Fig 5 and makes system convergence to fixed-point polymorphism most likely. This is because the presence of viable mating-bias hybrids, just like the presence of viable ecological hybrids, can inhibit system convergence and prevent, or reverse, the emergence of premating RI. In ecological selection, the unfit intermediate hybrid ecotypes are eliminated by disruptive ecological selection. Therefore, to have the most favorable condition for system convergence, it is desirable to have many gene loci for the ecological genotypes as this reduces the effective value of *f*. On the other hand, because there is no disruptive ecological selection on the mating-bias genotypes, all the mating-bias hybrids are ecologically viable. Consequently, it is desirable to have as few gene loci as possible for the mating-bias genotypes as this prevents increasing the effective value of *α*. These requirements support the conclusion of previous researchers that an increase in the number of loci affecting ecological adaptations facilitates speciation, while an increase in the number of loci affecting mating bias has the opposite effect [32, 74].

A system with single-locus mating-bias alleles can be realized by having a few alleles with large, dominant effects regulate mating biases. Furthermore, these alleles, or their resulting mating traits, should be inherited as discrete units from parents to offspring, thereby preventing the formation of intermediate hybrids. In our models, when the one-locus system converges, there is a differential assortment of the two mating-bias alleles, *X* and *Y*, in the two niches. The *Ax* and *Bx* values at the fixed point determine the proportions of the mating-bias alleles that are associated with the different niche ecotypes. For instance, assume that in a fish species, the *X* allele causes red coloration, and the *Y* allele causes blue coloration, traits that are used in mate discrimination. In the early stage of sympatric speciation, if premating RI is incomplete, we should observe the majority of fish in niche *A* to be red while a small minority of them are blue. Conversely, we should see exactly the opposite color proportions for fish in niche *B*. There seems to be some empirical evidence for the transmission of such discrete mating-bias traits that are inherited without giving rise to intermediate hybrids. For instance, the Midas cichlid fishes from the crater lakes of Nicaragua seem to exhibit gold/dark color dimorphism based on ecological niche specialization. This is possibly because their color pigmentation is a Mendelian trait determined by a two-allele locus with gold dominant over dark and almost complete penetrance [25, 26].

We do not need to have two discrete alleles on a single gene locus to generate distinct mating trait inheritance without producing hybrid offspring. Suppose there are two independent gene clusters, *C* and *S*, in the genome of a fish species. Let gene cluster *C* control the coloration of the fish based on the mating-bias mechanisms described in our model, and let cluster *S* contain gene circuits that could create vertical body stripe patterns. Initially, let us assume that, because of intrasexual selection, all fish have a red color and can interbreed freely. Now, if under disruptive ecological selection, a mutation in cluster *S* creates a mutant stripe pattern on the red fish, with a favorable mating-bias value *α* between the striped and the unstriped red fishes, then the striped pattern can invade and establish fixed-point polymorphism with the unstriped pattern through the invasion mechanism described in Fig 17. In this case, we can view the striped red fish as possessing mating-bias trait *X* and the unstriped red fish as possessing mating-bias trait *Y*, and the *X* and *Y* traits interact according to the matching compatibility table in Fig 3. Because the color and stripe traits are generated by separate gene circuits, they are inherited as distinct traits, and no intermediate mating-trait hybrid offspring are reproduced.

Our study results have shown that in a 2-niche system with single-locus mating-bias alleles, at most two types of mating-bias alleles can coexist because of competitive niche exclusion. In our model with three mating-bias alleles and two niches, a third mating-bias allele was always eliminated, which caused the system to revert to a model with just two mating-bias alleles (Fig 24).

Nonetheless, the seemingly discrete mating-bias alleles could themselves be “super-alleles” arising from interactions among lower-level constituent alleles [31, 32, 79]. Consequently, by mapping the fitness landscape of the mating-bias alleles, we can establish a corresponding fitness landscape for their constituent alleles. This combinatorial mechanism can be a source of mating-bias-allele innovation and polymorphism. However, even though more variants of mating-bias alleles may be created by the constituent alleles in this way, multi-locus mating-bias alleles can generate intermediate mating-bias hybrids that hinder system convergence and speciation. As long as premating RI between the niches is not complete, the reproduction of hybrid offspring by multi-locus mating-bias genotypes is inevitable.

Phase-portrait solutions of 2-niche models with 2-locus and 3-locus mating-bias genotypes showed that as we improve the conditions for the emergence of stronger premating RI—by decreasing *α* and *f*, increasing *n*, or making *NA* closer to 0.5— the system can converge and create increasing assortment of the most incompatible pair of mating-bias genotypes between the two niches. Meanwhile, the intermediate mating-bias hybrid genotypes are suppressed, and their ratio distributions are shifted toward the dominant mating-bias genotypes in each niche (see example in Fig 32). In the limit, when *α* = 0, only the most incompatible pair of mating-bias genotypes remain in the two niches; premating RI is complete, no gene flow exists between the niches, and all the intermediate mating-bias hybrids are eliminated.

Empirically, these results predict that we should see two dominant mating-bias morphs (e.g., red and blue fishes) distinguishing the two ecotypes in the early stage of sympatric speciation. Moreover, we should also expect to observe small populations of intermediate hybrid morphs that resemble the dominant mating-bias morphs in each niche (e.g., varying shades of reddish-purple fish in niche A and bluish-purple fish in niche B). Because more mating-bias hybrids are eliminated as RI becomes stronger, the prevalence and distribution of the mating-bias hybrids in different niche populations can be a measure of the degree of RI in incipient sympatric speciation.

Because the presence of viable mating-bias hybrids reduces the strength of sexual selection and makes the conditions for system convergence more stringent, any mechanism of disruptive sexual selection that can decrease the fitness of mating-bias hybrids will relax the requirements for fixed-point convergence and facilitate sympatric speciation. This could happen because the traits conferring ecological adaptation also serve as mating-bias traits, i.e., they are “magic traits” [50, 51]. Alternatively, the mating-bias traits may serve as honest signals of local adaptation, e.g., red coloration is correlated with local adaption to niche *A*, and blue coloration is correlated with local adaptation to niche *B*. In other words, mating-bias traits are indicators of ecological fitness [114-116]. In both cases, because of such correlations, the intermediate mating-bias hybrid genotypes are associated with the less-fit ecological hybrid genotypes, and they are also subject to elimination by disruptive ecological selection.

Other mechanisms of disruptive sexual selection include reducing the matching success of mating-bias hybrid morphs and reducing hybrid survival. For instance, changes in the turbidity of lake water may make it more difficult for a fish species to recognize the less conspicuous intermediate-colored hybrids as mates [117]. Females may perceive hybrid males with intermediate sexual traits as less attractive [116, 118, 119]. Male-male competition and aggression against intermediate mating-bias phenotypes may reduce hybrid fitness [120-122]. Our computer analysis has confirmed that incorporating these mechanisms in our models relaxes the parametric requirements for developing premating RI and sympatric speciation.

#### 7. Extension of the unisex, 2-mating-bias-allele, 2-niche model

So far, our computer simulation and mathematical models used haploid individuals that are seemingly hermaphrodites. It can be proved, however, that the same results apply if gonochoric individuals (which have either male or female sex) are used. This is because nature has a way to ensure an equal sex ratio through various negative feedback mechanisms. Parents tend to produce equal male and female offspring populations with the same niche allele compositions. Since eligible mating individuals of the same sex would essentially appear invisible to each other, the calculations ultimately yield the same results. To apply the same methodology to diploid, gonochoric individuals would require the construction of a new genotype-to-phenotype map that takes into account dominant and recessive effects among alleles.

A migration variable *P* can be incorporated in our sympatric models to create contact barriers between ecological niches and simulate conditions of reduced gene flow in parapatric, peripatric, and allopatric populations. Preliminary results from a 2-mating-bias-allele, 2-niche system without viable hybrids show that the presence of contact barriers can facilitate and increase premating RI by mating-bias-allele fixed-point polymorphism. However, as the contact barriers increase, the effect of sexual selection begins to dominate in the isolated niches. In the system’s phase portrait, the convergence percentage shrinks as sexual selection begins to drive the most numerous types of mating-bias alleles in populations near opposite ends of the diagonal line toward fixation (Fig 29). This also makes invasion by high-mating-bias mutant alleles difficult. When the contact isolation is complete (*m* = 0), there is no gene flow between the two niches, and sexual selection will drive the most numerous types of mating-bias alleles in the now allopatric niches toward fixation (Fig 30).

In Fig. 34b, simulation results of a “two-allele model” of sympatric speciation [3, 16] using gonochoric individuals demonstrate that under favorable conditions, random genotypes of male traits and female preferences will pair up in linkage disequilibrium and congregate in different ecotypes under disruptive selection to produce RI among niche groups [31, 32]. The crucial variable here appears to be σ (the variance of the female-preference sphere), which is equivalent to the variable *α* for discrete mating-bias alleles. The value of σ controls the degree and range that a female-preference genotype finds different male traits desirable. To have maximum mating bias, the value of σ should be such that the overlaps among female-preference ranges in different niche groups are minimized. However, a very small value of σ also exposes a picky female to elimination by sexual selection, especially when the mate searching cost is high. Despite the model’s increased complexity and dimensionality, we observed analogous population dynamics and invasion mechanisms as those in its simpler, unisex, 2-mating-bias-allele, 2-niche counterparts. This parallelism suggests the existence of fundamental, discoverable general operating principles underlying all forms of sympatric speciation. It remains to be seen whether the alleles of a metagene that can control the value of σ, when thrown into the mix of random matching and allele assortment, may also assort and align themselves in linkage disequilibrium to select the most appropriate value of σ for each niche group to achieve RI.

Multiple cues (color, hue, size, shape, olfactory, acoustic, etc.) can be used in mating discrimination. Theories and some empirical evidence suggest that having a few loci with large effects in mating trait differentiation is most effective in facilitating sympatric speciation [31, 32, 123, 124]. However, a recent study has argued that divergent selection acting on a large number of loci may be more effective in creating and maintaining genomic differentiation, e.g., by having one or several traits with polygenic basis or by having a combination of multiple traits each with a simpler genetic basis [11, 110]. Perhaps the number of traits necessary for mate discrimination also depends on context. The presence of many similar confounding traits in a species’ environment can make accurate mate discrimination difficult if too few criteria are used. On the other hand, parsimony would demand that the least number of traits necessary are used for mate discrimination, and using too many discriminating features can quickly tax and overwhelm an individual’s cognitive processing ability.

#### 8. Mechanisms and stages of sympatric speciation

Observations from our study prompt us to propose the following general statements regarding the role of sexual selection in sympatric speciation. Disruptive ecological selection creates new niches for suitable mating-bias alleles to assort and establish linkage disequilibrium among themselves and with different niche ecotypes to produce premating RI. The drivers behind these newly created niche resources come from minimizing hybrid loss in disruptive ecological selection and from acquiring unused resources in unfilled ecological niches under incumbent selection. Reproductive isolation in sympatric speciation, therefore, is an active process driven by positive selection pressures in newly created niches that are the result of disruptive selection and incumbent selection, and it can be achieved in hundreds to thousands of generations or less [15, 19, 125]. This is in contrast to RI in allopatric speciation, which occurs as a passive chance byproduct of genetic drift and geographically-isolated adaptations to different environments, and it can take an estimated 1000 to 100,000 generations to complete [14, 15].

If the above conjecture is true, that premating RI in sympatry is just the result of mating-bias alleles and their linkages finding and adapting to niche resources in a multi-dimensional genotype-phenotype space, then we should see adaptation phenomena such as niche competitive exclusion, invasion of a niche by a fitter variant, etc., in its dynamics. The simulation results from the 2-mating-bias-allele and 3-mating-bias-allele models seem to support this. Sexual selection, therefore, can help the genes of a species to more effectively extract and utilize available niche resources in its ecoscape, by minimizing wasteful hybrid loss and acquiring unused niche capacity.

When two ecological niches are so different that specialized phenotypes are necessary for their efficient utilization, spontaneous assortment of mating-bias alleles can produce reproductive isolation that splits a generalist species into two specialist species for the benefit (from a gene’s eye view) of all in the species [33, 126]. Yet, the existence of invasion thresholds helps avoid the danger of excessive speciation [45], in which small specialized populations lose the genetic diversity available to them and are prone to extinction in narrow, precarious niches [127]. There are advantages and disadvantages of being generalists, and there are advantages and disadvantages of being specialists. The number of habitats available in an ecosystem is not limited to the number of discrete niches. A generalist can subsist on a combination of discrete niches to hedge its chance of survival. Finding the best mixture of generalists and specialists to most efficiently extract resources from a vast combinatorial number of niches in an environment quickly becomes a complex optimization problem for evolution to solve so that the phenomenon of life can persist. When species divergence is advantageous in a sympatric environment, it makes sense that evolution would find a more expedient mechanism than allopatric speciation to split an ineffective generalist species that constantly suffers from inferior hybrid loss into more efficient and viable specialist species, instead of waiting patiently for the slow emergence of geographic barriers, and sexual selection might just be the vehicle that helps it accomplish this goal.

Because premating RI by mating-bias-allele fixed-point polymorphism in sympatry tends to be reversible (e.g., decreasing the hybrid loss between two sympatric specialist species can breakdown the premating RI and coalesce them into a single generalist species), other types of reproductive barriers are probably needed to make sympatric speciation permanent and irreversible. For instance, intrinsic postzygotic barriers can replace extrinsic environmental selection to ensure hybrid loss [73]. This can lessen the effect of environmental contingencies on the existing premating RI and make it less reversible.

It has been argued that in sympatric speciation, the process that causes the evolution of the first reproductive barriers is different from the processes that bring about the subsequent transition to strong RI and completion of the speciation process [87, 105, 127]. The rationale is that once the barrier to gene flow is sufficiently strong, it can facilitate a wide range of new processes—such as reinforcement, mutation-order divergence, or the Bateson–Dobzhansky–Muller (BDM) model of incompatibility—which were previously suppressed by gene flow, but now they can be recruited to further enhance and consolidate the reproductive barrier [66]. This distinction between the evolution of first barriers and the subsequent development of strong RI seems to be supported by our finding that it is much easier to incrementally increase RI in a convergent system than to convert a divergent system into a convergent one [128].

Based on our findings, we propose the following five stages of sympatric speciation:

In Stage 1, a sympatric population of a single species subsists on a single ecological niche. There is no ecological selection, and sexual selection favors the predominant type of mating-bias allele in the population (i.e., the low-mating-bias alleles) and eliminates the less common, less compatible (high-mating-bias) alleles. Consequently, the population harbors mostly low-mating-bias alleles.

In Stage 2, environmental change creates new niches in the habitat of the sympatric species. The ensuing disruptive ecological selection favors specialized phenotypes that are adaptive to the new niches and exerts negative selection pressure against the inferior intermediate hybrid phenotypes.

In Stage 3, when conditions are favorable, as determined by niche carrying capacities, assortative mating cost, clutch size, hybrid loss, invasion threshold, etc., selection against high-mating-bias alleles is reversed. High-mating-bias alleles are now under positive selection pressure to invade and establish fixed-point polymorphism (premating RI) between niche populations to stem the ongoing hybrid loss. The high-mating-bias alleles could come from de novo mutation, recombination of more elementary constituent alleles, transfer by introgression from other parapatric species or sympatric species or from secondary contact, hybridization or reuse of ancestral genes or standing genetic variations [75, 99-101], or co-optation of existing intermediate-strength mating-bias alleles in the population when hybrid loss is sufficiently severe. This process establishes the first reproductive barriers in sympatric speciation.

In Stage 4, after a certain degree of RI between niche populations has evolved (i.e., when the system has become convergent), the reduced gene flow could then recruit other mechanisms—such as the one-allele and two-allele models of incremental RI, and other prezygotic and postzygotic barriers through adaptive coupling—to further enhance and consolidate the reproductive barriers and complete the speciation process [87]. The splitting of the initial species into sister sympatric species with ever diminishing degrees of introgression between them becomes more and more difficult to reverse. As hybrid loss is reduced, so is the selection pressure to recruit additional RI, and eventually the BDM model of incompatibility through drift and differential adaptations takes over. The BDM mechanism of divergence is accelerated because the reproductively separated sympatric ecotypes are adapting to very different niche environments.

In Stage 5, sympatric speciation is complete. Fixed-point polymorphism of high-mating-bias alleles (premating isolation) and other prezygotic and postzygotic reproductive barriers recruited through adaptive coupling consolidate interspecies reproductive isolation. Very little gene flow occurs between niche populations, which minimizes the effects of hybrid loss and ecological selection. Within the now separate and reproductively isolated sympatric species, the effect of sexual selection again dominates to eliminate other less common, incompatible intraspecies high-mating-bias alleles. The speciation process returns to Stage 1, when it is again uncommon to find high-mating-bias alleles within each sympatric species.

Notice that in our model, reproductive isolation can evolve rapidly once certain threshold requirements are met, as demonstrated by the sharp transitions in Fig 15, 17, 25, 26, and 28. Linkage disequilibrium and the effects of a positive feedback loop appear to be the key components driving the dynamics of such rapid transitions [129-131]. In Fig 17, for instance, a sigmoid-shaped transition in mutant population growth is present because the fitness of the invading mutant is both frequency-dependent and resource-dependent. Initially, as the invading mutants increase in number, their fitness increases because it becomes easier for a mutant to encounter and match with other compatible mutants in the population. A positive feedback loop propels the mutant population into the accelerated growth phase of the sigmoid curve. Later, as available niche resources are exhausted, mutant population growth levels off, and it becomes more difficult to eliminate the remaining less-fit native populations (similar to the difficulty of eliminating harmful recessive alleles in a population when they become rare).

Because adaptive coupling of reproductive barriers also involves linkage disequilibrium as well as frequency- and resource-dependent fitness, we can expect it to exhibit similar types of sigmoid-shaped transitions. Sudden evolutionary transitions can occur when small changes become coupled to each other in a positive feedback loop [130]. This selection-driven rapid pace of sympatric speciation appears to be compatible with the phenomena of “tipping point,” “snowball effect,” and “threshold effect” described by other researchers [36, 73, 87, 129, 132-134].

The short transition phase in sympatric speciation means that the presence of high-mating-bias alleles in a sympatric population during the speciation process is ephemeral. Once speciation is completed, only low-mating-bias alleles can be detected in sympatric species because of intraspecies sexual selection.

#### 9. Is sympatric speciation the ugly duckling that has grown up to prove Darwin wrong?

This study has presented a model of sympatric speciation that is driven primarily by selection (sympatric RI). This contrasts with the Neo-Darwinian model of allopatric speciation, which is based primarily on the BDM model of genetic incompatibilities that arise as chance byproducts of neutral drift and differential local adaptations of geographically isolated populations (geographic RI). Our hope is to find a mechanism of sympatric speciation that is theoretically and biologically realistic enough to rival the prevailing model of allopatric speciation. The crux of the question appears to be how likely speciation is in the absence of geographic barriers. We can attempt to find the answer to this question by examining how sympatric speciation and allopatric speciation can occur in the following two scenarios.

First, let us examine how neutral genetic drift can cause speciation. Consider a sympatric population that has adapted to a single uniform ecological niche. According to our model, because there is no disruptive ecological selection, sympatric speciation is impossible, and sexual selection favors the predominant mating-bias alleles (low-mating-bias alleles) and acts to eliminate the less numerous, incompatible mating-bias alleles (high-mating-bias alleles).

Suppose a major environmental upheaval now divides the habitat into two halves and establishes complete geographic RI between the two divided populations. Theoretically, the two allopatric subpopulations can now diverge through the BDM model of genetic drift, but this is likely to take a very long time. Because the divided ecological niches are the same, the divided allopatric populations are not under any selection pressure to diverge, and the same stabilizing selection in the two niches will tend to maintain the same ecological genotypes and mating-bias genotypes in the two isolated populations. Such stabilizing selections are probably responsible for the stable phenotypes of many living fossil species, such as horseshoe crabs, African coelacanth fish, Komodo dragons, and pig-nosed turtles. The conserved morphological appearances of these species have shown little change in the fossil record for hundreds of millions of years, and by inference, perhaps their underlying ecological genotypes and mating-bias genotypes as well. The effects of stabilizing selection can be eroded by fluctuating population sizes, however. This is because the impact of randomness in small populations can create founder and bottleneck effects that exaggerate genetic differences and accelerate the pace of allopatric speciation by neutral drift (as in peripatric speciation). Likewise, in the special case when both allopatric populations need to adapt to new, but otherwise identical habitats, mutation-order divergence [135] is facilitated because there is already geographic RI between the populations.

Second, let us consider the scenario in which the population of a species has to adapt to two different ecological niches. This is the primary model of allopatric speciation proposed by Charles Darwin after he visited the Galapagos islands in 1835. Darwin realized that the many species of finches he found on the Galapagos islands actually originated from common ancestors on the mainland, and those finches evolved into different species because of the geographic isolation that developed after their migration from the mainland and because of their differential adaptation to local island environments. The Neo-Darwinists later buttressed the theory with the BDM model of genetic incompatibility.

Darwin’s proposal of allopatric speciation via geographic isolation and gradual adaptive change through natural selection in different environments is powerful and intuitive and has received wide empirical support. We do not doubt the validity of Darwin’s mechanism, but we do ask (1) whether the existence of geographic barriers is necessary for a population to speciate when two different ecological niches arise in its habitat that cause disruptive ecological selection of its phenotypes and (2) whether allopatric speciation, rather than sympatric speciation, is the predominant mode of speciation in nature.

The models in our study have shown that, under the right conditions, sympatric speciation can occur in a population undergoing disruptive ecological selection without the need for geographical barriers. Intuitively, it also makes no sense for a sympatric species to continue to suffer ongoing hybrid loss— and thus reduced species fitness because it has become less effective at extracting resources from its environment—without evolving some ways to rectify the situation.

For speciation to happen in a population undergoing disruptive ecological selection, initially some sort of reproductive barrier needs to be erected between the different niche populations to jumpstart the process. Premating barriers through mating-bias-allele polymorphism in sympatric RI is fast but easy to reverse, while the BDM model of post-zygotic barriers through geographic RI tends to be slow but more difficult to reverse. Yet, both sympatric RI and geographic RI produce the same isolating effects. In the later stages of speciation, they both reduce gene flow and allow the BDM mechanism of incompatibility to kick in and complete the speciation process. Perhaps the only difference is that in the beginning stage of speciation, geographic RI can produce reproductive barriers right away without any preconditions, while initial sympatric RI has more stringent conditions and invasion thresholds to overcome.

In allopatric speciation by differential adaptation, the more different the ecological niches are, the faster reproductive incompatibilities may emerge by chance between niche populations. Because sympatric speciation is mainly driven by selection, it can recruit other selection-driven processes such as adaptive coupling and incremental RI to consolidate the initial premating barriers. The BDM mechanism of divergence by differential adaptation is facilitated because the reproductively isolated sympatric ecotypes can adapt to their very different niche environments without the hindrance of recombination effects. Initial sympatric RI tends to be leaky and allows for a large degree of introgression between incipient species. Geographic RI can also be leaky, because divided populations can migrate both ways across geographical barriers. If migration is predominantly one way, then the founder effect and adaptive radiation commonly seen in island biogeography are likely to dominate the speciation process.

Because of the glacial pace and low likelihood of developing geographical barriers by chance amidst a sympatric population, the most likely scenario for allopatric speciation to occur is for a founder population to migrate across an existing geographic barrier and become adapted to different niches, as exemplified by the speciation of Darwin’s finches. Still, once a founder population reaches a new, unfamiliar territory, it is likely to branch off into many different subspecies to occupy all the available niches by adaptive radiation. Adaptive radiation is common in island biogeography and is mostly sympatric in nature [134, 136]. Besides cichlids [5-11], other examples of island adaptive radiation and sympatric speciation include the anole lizards of the Caribbean islands [137, 138] and Darwin’s finches of the Galapagos islands [139, 140], which have been observed to speciate in as fast as a few generations.

Allopatric speciation is hard-pressed to account for the immense number and diversity of species in tropical rainforests, where no obvious geographical barriers exist. In a resource-rich and heterogenous environment such as a rainforest, many niches are always opening and closing in a species’ habitat, as all the species compete in an evolutionary arms race [141, 142], and new niches are created when a species inadvertently acquires new capabilities, either through genetic drift, preadaptation, or co-optation [60]. Bats, for instance, underwent explosive diversification when they took to the sky and found a new world of unexplored niches. Today, there are more than 1,300 species of bats in the world, making them the second most common group of mammals after rodents. Allopatric speciation no doubt happened, but given the more abundant opportunities for sympatric speciation, maybe it is time for us to reevaluate our null hypothesis to examine whether sympatric speciation, not allopatric speciation, is the predominant mode of speciation in nature. [4, 69].

Nonetheless, firm empirical support for the primacy of sympatric speciation is still lacking [2, 18, 43, 143], mostly because of the rarity of observable speciation events in human time. The estimated time to speciation among plants and animals based on models of allopatric speciation is around two million years in one study [144]. This may be the reason why most scientifically observable speciation events in nature are sympatric. Allopatric speciation and sympatric speciation both produce the same speciation end results, making it impossible to know for sure which one was the responsible mechanism because we were not there to witness it. Compounding the difficulty, our model has shown that the existence of high-mating-bias alleles in sympatric speciation is transient, and once speciation is completed, intraspecies sexual selection would erase any evidence that sympatric speciation ever happened.

Still, some observable predictions can be made based on the different mechanisms of speciation. Because sympatric RI is actively driven by selection, not by chance RI as in allopatric speciation, we can expect speciation events to happen rapidly in sympatric speciation [129, 130, 134]. In the fossil record, we should see rapid splitting of new species from their common ancestors, even in the absence of major geological events, followed by long periods of relative stasis, as stabilizing ecological and sexual selections confine the species in their respective niches. This saltatory mode of evolution is most consistent with the phenomenon of punctuated equilibrium [34, 35], in contrast to the gradualism advocated by Darwin, and could explain the rarity of intermediate forms in the fossil record.

#### 10. Limitations of the study

The British statistician George Box famously wrote, “All models are wrong, but some are useful.” All models are simplified approximations, and none can perfectly represent real-world conditions. This is especially true in biology, where myriad entities and variables often interact in highly stochastic and nonlinear fashions. In our study, we have deliberately limited the parameter space to make the analyses of our models tractable, at the risk of omitting many important variables— such as different mating systems, operational sex ratios, genomic structures, epistasis and pleiotropy effects, genetic drift, mutation rates, dominant and recessive traits, environmental heterogeneity, etc.—factors that may be important for specific biological populations being studied. Nonetheless, our skeletal models could serve as a foundational framework that can be expanded to tackle more complicated systems.

Our mathematical models assume an infinitely large population size. Consequently, their predictions may not hold true when taking into account the effects of random events in finite and small populations. For instance, in solving for the invasion dynamics of our models, the population ratio of a mutant could be lowered to 1 × 10^−15^ without any fear of it being eliminated by random chance, an assumption that is unrealistic in natural systems. In the real world, stochasticity and small population size can have a major impact in determining the plausibility, speed, and waiting time of a successful mutant invasion. At best, our models’ phase-portrait solutions only provide a rough guide of the underlying expected vector-field forces driving the dynamic evolution of natural populations.

#### 11. Suggestions for future research

Our study has shown that modeling the development of premating RI in sympatric speciation can be a messy, complicated process, involving complex nonlinear dynamics and difference equations that are intractable to analytical solution. Real-life genotype-to-phenotype maps and fitness landscapes are much more dynamic and complex than those in our simplified toy models. This complexity likely reflects the reality in nature, that sympatric speciation is also a complicated process involving many nonlinear processes, variables, and contingencies. In this paper, we have presented a simple model of a multiple-mating-bias-allele, 2-niche sympatric ecosystem with few input parameters, which, hopefully, still accurately captures and clarifies the fundamental operating principles in sympatric speciation. Our goal has been to find the foundational laws of sympatric speciation that are applicable even in increasingly complex scenarios. Next, it will be interesting to investigate how different matching and mating arrangements [51] and different forms of assortative mating cost can affect the behaviors of our models. We may also examine how multiple mating-bias alleles with different strengths, and meta-alleles that can modulate their strengths, could evolve and interact to bring about, or prevent, sympatric speciation in hermaphroditic and gonochoric populations. Models of sympatric speciation based on resource competition also warrant further study [31]. It is possible that we could model such systems, which usually consist of a single niche with a unimodal resource distribution and have individuals with varying degrees of ecotype specialization, by creating discrete contiguous niches of graded carrying capacities in our gonochoric, multi-niche system.

We hope that the findings of this study will open the door to yet another promising approach to tackle and solve the mysteries of sympatric speciation.

## APPENDIX

This appendix details the derivation of mathematical equations used in the paper.

Assume in a sympatric species, there are two groups of individuals with distinct ecotypes that are adaptive to two niches, *A* and *B*, in a 2-niche sympatric ecoscape. In a mating generation, all the eligible unmated individuals in the sympatric ecoscape encounter one another randomly in matching rounds to find compatible mates. After a matching round, all the matched individuals pair off to produce offspring for the next generation and are taken out of the mating pool. After a maximum allowable number of *n* matching rounds in a generation, all the unmatched individuals die without offspring.

If *PA* is the number of individuals in niche *A* and *PB* is the number of individuals in niche *B, P*_*p*_ = *PA*+ *PB* is the total population in the ecoscape. Let *NA* be the ratio of population in niche *A*, and *NB* be the ratio in niche *B*, i.e., *NA* = *PA*/(*PA*+*PB*), *NB* = *PB*/(*PA*+ *PB*). Assume also that individuals in the population have a multi-locus ecological genotype that defines their ecological fitness in different niche environments, as well as a single gene locus for two mating-bias alleles *X* and *Y* that determine their mating compatibility. Fig 2 shows an example of a genotype that has three ecological-allele loci and one mating-bias-allele locus that was used in our computer simulation. Let *Axr* be the ratio of individuals in niche *A* that have the *X* mating-bias allele, and *Ayr* = 1 − *Axr*, the ratio of individuals in niche *A* that have the *Y* mating-bias allele. Similarly, let *Bxr* and *Byr* represent the ratios of niche-*B* individuals that possess the *X* and *Y* mating-bias alleles.

The population *P* in the 2-niche ecoscape can be divided into four normalized groups according to the schematic diagram shown in Fig 5. *NAx* = *NA*× *Axr* is the ratio of niche *A* individuals that have the *X* mating-bias allele. Similarly, *NAy* = *NA*× *Ayr, NBx* = *NB* × *Bxr, NBy* = *NB* × *Byr*, such that *NA* = *NAx* + *NAy, NB* = *NBx* + *NBy*, and *NAx* + *NAy* + *NBx* + *NBy* = 1.

In Fig 5, the arrowed lines represent the probabilities of encounters among the different groups. For instance, the arrowed line connecting *NAx* and *NBx* denotes that their probability of encounter is 2 × *NAx* × *NBx*. The symbol *f* at the head of the arrows specifies the ratio of *NAx* (or *NBx*) offspring that the *NAx* (or *NBx*) group can expect to recover from such an encounter, assuming that a matched individual from each group only produces one offspring (i.e., a unit offspring). For our 3-ecological-locus individuals in the computer simulation, assume there are no viable hybrids (i.e., all the hybrid offspring are eliminated because there is no niche for their ecological genotypes), then *f* = 1/8, and the unit offspring ratio that the encounter between *NAx* and *NBx* will produce is ⅛ × 2 × *NAx* × *NBx*.

Similarly, the symbol *α* denotes the ratio of matched individuals from an encounter, based on a prespecified matching compatibility table (Fig 3). For instance, the unit offspring ratio that group *NAx* and group *NAy* can expect to produce from their encounter is *α* × 2 × *NAx* × *NAy*. For groups that belong to different ecological niches and possess different mating-bias alleles (e.g., between groups *NAx* and *NBy*), the expected ratio of unit offspring that will be produced from their encounters is a multiple of *α* × *f* (e.g., *α* × *f* × *NAx* × *NBy*).

Notice that all the probabilities of encounters add up to 1: *NAx*^2^ + *NAy*^2^ + *NBx*^2^ + *NBy*^2^ + 2 *NAx NBx* + 2 *NAx NAy* + 2 *NBx NBy* + 2 *NAy NBy* + 2 *NAx NBy* + 2 *NBx NAy* = 1. In addition, without the influence of the multipliers, *α* and *f*, (i.e., when *α* = 1, and *f* = 0.5) the sum of unit offspring ratios (that is, assuming that each matched individual only reproduces one offspring) produced by all the encounters is 1.

### 2-Mating-Bias-Allele Model with No Viable Hybrids

Let us first consider the case when there are no viable hybrids.

In each matching round, the probabilities of encounters of the different groups (*NAx, NAy, NBx, NBy*) can be represented by the elements of a matching probability matrix, *M*, in Fig 7, which are just the product of a vector [*NAx NBx NAy NBy*]^T^ and its transpose. At the beginning of each generation, let us specify the normalized population ratios before the first matching round as: *Ax*_*o*_ = *NAx. Ay*_*o*_ = *NAy*. *Bx*_*o*_ = *NBx. By*_*o*_ = *NBy*.

Notice that in matrix *MM*, we always have normalized population groups, so that the ratios *Ax* + *Bx* + *Ay* + *By* = 1, and the sum of all the elements (i.e., the probabilities of encounters of the normalized groups *Ax, Bx, Ay*, and *By*) in *M* is 1. The matrix *MM* can be viewed as being composed of the following four submatrices:

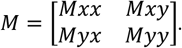

In the first matching round, the probabilities of encounters between the four different groups can be specified as follows: *Mxx*_*o*_= [*Ax*_*o*_ *Bx*_*o*_]^T^ × [*Ax*_*o*_ *Bx*_*o*_], *Myy*_*o*_ = [*Ay*_*o*_*By*_*o*_]^T^ × [*Ay*_*o*_*By*_*o*_], *Mxy*_*o*_= [*Ax*_*o*_ *Bx*_*o*_]^T^ × [*Ay*_*o*_ *By*_*o*_], *Myx*_*o*_ = [*Ay*_*o*_ *By*_*o*_]^T^ × [*Ax*_*o*_ *Bx*_*o*_].

After the first matching round, there is a 100% match for the probability encounters in *Mxx* and *Myy* because they have the same mating-bias alleles. However, the match rate in *Mxy* and *Myx* is only *α*, because they have different mating-bias alleles. We can then specify the accumulated matched pairs after the first matching round in terms of the probabilities of encounters in the submatrices: *ΣMxx*_(1)_ = *Mxx*_*o*_, *ΣMyy*_(1)_ = *Myy*_*o*_, *ΣMxy*_(1)_ = *α* × *Mxy*_*o*_, and *ΣMyx*_(1)_ = *α* × *Myx*_*o*_.

Let us consider the unmatched individuals after the first matching round. The proportions of unmatched individuals are (1 – *α*) × *Mxy* and (1 – *α*) × *Myx*. Because the ratio of individuals in *NAx* that participate in each probability encounter that involves *NAx* is exactly 1/2 of the encounter’s probability ratio, the ratio of unmatched *NAx* individuals is:

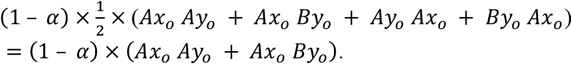

Next, we can calculate the total ratio of the unmatched population after the first matching round, *P*_1_, as

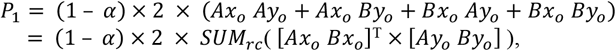

where the function *SUM*_*rc*_(*A*) sums all the elements of matrix *A*. Alternatively, if we let *X*_*o*_ = *Ax*_*o*_ + *Bx*_*o*_ and *Y*_*o*_= *Ay*_*o*_+ *By*_*o*_, then

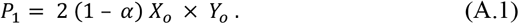

The new normalized ratio of unmatched *NAx Ax*_(1)_ in the unmatched population *P*_1_ is

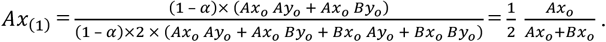

Similarly, the normalized ratios of other unmatched groups in the unmatched population are

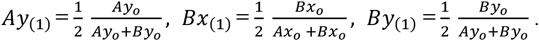

Note that the normalized ratio of unmatched individuals with the *X* alleles is *X*_(1)_ = *Ax*_(1)_ + *Bx*_(1)_ = 1/2, and the normalized ratio of unmatched individuals with the *Y* alleles is *Y*_(1)_ = *Ay*_(1)_ + *By*_(1)_ = 1/2.

Using the same calculations, it can be shown that after matching round 2,

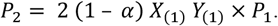

But *X*_(1)_ = *Ax*_(1)_ + *Bx*_(1)_ = 1/2, *Y*_(1)_ = *Ay*_(1)_ + *By*_(1)_ = 1/2.

Therefore,

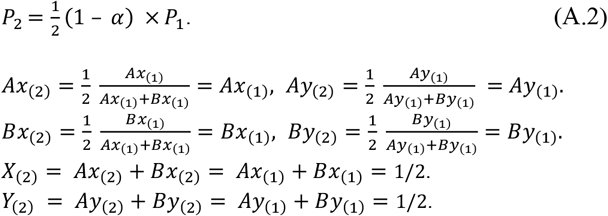

The additional matched individuals produced by matching round 2 are then

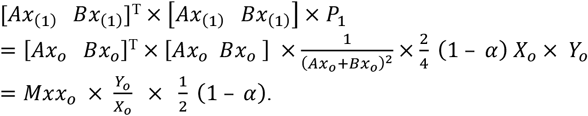

Similarly, using the same derivation, it can be shown that

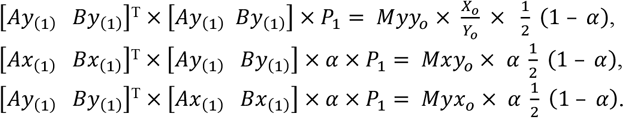

The accumulated matched individuals after matching round 2 are then

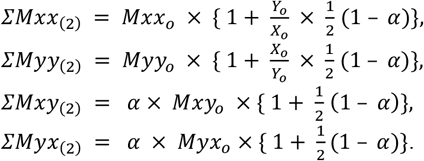

After matching round 3:

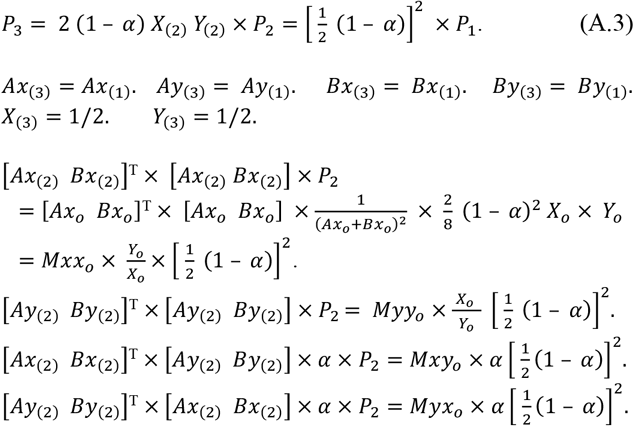

Therefore,

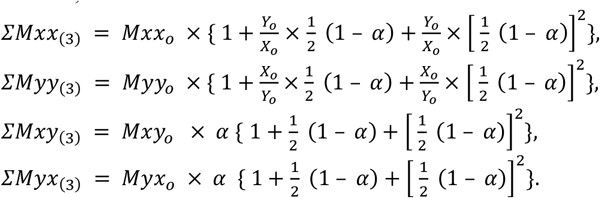

Utilizing the property of the geometric series,

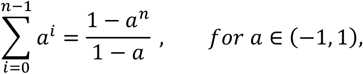

we can show that after the *n*^*th*^ matching round, the accumulated matched individuals are

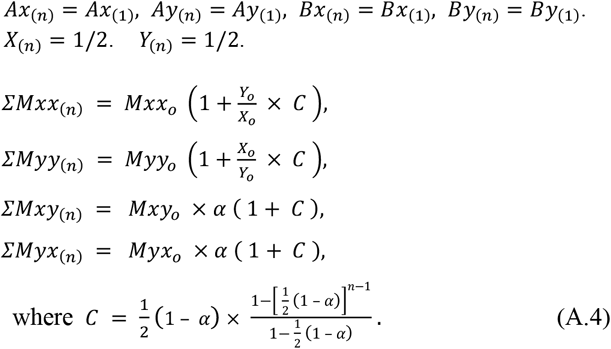

The unmatched individual after the *n*^*th*^ matching round is

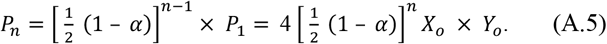

Now if we reconstitute Matrix *ΣM*_(*n*)_

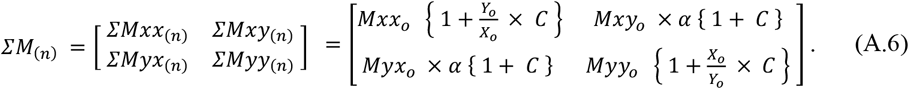

The elements in matrix *ΣM*_(*n*)_ specify the probability ratios and compositions of all the matched individuals in the ecoscape after the *n*^*th*^ matching round in a generation. Notice that summing all the elements of *ΣM*_(*n*)_ and adding the result to the ratio of unmatched individuals return a probability of one: *SUM*_*rc*_ (*ΣM*_(*n*)_)+ *P*_*n*_ = 1.

Next, we can determine the composition of unit offspring produced by each matched pair of individuals in *ΣM*_(*n)*,_ according to independent assortment at each locus of the parents’ ecological alleles and mating-bias alleles, using a unit offspring matrix, *U*, (see Fig 8). Unit offspring means that each matched individual is only allowed to reproduce one offspring to replace itself. Each cell of the unit offspring matrix *U* specifies the ratios of the different types of offspring (*Ax, Bx, Ax, Bx, Hx, Hy*) that each combination of matched parents will produce, based on independent assortment of alleles and the ecological selection variable *f*. Notice that all the ratios in each cell of *U* add up to 1. Therefore, if *α* = 1 or *n* = ∞, i.e., everyone in the population gets to mate, then a normalized mating population of 1 will produce a normalized total unit-offspring population of 1. The unit offspring produced may be expressed mathematically as

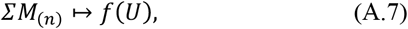

where the symbol ↦ *f*(*U*) denotes element-wise multiplication of the elements of *ΣM*_(*n*)_ (which specify the probability ratios of different parental matches) by the offspring proportions in each cell of *U* to produce different types of offspring. The different types of offspring are then added up to produce an output vector 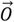 that shows the total unit offspring ratios (not normalized) produced from the generation:

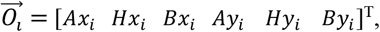

where *i* is the generation number.

Alternatively, we can decompose *UU* into submatrices that only contain ratios specific to each individual offspring type, i.e., 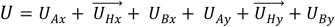. Then,

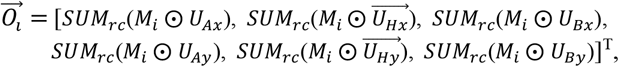

where ⨀ is the Hadamard product that performs element-wise multiplication of two matrices of the same size. *M*_*i*_ = *ΣM*_(*n*)_ in generation *i*.

Or it can be calculated in another way to yield the same result:

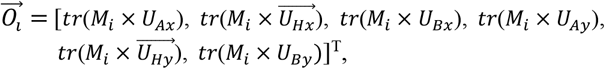

where *tr* (*A*) is the trace of a square matrix *A*.

Notice again that because each matched parent only produces one offspring to replace itself,

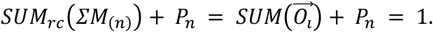

Now we can summarize the equations as follows:

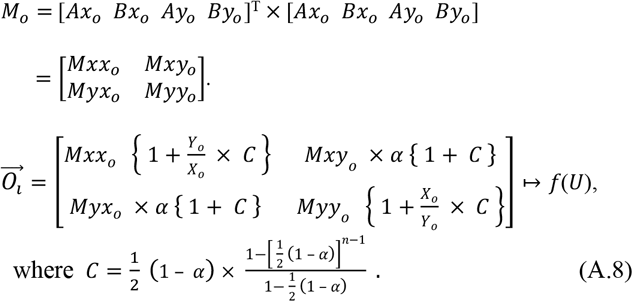

If we assume there are no viable hybrids, i.e., *Hx*_*i*_ and *Hy*_*i*_ are not in our consideration (*Hx*_*i*_ = *Hy*_*i*_ = 0), then the equations can be expanded and written in the following form:

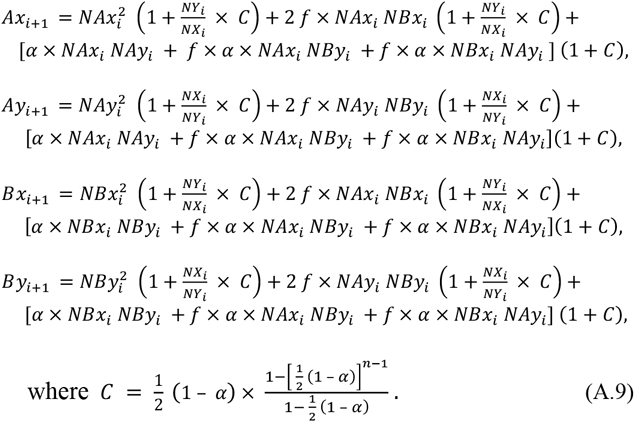

*NAx*_*i*_, *NAy*_*i*_, *NBx*_*i*_, and *NBy*_*i*_ are the normalized population ratios at the start of mating generation *i*, and they represent offspring from the previous (*i* − 1) generation that have survived ecological selection to become eligible mating individuals for generation *i. NX*_*i*_ = *NAx*_*i*_ + *NBx*_*i*_ and *NY*_*i*_ = *NAy*_*i*_ + *NBy*_*i*_. *Ax*_*i*+1_, *Ay*_*i*+1_, *Bx*_*i*+1_, and *By*_*i*+1_ are the ratios of unit offspring (not normalized) that were produced from the matching rounds in generation *ii*. Notice that the unmatched population from generation *i* is

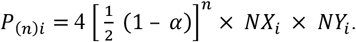

For the special case when *NAx*_*i*_ + *NBx*_*i*_ = *NX*_*i*_ = 0 or *NAy*_*i*_ + *NBy*_*i*_ = *NY*_*i*_ = 0, i.e., when the *X* or *Y* allele is absent,

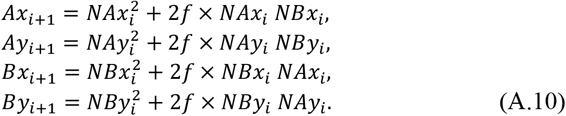

Next, we can multiply the unit offspring ratios (*Ax*_*i*+1_, *Ay*_*i*+1_, *Bx*_*i*+1_, and *By*_*i*+1_) by the number of offspring (*off*) that each matched mating individual is supposed to reproduce, and then multiply the result by the generation’s starting population 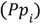 to obtain the offspring population for the next generation. From these results, we can calculate the starting population 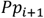 and the normalized ratios *NAx*_*i*+1_, *NAy*_*i*+1_, *NBx*_*i*+1_, and *NBy*_*i*+1_ for the next generation.

In the case that *off* is sufficiently high so that the number of offspring produced in each generation can completely saturate the carrying capacities of the two niches, *NAmax* and *NBmax*, then the population ratios of niche *A* and niche *B* going into the next generation are fixed. The normalized *NAx, NAy, NBx*, and *NBy* ratios going into the next generation can then be calculated as follows:

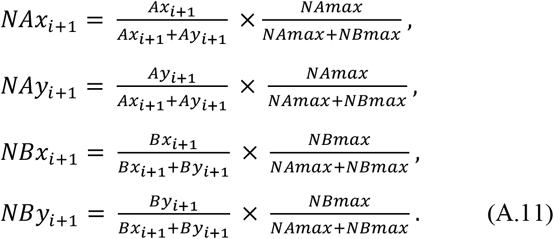

These difference equations (A.9–A.11) describe the dynamic behavior of a 2-niche, 2-mating-bias-allele sympatric system with no viable hybrids in our mathematical model.

### 2-Mating-Bias-Allele Model with Viable Hybrids

We can use the same type of mathematical derivation to obtain the difference equations for ecosystems in which hybrids are viable. We will use the derivation of our 3-locus-ecotype, 2-mating-bias-allele, 2-niche sympatric system with viable hybrids as an illustrative example (Fig 2, 3, and 4).

When hybrids are viable, instead of just having 2 ecological niches, *NA* and *NB*, as shown in Fig 5, we now have six additional ecological niches for the hybrids, *NH*_2_, *NH*_3_, *NH*_4_, *NH*_5_, *NH*_6_, and *NH*_7_, with maximum carrying capacities of *NHmax*_2_, *NHmax*_3_, *NHmax*_4_, *NHmax*_5_, *NHmax*_6_, and *NHmax*_7_. Assume that the alleles in each ecological allele locus are either *a* or *b*. Let *aaa* represent the genotype of individuals in niche *NA*, let *bbb* represent the genotype of individuals in niche *NB*, and arrange the genotypes of individuals in the hybrid niches according to their proximity to the *NA* and *NB* genotypes; then an individual can only have one of the following permutations of alleles in their 3-locus ecological genotype: *NA* = *aaa. NH*_2_ = *aab. NH*_3_ = *baa. HN*_4_ = *aba. NH*_5_ = *bab. NH*_6_ = *abb. NH*_7_ = *bba. NB* = *bbb*. The resulting eight ecotypes can be mapped by the corners of an imaginary cube in Fig 4a.

This will then allow us to construct a unit offspring matrix *U* (analogous to the one in Fig 8) that specifies the types and proportions of offspring that a matched pair of parents will produce, according to random assortment of the parental alleles. For instance, a pair of matched individuals with genotypes *aaa*(*NA*) and *aba*(*NH*_4_) will produce unit offspring ratios that are 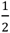 *aaa*(*NA*) and 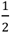 *aba*(*NA*_4_).

Similarly, instead of having just four interacting groups of populations, *NAx, NAy, NBx*, and *NBy*, we now have 12 additional interacting hybrid populations that can be represented by the elements of two vectors:

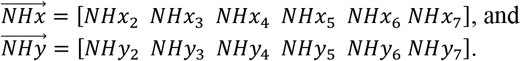

We can normalize all 16 interacting groups so that

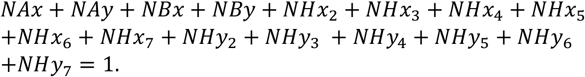

At the beginning of each mating generation, let us specify the normalized population ratios before the first matching round as 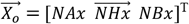 and 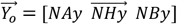. Then the vector 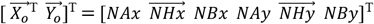 multiplied by its transpose will produce the matching probability matrix *M* in Fig 20. Let

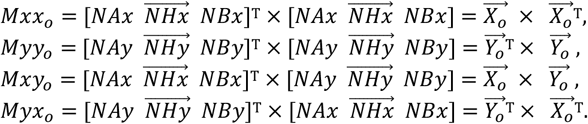

We can then specify the accumulated matched pairs after the first matching round as the probabilities of encounters in the submatrices:

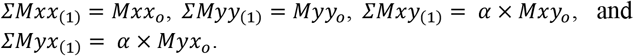

Using the same derivation as that in the system with viable hybrids (A.1), after the first matching round, the total proportion of unmatched individuals is:

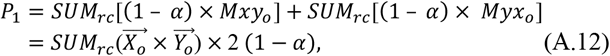

where the operation *SUM*_*rc*_ (*A*) sums all the elements of matrix *A*.

Because the ratio of individuals in each group of 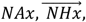, and *NBx* that participate in a probability encounter that involves the group is exactly 1/2 of the encounter’s probability ratio, the ratios of unmatched individuals in groups 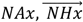, and *NBx* can be represented by the vector

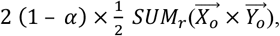

where the operation *SUM*_*r*_(*A*) sums the elements across each row of matrix *A*. If we normalize this vector by the total unmatched population, then

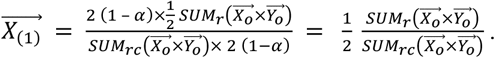

Similarly, it can be shown that

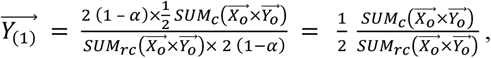

where the operation *SUM*_*c*_(*A*) sums the elements down each column of matrix *A*.

Notice that the sum of all the elements of 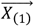 and 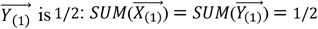.

Using the same calculations, it can be shown that after matching round 2, the total ratio of unmatched individuals is

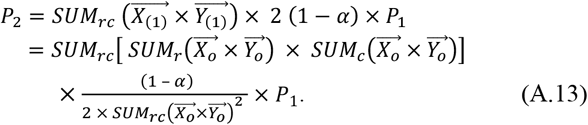

However, because

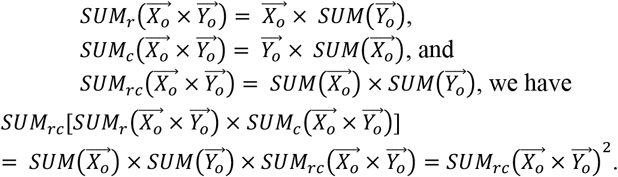

Substituting back into A.13:

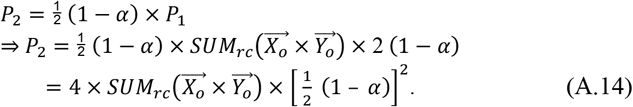

It follows that the normalized population vector

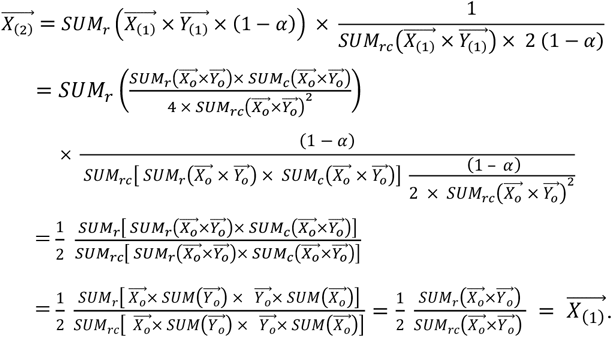

Similarly, it can be shown that 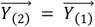,

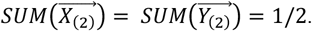

The additional matched individuals produced by matching round 2 are then

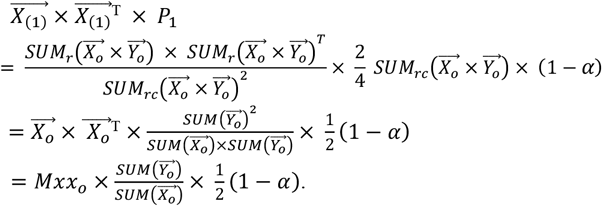

Similarly, using the same derivation, it can be shown that

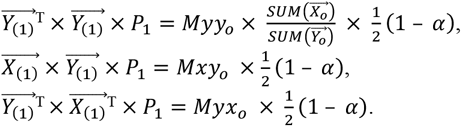

The accumulated matched individuals after matching round 2 are then

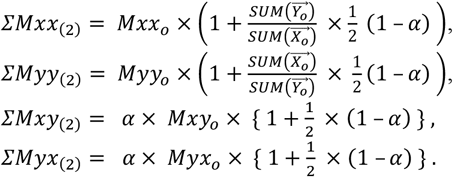

When we continue using the same derivations, it can be shown that after the *n*^*th*^ matching round:

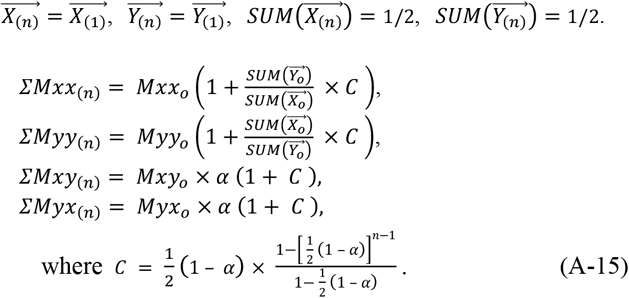

The proportion of unmatched individuals after the *n*^*th*^ matching round is

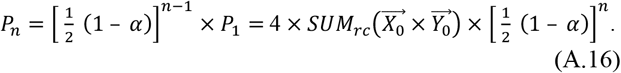

We can reconstitute Matrix *ΣM*_(*n*)_

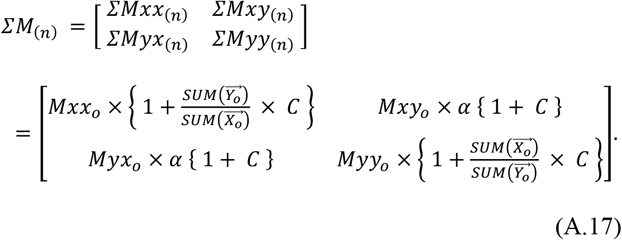

The elements in matrix *ΣM*_(*n*)_ specify the probability ratios and compositions of all the matched individuals in the ecoscape after the *n*^*th*^ matching round in a generation. Notice that summing all the elements of *ΣM*_(*n*)_ and adding the result to the ratio of unmatched individuals return a probability of one:

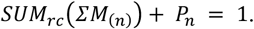

The unit offspring produced from the matched parents may be expressed mathematically as *ΣM*_(*n*_) ↦ *f*(*U*), where the symbol ↦ *f*(*U*) denotes element-wise multiplication of the elements of *ΣM*_(*n*_) (which specify the probability ratios of different parental matches) by the offspring proportions in each cell of *U* to produce different types of offspring. The different types of offspring are then added up to produce an output vector 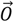 that shows the total unit offspring ratios (not normalized) produced from the generation:

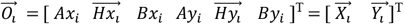, where *i* is the generation number.

Now, we can summarize the equations for each generation as follows:

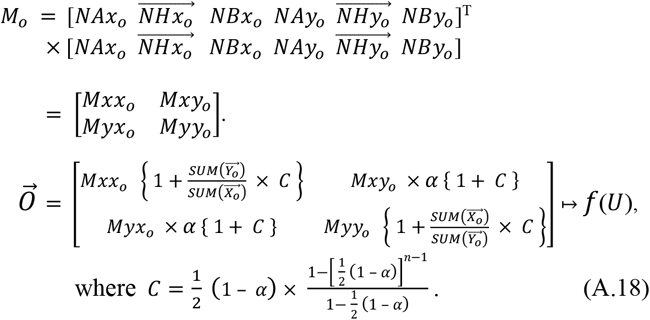

Finally, the difference equations for our system with viable hybrids are

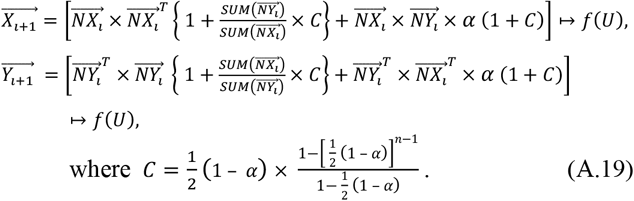

For the special case when 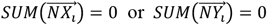, i.e., when the *X* or the *Y* allele is absent:

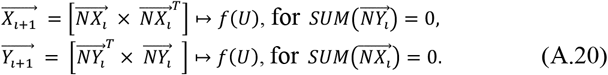

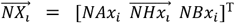 and 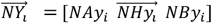 specify the normalized offspring ratios at the start of generation *i* that have survived ecological selection to become eligible mating individuals. 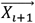 and 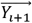 specify the unit offspring ratios (not normalized) produced from the matching rounds in generation *i*. Notice that the unmatched population from generation *i* is

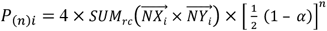, and that 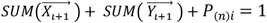.

Next, we can multiply the unit offspring ratios above by the number of offspring (*off*) that each matched individual is supposed to reproduce, and then multiply the result by the generation’s starting population 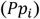 to obtain the offspring population for the next generation 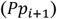. From these results, we can then calculate the normalized 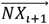 and 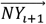 for the next generation.

In the case that *off* is sufficiently high so that the number of offspring produced in each generation is able to completely saturate the carrying capacities of all the niches, *NAmax*, 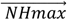, and *NBmax*, then the population ratios in niche *A*, niches *Hs*, and niche *B* going into the next generation are fixed. The normalized *NAx*, 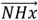, *NAy, NBx*, 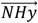, and *NBy* ratios going into the next generation can be calculated as follows:

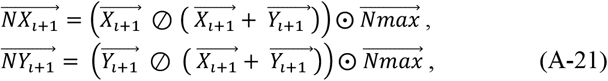

where 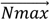 is a vector with elements that are the normalized ratios of all niche carrying capacities: 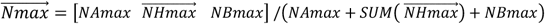.

The Hadamard operators ⨀ and ⊘ perform element-wise multiplication and division of two same-sized vectors or matrices.

This set of difference equations (A.19 – A.21) describes the dynamic behavior of a 2-niche, 2-mating-bias-allele sympatric system with viable hybrids in our mathematical model.

### 2-Mating-Bias-Allele Equivalent Model

Intuitively, the presence of viable hybrids should increase the unit offspring ratios that niche-*A* and niche-*B* groups can recover from their inter-niche matching encounters and reduce the impact of ecological selection (i.e., it effectively increases the value of *f*). To examine the effect of having viable hybrids on a hypothetical system without hybrid viability (Fig 5), Fig 22 shows how the matching probabilities and offspring proportions of *NA* and *NB* are affected by the presence of viable hybrids.

Let *Ax*, 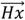, *Bx, Ay*, 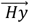, and *By* represent the normalized population ratios at the beginning of each mating generation. If 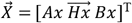 and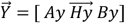, Fig 20 shows the matching probability matrix

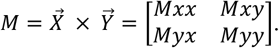

When viable hybrids are present, *Ax* acquires additional offspring from matches of 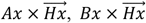, and 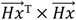, (shorthand 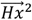) in submatrix *Mxx*; and 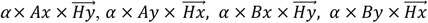, and 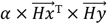 (shorthand 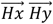) in submatrix *Mxy*.

In Fig 22, the multiplier 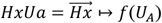 specifies the additional group *A* unit offspring that are produced because of the existence of 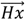 in the mating pool; and 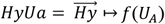 specifies the additional group *A* unit offspring that are produced because of the existence of 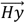. The multiplier 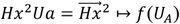 specifies the additional group *A* unit offspring that are produced because of the presence of matched mating pairs in 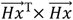 ; and similarly for 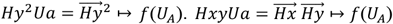 specifies the additional group *A* unit offspring that are produced because of the presence of matched mating pairs in 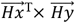. After *n* matching rounds in a generation, the additional unit offspring that are added to *Ax* due to matching with viable hybrids can be derived as

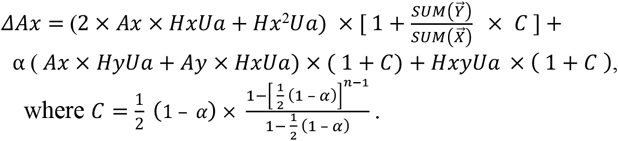

Similarly, multipliers 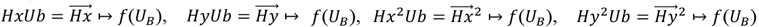, and 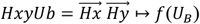 specify the additional group *B* unit offspring that are produced in a matching round because of the presence of viable hybrids in the mating pool.

The multipliers 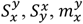, and 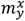 are defined such that

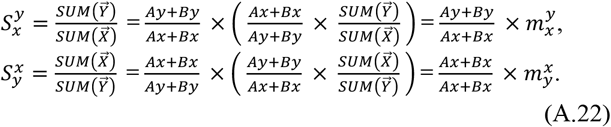

Then, 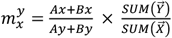 is a multiplier of the ratio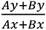, which is equivalent to the 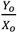 term used in the hypothetical model without viable hybrids that only has groups *NAx, NBx, NAy*, and *NBy* (A.8). Consequently, assuming that enough offspring are produced to saturate all niches in the ecoscape:

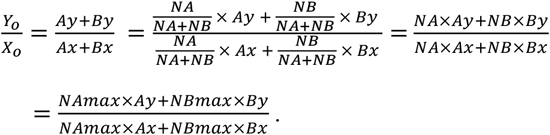

Similarly, we obtain

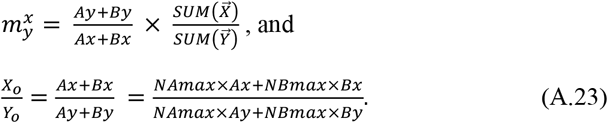

The following equations can be derived to calculate how the presence of viable hybrids in a mating generation changes the ratios of *NAx, NAy, NBx*, and *NBy* in a hypothetical system that does not have viable hybrids:

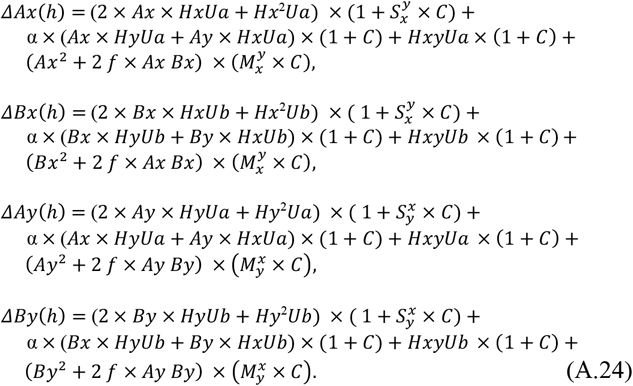

Fig 22 illustrates how viable hybrids impact a hypothetical system devoid of viable hybrids and how they interact with all the relevant multipliers.

### 3-Mating-Bias-Allele Model

The following details the mathematical algorithms used to calculate the normalized population ratios at the beginning of a mating generation *i* and the resultant normalized offspring ratios at the end of the mating generation, in a 2-niche, 3-mating-bias-allele sympatric system.

Let there be two ecological niches *A* and *B*. 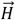 is a vector containing all possible hybrid ecotypes that can be produced by mating between individuals in niche *A* and niche *B. NA, NB*, and 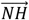 denote the normalized population ratios of ecotypes in their respective niches, so that 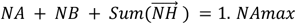 is the normalized maximum population ratio that niche *A* can support (i.e., its carrying capacity), and *NBmax* is similarly defined for niche *B*, and 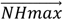 for niches in 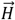, so that 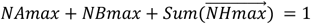. Next, we specify three mating-bias alleles, *X, Y*, and *Z*, with their corresponding mating-bias values *α, β*, and *γ* as shown in Fig 23. Let *Ax* be the ratio of *X* alleles in niche *A, Ay* be the ratio of *Y* alleles in niche *A*, and *Az* be the ratio of *Z* alleles in niche *A*, so that *Ax* + *Ay* + *Az* = 1. The mating-bias alleles in niche *B* are similarly defined so that *Bx* + *By* + *Bz* = 1. The same relationship holds for the mating-bias alleles in niches 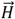, i.e., for 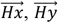 and 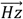. Then, *NAx* = *NA*× *Ax* is the normalized ratio of *X* mating-bias allele in niche *A*. If all the other mating-bias alleles in their respective niches are similarly defined, then

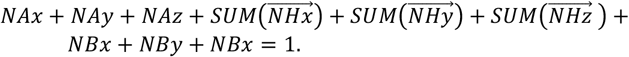

In a mating generation, the population goes through *n* matching rounds. The index *m* specifies the population’s current matching round. At the beginning of a mating generation, *m* = 0, and the normalized offspring population ratios from the previous generation become the normalized population ratios of the current generation, so that

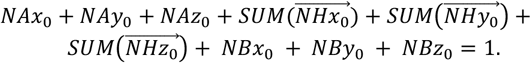

Let the vectors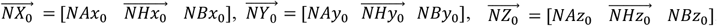, then 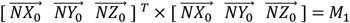, the matching probability matrix for the first matching round.

The elements of *M*_1_ represent the matched population ratios after the first matching round. *M*_1_ is composed of the following submatrices:

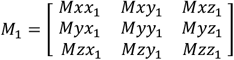, where 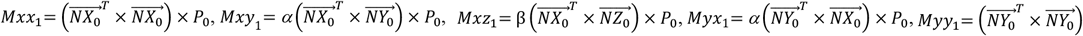 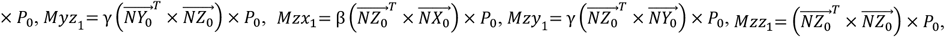 and

*P*_0_ = 1 for the 1^st^ matching round.

The ratio of total unmatched individuals after matching round *m* = 1 is

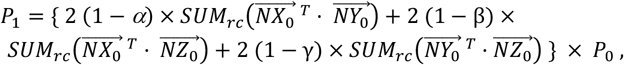

where the operation *SUM*_*rc*_ (*A*) sums all elements of matrix *A*.

The normalized ratios of unmatched individuals going into matching round 2 are then

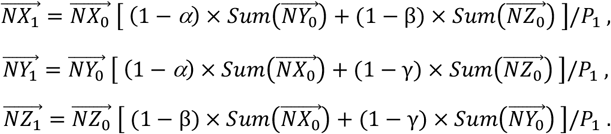

The proportional ratios of matched individuals in the 2^nd^ matching round can then be expressed as

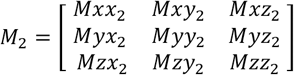, where

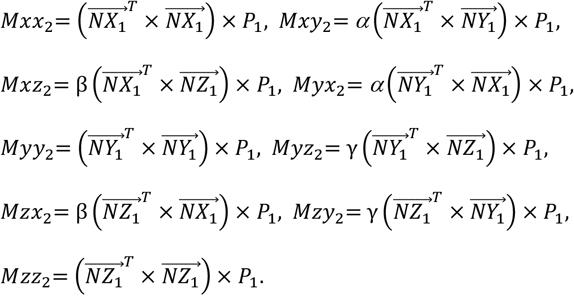

Generalizing from the above derivations, we can show that the total ratio of unmatched individuals after the *m*^*th*^ matching round is

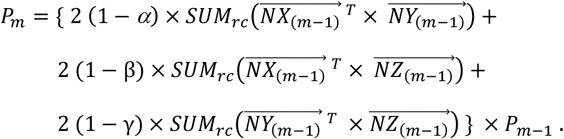

The total accumulated population ratios of matched individuals can be expressed by the matrix:

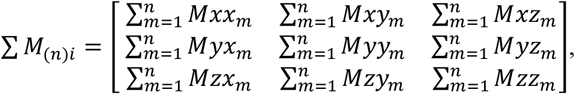

where

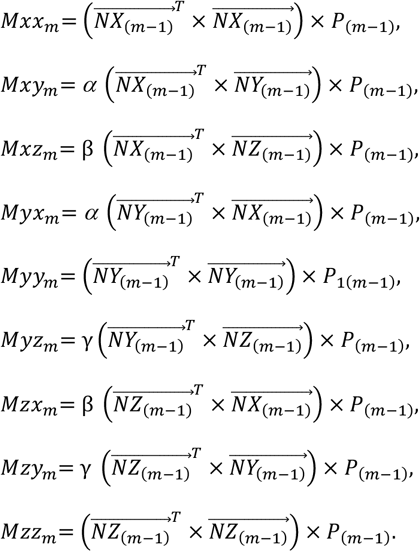

Let 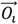 be a vector that shows the total unit offspring ratios (not normalized) produced from generation *i*:

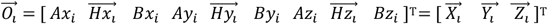, then

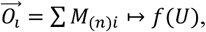

where the symbol ↦ *f*(*U*) denotes element-wise multiplication of the elements of *ΣM*_(*n*)*i*_ (which specify the probability ratios of different parental matches) with the offspring proportions in each cell of *U*, the unit offspring matrix for the matrix *ΣM*_(*n*)*i*_, to produce different types of offspring. The different offspring types are then added up to produce an output vector 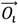 that shows the total unit offspring produced from the generation.

Next, we normalize the offspring ratios in 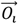 to obtain the normalized starting population ratios for the next mating generation. Assuming that the number of offspring produced in each generation is able to completely saturate the carrying capacities of all the niches, *NAmax*, 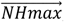, and *NBmax*, then the ratios of populations in niche *A*, niches *Hs*, and niche *B* going into the next generation are fixed. The normalized *NAx*, 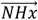, *NBx, NAy*, 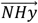, *NBy, NAz*, 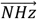, *NBz* groups going into E109. doi:10.1086/667586. PMID:22976018. the next generation can be calculated as follows:

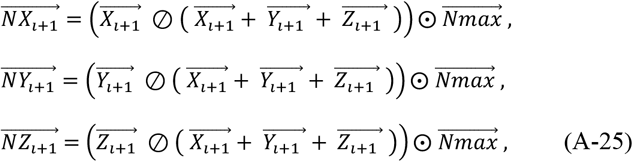

where 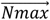 is a vector with elements that are the normalized ratios of all the niche carrying capacities:

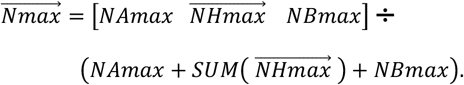

The Hadamard operators ⨀ and ⊘ perform element-wise multiplication and division of two same-sized vectors or matrices.

## Acknowledgments

We would like to express our appreciation to Editage for their English language editing and pre-submission peer review services. We thank Penny Brucker for copyediting and proofreading the manuscript and Durgesh Kushawaha for verifying the mathematical derivations. We are also grateful to Ashley Carter and Zakia Sultana for reviewing the manuscript and providing valuable feedback.

## Notes

### Competing Interest Statement

The authors have declared no competing interest.

### Summary of Updates

This updated version of the manuscript includes additional results and analyses concerning multi-locus mating traits in the models. The new findings are presented in Results Section X, titled 'Mating-Bias Alleles with Multiple Gene Loci.' Analyses of these findings are presented in Discussion Section 6 under the heading 'The effect of having multi-locus mating bias alleles.'

